# Informing NMR experiments with molecular dynamics simulations to characterize the dominant activated state of the KcsA ion channel

**DOI:** 10.1101/2020.12.14.422800

**Authors:** Sergio Pérez-Conesa, Eric G. Keeler, Dongyu Zhang, Lucie Delemotte, Ann E. McDermott

## Abstract

As the first potassium channel with a X-ray structure determined, and given its homology to eukaryotic channels, the pH-gated prokaryotic channel KcsA has been extensively studied. Nevertheless, questions related in particular to the allosteric coupling between its gates remain open. The many currently available X-ray crystallography structures appear to correspond to various stages of activation and inactivation, offering insights into the molecular basis of these mechanisms. Since these studies have required mutations, complexation with antibodies, and substitution of detergents in place of lipids, examining the channel under more native conditions is desirable. Solid-state NMR (SSNMR) can be used to study the wild-type protein under activating conditions (low pH), at room temperature, and in bacteriomimetic liposomes. In this work, we sought to structurally assign the activated state present in SSNMR experiments. We used a combination of molecular dynamics (MD) simulations, chemical shift prediction algorithms, and Bayesian inference techniques to determine which of the most plausible X-ray structures resolved to date best represents the activated state captured in SSNMR. We first identified specific nuclei with simulated NMR chemical shifts that differed significantly when comparing partially open vs. fully open ensembles from MD simulations. The simulated NMR chemical shifts for those specific nuclei were then compared to experimental ones, revealing that the simulation of the partially open state was in good agreement with the SSNMR data. Nuclei that discriminate effectively between partially and fully open states belong to residues spread over the sequence and provide a molecular level description of the conformational change.

## I. INTRODUCTION

Transmembrane allostery is at the heart of many signalling events in human biochemistry.^1^ KcsA has proved to be an excellent model system for studying this intriguing phenomenon, and has also served as a prototype for ion channel function.^2^ Activation coupled inactivation, a hallmark functional feature of essentially all ion channels, has been studied in KcsA by numerous biophysical and biochemical tools^3–11^. The literature in aggregate has resulted in a dominant hypothesis in which manipulation of an intracellular gate results in opening to a metastable activated state that lives for some milliseconds, during which time it directly triggers a slow allosteric inactivation involving loss of ions in the selectivity filter^3,5,12^. The details of the opening and coupled inactivation processes remain unclear and of great interest. Identifying the stable intermediates of these processes is central to clarifying the molecular basis for inactivation coupling in channels.

Solid-state nuclear magnetic resonance (SSNMR) studies of KcsA have provided unique insights regarding dynamics and thermodynamics of full length wild-type channels in hydrated lipid bilayers. These studies have the advantage of being carried out in near-native conditions, without use of antibodies, and with freely varied ligand concentrations ([K^+^] and [H^+^]). In our NMR studies, we have stabilized three states, which we have referred to as deactivated (stabilized at neutral pH and high [K^+^]), activated (stabilized at low pH and high [K^+^]) and inactivated (stabilized at low [K^+^]), with a naming convention selected because of the functional consequences of varying pH and [K^+^] (Fig. 1.a). An important prerequisite for interpretation of the NMR data is identifying (for each of these experimental conditions) the atomic arrangement that best represents the majority species in the sample. Identification of the deactivated and inactivated states is largely straightforward given the sample conditions used and the stability of these states: both X-ray diffraction (XRD) and solid-state NMR data suggest that the samples are homogeneous and robustly reproducible. Here the deactivated state corresponds to a channel with a closed activation gate (also called hydrophobic bundle crossing or inner helix bundle or inner gate) and a conductive selectivity filter fully loaded with potassium ions (PDB ID 1K4C)^13–17^. The inactivated state corresponds to a channel with a wide open inner gate and a pinched selectivity filter depleted of ions (PDB ID 5VKE, 5VKH, 3F7V, 3F7Y, 3F5W)^4,9,18,19^ (Fig. 1.a).

**FIG. 1.**
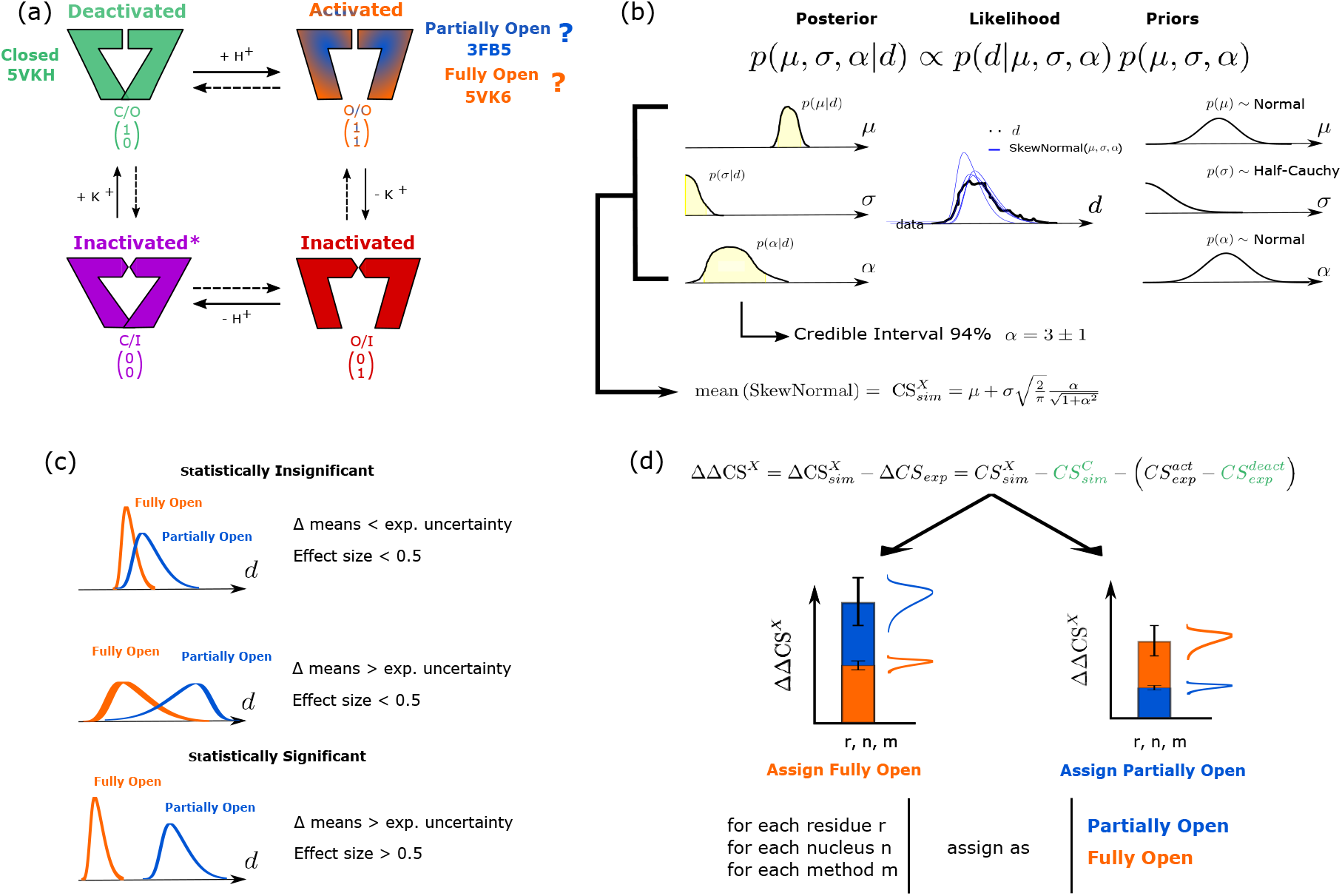
Summary scheme of the various computational steps carried out in this work. (a) Illustration of the gating cycle of KcsA and the corresponding X-ray crystallography structures with the nomenclature used for the different states. (b) Bayes law (top equation) and its terms: the posterior distribution (*p*(*μ, σ, α*| *d*)) is proportional to the product of the likelihood (*p*(*d| μ, σ, α*)) and the priors (*p*(*μ, σ, α*)) where *μ, σ, α* are the position, scale, and skewness parameters of a skew-normal distribution and *d* is the chemical shift data for a particular nucleus of the protein. The posterior probability of the parameters of a skew-normal distribution representative of the CS calculated from the MD simulation ensemble is proportional to the product of the likelihood of the data (predicted chemical shifts) multiplied by the prior distribution of the parameters (chosen here as uninformative). The definition of the mean of this skew-normal distribution is reported as the 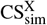 with its 94% credible interval. (c) Criteria imposed to consider a given residue’s CS as statistically significant. This is illustrated using two idealized CS-likelihoods obtained from the posteriors inferred for the two possible activated states, showing how their overlap and location affects statistical significance. This was done by assessing the difference in means between the two states 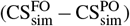 and the effect size of this difference 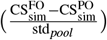 where 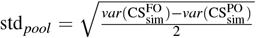 and *var* is the variance function. (d) Assignment of nuclei indicating compatibility of the activated state NMR sample with the PO or FO MD simulation ensembles. The method we use involves calculating the CS difference between the Closed (C) and the Partially Open / Fully Open (PO/FO) states (labeled X) measured using SSNMR (ΔCS_exp_) and predicted from the MD simulations ensemble 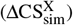 for the nuclei determined in (c). If the difference between the ΔCS measured experimentally and using the simulation ensemble (ΔΔCS^X^) is smaller for the PO than the FO state, this nucleus supports the conclusions that the experimentally observed activated state is the PO state seen in crystal structures. The reverse is true if the difference is smaller for the FO state.

The activated state, the only state that conducts ions, on the other hand, presents a particular challenge. Indeed, the seminal crystallography work by Cuello et al. showed that the development of a wide open activation gate was coupled to the appearance an inactive selectivity filter, and many other studies showed that the selectivity filter tended to collapse when the gate was held open, an observation compatible with the metastable short-lived nature of the activated state.^3,6,18^ Although the activated state has proven difficult to stabilize, X-ray crystallography studies involving a number of mutations and antibody binding succeeded in stabilizing a family of configurations with a conductive selectivity filter and the activation gate assuming a range of opening degrees, suggesting an almost continuous opening process^18^. A mutant channel (where all but one of the proton sensor charges were neutralized) was captured together with a conductive selectivity filter/Partially Open (PO) pore, with degrees of opening varying between 14 Å and 16 Å cross bundle distances at the narrowest site in the pore (measured as the distance between T112C_*α*_ of opposing subunits) (PDB IDs 3FB5 3FB6)^18^. Recently a non-inactivating mutant E71A structure was engineered with cysteines at positions 28 and 118 forming disulfide bonds to capture the channel in a Fully Open (FO) state, revealing a wider pore, of 23 Å cross bundle distance at the inner gate (PDB ID 5VK6). In this structure, interestingly, the gate was as wide open as in the inactivated state structure^4^.

These results raise a question about the degree of opening in the low pH near-native condition SSNMR activated samples. *De novo* structure determination of such a system from NMR restraints has not yet been reported. In this work, we have used an alternative approach to compare the compatibility of configurational ensembles obtained using MD simulations initiated from XRD structures with the NMR spectra. Chemical shifts (CS) have been shown to be very sensitive to biopolymer conformation, and in some cases contain sufficient information to correctly identify 3-dimensional structures.^20–26^ The use of chemical shift prediction tools has also been extended to restrain MD simulations in order to determine ensemble configurations compatible with the experiment^27^ and even to account for averaging of the experimental chemical shifts due to internal dynamics.^28–32^ Therefore we leverage CS information to identify the most likely structure for the NMR activated state. To do so, we ran MD simulations based on the two most likely representative XRD structures of the activated state (Partially Open PO, 3FB5 and Fully Open FO, 5VK6) and the deactivated state (Closed C, 5VKH) and predicted the ensemble of CS sampled by the trajectories of the NMR-assigned nuclei in the different residues. We then calculated relative chemical shifts (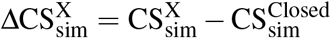 where X represents the FO or PO state), which are expected to be less affected by systematic errors than absolute chemical shifts. Using Bayesian inference, we determined signals that distinguished the PO and FO ensembles. We then calculated the deviation of these 49 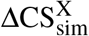 from the ΔCS_exp_ derived from the SSNMR experiments, leading us to conclude that the activated state probed in the SSNMR is closer to the PO than the FO state.

## II. METHODS

### A. Expression and purification of U-^13^C,^15^N wild-type KcsA

Uniformly ^13^C,^15^N labeled KcsA was overexpressed and purified as described previously from *Escherichia coli* PASK90/JM83 and PASK90/BL21(DE3) cells in M9 minimal media.^2,12,14^ Purified, uniformly ^13^C-^15^N labeled, KcsA in detergent dodecyl *β* -D-maltoside (DM) (Anatrace, Maumee, OH) was reconstituted into liposomes by reducing the detergent concentration through dialysis; the liposomes consisted of 3:1 1,2-dioleoyl-sn-glycero-3-phosphoethanolamine (DOPE): 1,2-di-(9Z-octadecenoyl)-sn-glycero-3-phospho-(10-rac-glycerol) (DOPG) (Avanti Polar Lipids, Alabaster, AL) at pH 4.0 and 50 mM KCl with a lipid to protein weight ratio of 1:1. The total ionic strength was kept at 100 mM by compensation with NaCl.

### B. NMR Spectroscopy

25 (or 9) mg of hydrated proteoliposome sample was centrifuged into a Bruker 3.2 (or 1.9) mm rotor after 3-5 freeze-thaw cycles. SSNMR experiments were performed on either a Bruker AVANCE II or NEO spectrometer operating at 21.1 T (ω_0,H_/2π = 900 MHz) or 17.6 T (ω_0,H_/2π = 750 MHz). Experiments were collected with either a 3.2 mm HCN E-free probe (900 MHz) or a 1.9 mm HCN probe (750 MHz) (Bruker BioSpin, Billerica, MA), as indicated.

Experiments at 900MHz were collected with a spinning frequency, *ω_r_*/2π, of 16666 ± 10Hz and a set point temperature of 267 ± 1K. The sample temperature is estimated to be 290 ± 5 K due to sample heating from MAS and RF pulses.^33^ Standard π/2 pulse lengths of 2.55 μs, 4μs, and 5.1 μs were used for the ^1^H, ^13^C, and ^15^N channels, respectively, corresponding to ω_1_ /2π = 98 kHz (1H), 62.5 kHz (^13^C), and 49kHz (^15^N). Two-dimensional (2D) ^13^C-^13^C, 2D ^13^C-^15^N (NcaCX, NcoCX), and three-dimensional (3D) ^13^C-^13^C-^15^N (NCOCX, NCACX) correlation experiments were used to determine the chemical shift assignments.^34–36^ The ^13^C-^13^C mixing was performed using dipolar assisted rotational resonance (DARR) mixing of 50 ms.^37^ The heteronuclear polarization transfer used SPECtrally Induced Filtering In Combination with Cross Polarization (SPECIFIC-CP).^38^ SPECIFIC-CP standard RF conditions of *ω*_1,C_ = 1.5 *ω_r_* and ω_1,N_ = 2.5 *ω_r_* for NCA, and ω_1,C_ = 2.5*ω_r_* and ω_1,N_ = 1.5*ω_r_* for NCO and contact times between 4.5 and 5.0 ms were used for SPECIFIC-CP experiments. ^1^H decoupling of ω_1,H_/2π > 90 kHz during acquisition was performed using SPINAL-64 decoupling.^39^ More detailed experimental parameters for the 3D experiments are given in Table S1.

Experiments at 750 MHz were collected with spinning frequencies of 16000 ± 10 or 33333 ± 10Hz and a set point temperature of 230 (ω_r_/2π = 33333Hz) or 240 (ω_r_/2π = 16000Hz) ± 1 K. The sample temperature is estimated to be 290 ± 5 K due to sample heating from MAS and RF pulses.^33^ Standard π/2 pulse lengths of 2.55 μs, 3.3 μs, and 4.3 μs were used for the ^1^H, ^13^C, and ^15^N channels, respectively, corresponding to ω_1_ /2π = 98 kHz (^1^H), 76kHz(^13^C), and 58 kHz (^15^N). The same 3D experiments for chemical shift assignments were performed at 750 MHz. The ^13^C-^13^C mixing at 750 MHz was performed using COmbined 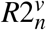-Driven (CORD) mixing of 50ms.^40^ SPECIFIC-CP RF conditions of ω_1,C_ = 0.5ω_r_ and ω_1,N_ = 1.5ω_r_ for NCA and NCO at *ω_r_*/2π = 33333 Hz and ω_1,C_ = 1.5ω_r_ and ω_1,N_ = 2.5ω_r_ for NCA, and ω_1,C_ = 0.5ω_r_ and ω_1,N_ = 1.5ω_r_ for NCO at *ω_r_*/2π = 16000Hz. SPECIFIC-CP contact times of between 4.5 and 5.0 ms were used. A 2D ^13^C-^15^N ZF-TEDOR experiment^41^ was collected with 1.5 ms mixing and *ω_r_*/2π = 16000 Hz for identification of the proline chemical shifts. ^1^H decoupling of ω_1,H_/2π > 90 kHz during acquisition was performed using swept-frequency two-pulse phase-modulated (SW_*f*_-TPPM) decoupling.^42^ More detailed experimental parameters for the 3D experiments are given in Table S2.

All data were processed in Topspin (Bruker BioSpin) or NMRPipe.^43^ The ^13^C and ^15^N chemical shifts were referenced via the substitution method to the methylene peak of adamantane at 40.48ppm (0.5% DSS in D_2_O scale), and to the peak of ^15^NH4Cl to 39.3ppm (at 298K, NH_3_ (liquid) scale), respectively.^44–47^ Peak picking and assignments were made in CcpNmr.^48^ All the spin systems were identified first in each 3D spectrum and then the residue type and their secondary structure were predicted using PLUQin.^49^ Backbone resonance assignments were determined based on correlations in the 3D NCACX, NCOCX, and CANCO spectra, beginning from unique spin systems (for which the residue type was identified using PLUQin). Sidechain residue assignments were determined mainly based on 3D NCACX, NCOCX, and 2D ^13^C-^13^C DARR experiments. The following tolerance thresholds were applied for the agreement of chemical shifts when comparing the various spectra of the activated state for assignments: 0.2 ppm for ^13^C, and 0.5 ppm for ^15^N. Monte Carlo/simulated annealing (MC/SA) algorithms in the software packages NSGA-II and MC/SA were used to facilitate assignments.^50^

### C. Molecular dynamics simulations

Coordinates for PO, FO, and C KcsA structures (PDB IDs: 3FB5, 5VK6, 5VKH)^4,18^ were prepared using CHARMM-GUI^51^. The constructs included residues 26-121. For 3FB5, Rosetta^52^ was used to add missing residues 114-121 in *α*-helical conformation. The N- and C-termini were capped by methylation (NME) and acetylation (ACE). We note that the pH sensor was not included in the simulations to keep consistent the chain-lengths across states. Using PYMOL^53^ the following mutations were reverted: for the FO state (5VK6), A28C, E71A, L90C, R117Q, E118C, E120Q, R121Q; for the PO state (3FB5), L90C, R117Q, E120Q, R121Q and for the closed state (5VKH), Y82A, L90C, F103A. The sidechain of residue E71 was oriented to form a hydrogenbond with D80 as done in previous works in the literature^3^. None of the residues that were mutated in the activated states (3FB5 and 5VK6) were later found to be important to discriminate between the PO and FO states. For all states, residues E71, E118, and E120 were protonated since H-NMR spectra showed these residues to be protonated in the activated state.^15,54^ We note that the simulated protein contains no histidines due to the truncation of the sequence. The selectivity filter was initially fully loaded with K^+^ in agreement with the X-ray diffraction and crystallization waters bound to the channel were included. The protein was embedded in a DOPE:DOPG (3:1) membrane bilayer containing 150 lipids and surrounded by a neutralizing solution of 50 mM NaCl and 50 mM KCl at a ratio of 75 TIP3P^55^ rigid water-molecules for each lipid molecule. All states were modelled including the co-purifying DOPG lipids, as identified in the X-ray diffraction structures.

The systems were equilibrated using the CHARMM36 force field^56^ in an extended CHARMM-GUI protocol. 1μs-long simulations were performed using GROMACS2019^57^. The Parrinello-Rahman barostat^58^ (*P* = 1bar τ = 5ps) and Bussi thermostat^59^ (*T* = 290K τ = 1ps) were employed to mimic experimental conditions. The parts of the simulation after a water molecule enters the selectivity filter were excluded since such is a symptom of inactivation (see Fig S1)^6^. Additionally, the initial parts of the simulation where the inner-gate distance was unstable were also excluded from analysis (see Fig S2). Additional technical details of the simulation can be obtained from the simulation files and trajectories freely available at https://github.com/delemottelab/Informing_NMR_experiments_w_MD and https://osf.io/6h2z5/.

### D. Statistical Inference of Chemical Shifts from MD simulations

N, C, C_*α*_, and C_*β*_ CS of the simulation snapshots were calculated using the SPARTA+^60^ and SHIFTX2^61^ chemical shift prediction tools, with their default options. For SHIFTX2, this implies turning on using the homology-based algorithm SHIFTY+ if possible. Nevertheless, due to a lack of homology with the sequences in RefDB^62^, this selection had no effect. Although quantum mechanical NMR chemical shift prediction tools are more accurate than empirical methods, using them would constitute an overly ambitious understaking due to the size of the protein and the number of MD snapshots in the configurational ensemble. Only nuclei experimentally assigned for both states were considered. Since KcsA is a homotetramer, we were able to collect four times the data per residue under the assumption that the trajectories of the monomers were independent. Nevertheless, since there are contact surfaces between the monomers that can affect the CS prediction, the prediction software was given the full tetramer and the data obtained for the four equivalent subunits was concatenated *a posteriori*. The CS-prediction error for each MD simulation snapshot that is associated to the accuracy of the prediction tool was not modelled. The overall agreement of the conclusions drawn for both methods highlights the general robustness of the methods.

The statistical distributions of the CS were modelled using Bayesian inference with the python libraries PYMC3^63^ and Arviz^64^. In line with this work, there have been previous studies that used Bayesian inference methods to analyze heterogeneous ensembles of CS derived from chemical shift predictions or calculations.^65–67^ Bayesian inference assumes that the probability distribution of a random variable has a given functional form and enables to infer the probability of the parameters of the distribution given the data sampled (*d*). A schematic description of the procedure is found in Fig. 1b. The CS of each particular nucleus, state, residue, and prediction method was considered as an independent variable and was therefore modelled separately in a non-hierarchical framework. For every variable, two thousand data points were selected randomly out of all the time frames obtained from each MD trajectory.

The likelihood function chosen was a skew-normal distribution as implemented by PMC3:

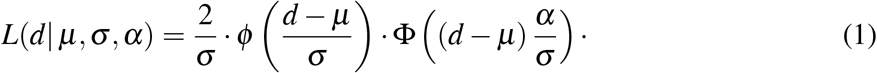

where *d* is the data sampled for the 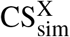 of a particular nucleus, and for a particular residue in that particular state X. *ϕ* and Φ are the probability distribution function and cumulative distribution function of the Gaussian distribution:

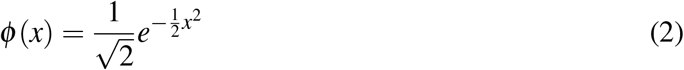

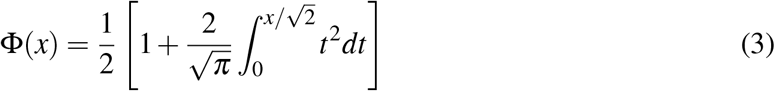

The skew-normal function has three parameters: the location (*μ*), the scale parameter (*σ*), and skewness parameter (*α*). In order to conduct Bayesian inference, previous assumptions of the parameters are included by assigning probability distribution functions, *priors*, to them. For the *μ* parameters, normal distributions are assigned:

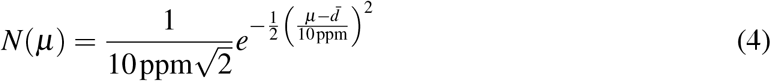

Where 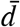 is the average value of the sampled 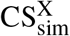 data (d) and 10 ppm the scale of the normal distribution. For σ, the priors assigned are half-Cauchy distributions, assigned with a beta parameter of 10 ppm:

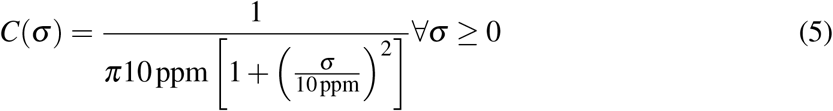

For *α*, normal distributions are also assigned:

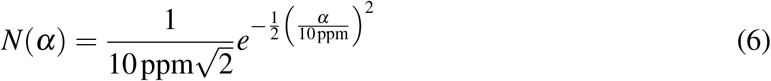

The posterior distribution was sampled using the NUTS^68^ algorithm with 4 chains. It is important to remark that the mean of a skew-normal distribution is not the location parameter μ but rather 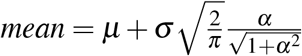. This mean value of the distribution is what we will consider the chemical shift 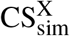 of state X predicted by our MD simulation, or in other words the simulated position of the peak in the NMR spectrum. This corresponds to a physical assumption of rapid population weighted average of the chemical shifts sampled in the protein dynamics. In similar fashion, the distribution variance was calculated as 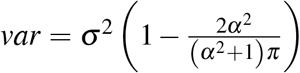 Posterior predictive checks were carried out after the inference to test the adequacy of the model (Fig S3). The main features of the data distributions are generally in good agreement with the model. Even in the case where the distribution is bimodal, the envelope of the skew-normal function covers the bimodal distribution (Fig S3). Additional tests and inference information can be found in the SI and https://github.com/delemottelab/Informing_NMR_experiments_w_MD.

All quantities obtained with Bayesian inference are not assigned a single value, but rather a probability distribution. As a consequence, for simplicity, the distributions of these random variables can be summarized by the interval that contains 94% of its probability. This interval is known as the 94% credible interval (CI) and is written as its center plus or minus the distance to the bounds of the interval. A graphical representation can be found in Fig 1b.

### E. Selecting nuclei whose CS differ statistically between states

The next step was to determine which nuclei were statistically different in the PO and FO trajectories. For this, the difference between the simulated chemical shifts for both states modeled as described above was calculated 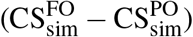, as well as the effect size of this difference 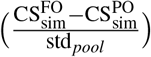 where 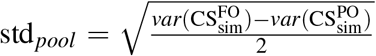 and *var* is the variance function. The first expression measures if there is a difference between the CS predicted for both states and the second if this difference is large compared to the variance of the distributions. The importance and graphical interpretation of the difference in means and effect size can be seen in Figure 1c.

A CS variable for a given nucleus in a given residue generated using a given chemical shift prediction method is considered to be have discrimination power if the center of the difference in means credible interval is greater in absolute value than the experimental tolerance (0.2 ppm for C, C_*α*_, C_*β*_ and 0.5 ppm for N) and the center of the effect size credible interval is larger than 0.5 in absolute value. An effect size greater than 0.5 is considered medium as a general rule in statistics69. This procedure is repeated independently for each studied nucleus, for each chemical shift prediction method, and for each residue.

### F. Comparing CS differences from MD simulations and NMR

Site specific CS for the PO and FO states were contrasted with experimentally measured CS to assess which state is present in the experiment. A schematic representation of the procedure can be found in Fig. 1d. 94% credible intervals were calculated for both the FO and PO states (designated “X” since it is to be determined which is correct), using the expressions 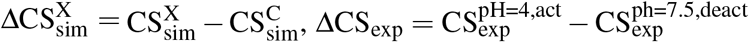, and 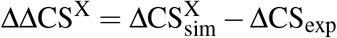. The state, X, for which ΔΔCS^X^ is lower in magnitude has a predicted 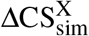 closer to the experimentally measured ΔCS_exp_ and we conclude it to be likely that this state is dominant in the experiment at that nucleus. This procedure was repeated for all nuclei in all residues for which experimental shifts were available, using both CS prediction software packages. It is interesting to note than point measures of error such as root mean squared error (RMSE) give differences between the states that are in the range of the experimental uncertainty (Table S3). These differences only start to become substantial if the statistical filtering information and the closed state is used as reference.

## III. RESULTS

### A. NMR CS Assignments of the activated state sample

Chemical shift assignments of the activated state (pH 4.0, 50 mM [K^+^]) of KcsA were determined using three-dimensional spectra (NCACX, NCOCX, and CANCO) acquired at two different external magnetic fields (900 and 750 MHz), see Table S4. Representative strip plots of the backbone walk for the assignments of residues T74 to G79 using these spectra are given in Figure S5. The completeness of the backbone and sidechain assignments, along with the experimental connectivities of each resonance in each experiment, are presented in a schematic representation in Figure S6. A number of the assigned resonances, including some that are used in the analysis presented here, are well resolved in various 2D spectra, as illustrated in Figures 2 and S7. Chemical shifts for the deactivated state at pH 7.5, 50mM [K^+^] (previously reported,^15^ Table S5) were re-referenced using the protocol described in this paper so as to be comparable to the shifts determined here for the activated state (pH 4.0, 50 mM [K^+^]). Distributions of the chemical shift differences (comparing activated and deactivated states) indicate that the chemical shift referencing is consistent for the two datasets (Figure S8). In fact, the site-specific chemical shifts were found to be similar for the activated and deactivated states, with few notable exceptions, suggesting that the two states adopt similar structural folds, as has been indicated from X-ray crystallography studies (Table S6). Resonances were considered to be experimental markers if the chemical shift difference (activated vs. deactivated, 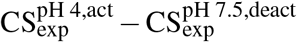) was greater than the experimental tolerance (0.2 ppm for ^13^C and 0.5 ppm for ^15^N). If this difference does not exceed this threshold the assigned resonance was considered to be an experimental spectator.

**FIG. 2.**
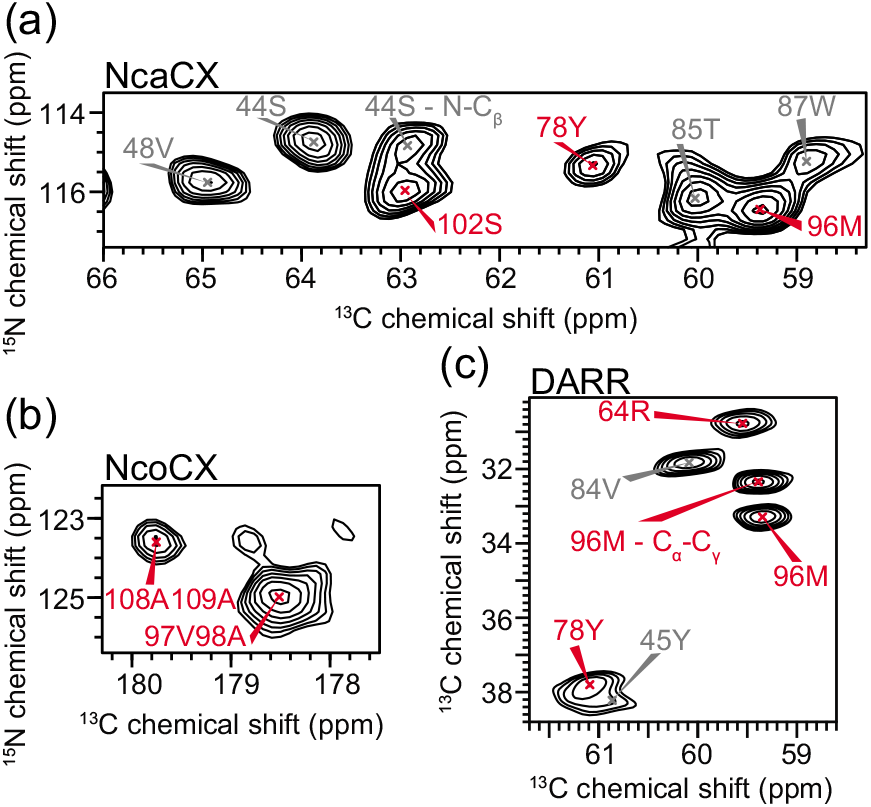
Representative experimental NMR spectra of KcsA in the activated state (3:1 DOPE:DOPG, 50 mM KCl, pH 4.0). (a) NCA region of 2D ^15^N-^13^C NcaCX spectrum with assigned peaks shown. All peaks in the NcaCX spectrum are 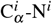 unless noted otherwise. (b) NCO region of 2D ^15^N-^13^C NcoCX spectrum with assigned peaks shown. All peaks in the NcoCX spectrum are C^*i*–1^-N^*i*^ unless noted otherwise. (c) C_*α*_ -C_*β*_ region of 2D ^13^C-^13^C DARR spectrum with assigned peaks labeled. All peaks in the DARR spectrum are 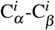 unless noted otherwise. Peaks used to distinguish between fully open vs partially open (identified by statistical inference analysis of the CS predictions of MD snapshots) are labeled in red, and those identified (by the same methods) as spectators are labeled in gray.

### B. Sampling the deactivated and activated KcsA conformational ensemble using MD simulations

In order to run MD simulations to characterize the activated state, the most plausible XRD structures were selected as starting coordinates. There is a rich set of such structures for KcsA, which mostly differ in their inner gate opening (measured as the distance between T112 C_*α*_ from opposing subunits): 3FB5 (14 Å), 3FB6 (16 Å), 5VK6 (22 Å) and 3F5W (31 Å)^4,18^. In this work, the 3FB5 structure was used instead of 3FB6 because of its higher resolution and therefore fewer missing side chains. Nevertheless, the two structures are very similar and the MD simulation initiated from 3FB5 coordinates explores openings compatible with 3FB6 (Fig. S3). 3F5W was found to have an unstable inner-gate that moved towards openings compatible with 5VK6 and therefore was not included in this study (Fig. S4). Therefore, the initial configurations were prepared based on the 3FB5 PO structure, and from the 5VK6 FO structure. We also ran simulations of the C state (conductive selectivity filter, closed inner gate), to compare the activated state to, preparing a system based on the 5VKH high resolution crystal structure.

The simulation conditions were chosen to match the experimental conditions as closely as possible. The lipid and solution composition in the computational system were identical to the ones in the NMR experiment: 3:1 DOPE:DOPG in a symmetrical bilayer and a concentration of 50 mM of NaCl and 50 mM of KCl in the solution. The simulation temperature and pressure were set to 290 K and 1 atm, respectively, thus close to experimental conditions. To mimic the pH of the experiments, acidic residues E71, E118, and E120 were protonated and the protonation states were kept fixed due to limitations of classical MD simulations. The main difference between the simulated and experimental constructs is the truncation of the sequence to keep residues 26 to 121 due to the lack of experimental coordinates to model the intracellular domains. In the three states, the models were overall stable over the hundreds of nanoseconds timescale, as characterized by the RMSD (Fig. S9).

The opening of the inner gate was found to be fairly stable for both the C and FO states (Fig S3). In contrast, the PO state inner gate experienced some initial instability, followed by asymmetrization, and finally a drift towards closing. For this reason data collected between 400 ns and 1000 ns, corresponding to a stable opening, was used in the analysis. This observation is consistent with observations from previous MD simulation studies^3^. The conductive selectivity filter of all three states was fully loaded with K^+^ at the start of the simulation and was found to rapidly evolve to the WKK0KW occupancy, where the symbols W, K and 0 stand for occupacy of the site by a water molecule, a K^+^ ion, or nothing, respectively (Fig. S1). In the FO state simulation, a water molecule entered the selectivity at t=400 ns. Water penetration in the selectivity filter has been linked to inactivation, particularly in this inactivation-prone wide-gate state^6^. Therefore the rest of the simulation data was discarded. In contrast, consistent with previous observations^3^, for the C state, no instabilities in the structure or the selectivity filter occupancy were noticed such that the full trajectory length was considered for the analysis. The simulation data for the different states that were used for CS analysis is summarized in figure S1.

### C. Chemical Shift Prediction from the MD Results

We used SPARTA+^60^ and SHIFTX2,^61^ two well established tools, to predict the chemical shifts expected for the ensemble of structures observed in the MD trajectories. Predictions were made based on numerous frames from the MD trajectory, and subsequently the average and the distribution in predicted shifts was analyzed, following a procedure analogous to Robustelli et al.^28^ The two tools (SPARTA+ and SHIFTX2) produced chemical shifts in qualitative agreement; however, the predicted chemical shifts were not identical for the two tools (Figure S10). The results from both methods were thus analyzed separately to ensure robustness of the conclusion. We note that direct comparison of the CS predicted directly from the XRD structures does not give a clear picture of which state is closer to experiment (Fig. S11).

The distribution of the predicted CS based on the MD simulation trajectory is strikingly non-Gaussian as can be seen in Figures S12 – S19. This fact introduces difficulty in assessing the statistical significance of differences, and a need for a robust method to assess the statistical significance of changes in CS between simulated states. Therefore, we chose to model the CS distributions of the simulation using Bayesian inference (Fig. 1b).

The goal of Bayesian inference is to numerically calculate the *posterior* distribution of the parameters of the distribution of CS shifts (chosen here as a skew-normal distribution, see Methods, Figure 1b), p(μ, σ, α |d), given our data (which is here the CS predictions from MD snapshots). The posterior distribution is thus a measure of the uncertainty of the CS shifts distribution parameters considering the data we have collected. Note that we assign uncertainty to our knowledge of the parameters and not to the data directly, contrary to a classical statistics approach in which parameters that we assume to be certain are estimated, and then the probability to obtain the data sampled calculated by a p-value given the obtain parameters.

To calculate the posterior probability we must choose the terms of the right hand side of Bayes law. The *likelihood* term or probability of the data given the parameters, p(*d|μ, σ, α*), is specified by choosing the distribution with which we model our data presented in the central column of Figure 1b. After the inference has been carried out, the posterior distribution of the parameters can be sampled to obtain the ensemble of skew-normal distributions inferred. This ensemble can be compared to the data to see the adequateness of the model. This procedure is known as *posterior predictive check* (Figure S3). As Figures S12 – S19 shows, our data is adequately characterized by skew-normal distributions. A skew-normal distribution can be defined by three parameters: the location (μ), the scale parameter (σ), and skewness parameters (*α*).

Finally the posterior distributions are estimated numerically using the NUTS68 algorithm. This algorithm also calculates the normalization factor *p*(*d*) or the probability of the data. Once these posterior distributions are known, we can calculate the probability distributions of the mean of the skew-normal distribution 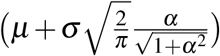, which we ascribe to the NMR-peak position and call 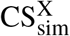, where X is the state simulated. 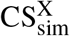, like all quantities obtained with Bayesian inference, is thus not assigned a single number but rather a probability distribution. As a consequence, for simplicity, the distributions of these random variables can be summarized by the interval that contains 94% of its probability. This interval is known as the credible interval (CI) and is written as its center plus or minus the distance to the bounds of the interval. This is analogous to the confidence intervals in classical statistics.

### D. Activated State Characterization Through Statistical Analysis

We approached the question of what structure best represents the activated samples studied by NMR spectroscopy by asking two questions. First, we asked which chemical shifts are predicted to distinguish the FO and the PO states. Then we asked whether the NMR shifts corresponding to the activated state suggest better agreement with the FO or the PO configuration.

Many of the predicted 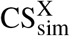 shifts do not change sufficiently between the PO and FO states to offer any statistical power of discrimination (Figure S20, S21). We thus developed the following protocol to identify nuclei capable of discriminating between the two states.

The difference between the simulated chemical shifts of both states 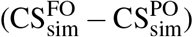 and the effect size of this difference 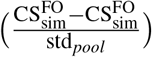, where 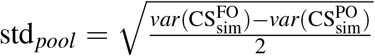 and *var* is the variance function, were calculated. The first expression estimated whether there was a difference between the CS predicted for both states and the second if this difference was large compared to the variance of the CS distributions. A graphical interpretation of this procedure can be found in Figure 1c. Due to the effect size condition for statistical significance, this methodology relies on having a chemical shift distribution per nucleus and can only be performed having an ensemble of protein configurations obtained from statistical simulation. In addition, this filtering relies solely on simulated data.

An example of effect size and difference in means for C_*α*_ and the chemical shift prediction method SPARTA+ is presented in Figure 3. In this figure, we present the difference in means (left graph) and the effect size (right graph) for amino acids whose experimental C_*α*_ CS have been assigned in both activated and deactivated states.

**FIG. 3.**
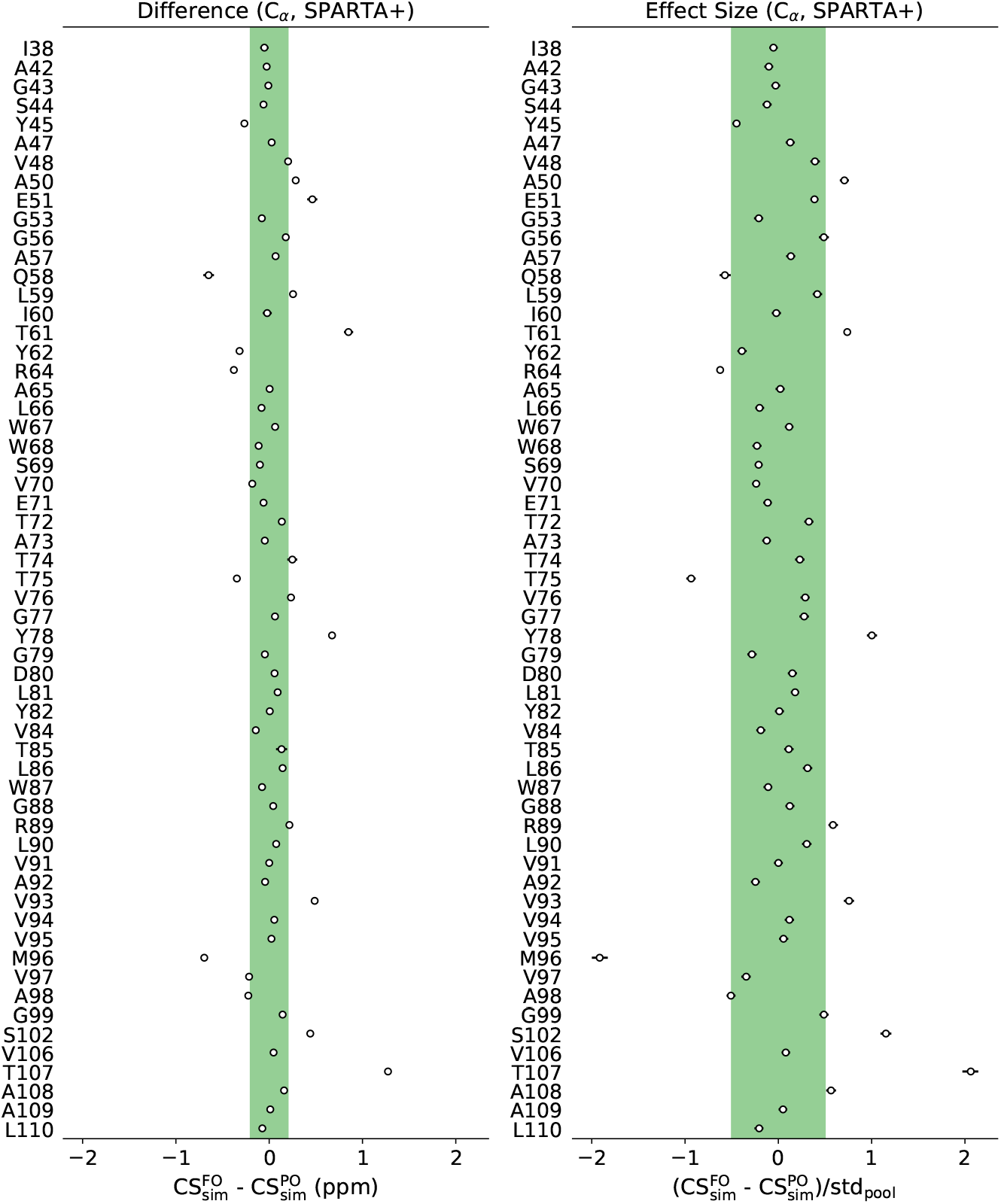
Statistical filtering of simulated CS. Circles represent the center of the 94% credible interval of the variable distribution and horizontal bars represent the full range of the corresponding credible interval (which are not visible in cases where the interval is smaller than or comparable t the symbol size). The vertical axis represents the residues that are part of the experimental dataset in both pH conditions. We show here CS differences for the C_*α*_ nuclei using the SPARTA+ chemical shift prediction method. Left: credible interval of 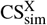 difference in means between the Fully Open (FO) and Partially Open (PO) states. If a credible interval center is outside the experimental tolerance range (green shade), the difference is considered significant. Right: credible interval of the effect size (difference in means scaled by the pooled standard deviation) between the FO and PO states. If the credible interval center is greater than 0.5 in absolute value (green shade), the effect size is considered medium-large^69^. A residue must have a significant difference in means and a large effect size to be considered as having discrimination power. The same procedure was carried out for the rest of the nuclei and chemical shift prediction methods (Figs. S23,S24)

A CS variable of a given nucleus in a given residue and generated by a given chemical shift prediction method is considered as having discrimination power if the center of the CS difference CI is greater in absolute value than the experimental tolerance (0.2 ppm for C, C_*α*_, C*_β_* and 0.5 ppm for N) and the center of the effect size CI is larger than 0.5 in absolute value. This procedure is repeated independently for each studied nucleus, for each chemical shift prediction method, and for each residue (Figures S23, S24). 14 C_*α*_ nuclei, 11 C*_β_*, 11 C, and 13 N nuclei were found to fit these criteria, yielding a total of 49 nuclei with potential to distinguish the two open states.

As mentioned above, the strategy developed in this work involved analysis of open-vs-closed or activated-vs-deactivated difference shifts rather than absolute chemical shifts. Specifically, the experimental chemical shift changes between the activated and deactivated states (ΔCS_exp_) was compared to the predicted chemical shifts changes between the FO/PO vs the C simulation MD ensembles 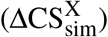. An overview of the method can be found in Figure 1d. Such a strategy of calculating difference chemical shifts is likely to be less prone to a number of the errors that make calculations of absolute shifts very challenging. For example, CS referencing errors will not be present in the difference shift calculation. Also, consistent effects that occur in the experimental data but are not captured by the prediction algorithm (e.g. effects of the lipid environment, or tertiary interactions) will presumably cancel and produce less systematic error in the calculation of difference shifts if the states (open vs closed) are relatively similar. Indeed, the use of ΔΔCS^X^ 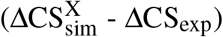 has allowed us to make stronger predictions than absolute CS (CS_sim_ – CS_exp_) would have (Figure S25). Additionally, the error distribution of the ΔΔCS^X^ is narrower and more centered at zero than the analogous distribution of absolute chemical shifts (Figure S26) implying that the use of ΔΔCS^X^ has smaller random and systematic errors and is thus is more suited for state prediction than absolute chemical shifts. Alternatively, correcting the center of these distributions (CS_sim_ – CS_exp_) by fitting to a Gaussian distribution (Figures S27-S28) and then shifting the center of this Gaussian distribution to zero yields a small improvement over the absolute CS difference (Figure S29). This strategy of computing difference shifts may be suitable only to specific situations. Considering absolute chemical shift predictions appears to be preferable if the system undergoes major conformational transitions or if only one state of the system is accessible experimentally. The approach we propose should instead be considered when the structure of a reference state is available and changes in the system are relatively subtle, with many structural features preserved between the various conformational states of the system.

Figure 4a shows the ΔΔCS^X^ for the PO (blue bars) and FO states (orange bars) for the different nuclei and predicted using the two different prediction software tools. Missing bars correspond to an absence of experimental CS signal or to a statistical non-significance of the simulated value difference between states (Figure S22). The lower the ΔΔCS^X^, the closer the simulation prediction is to the experiment. Therefore if, for a given nucleus, a given method, and a given residue, the bar corresponding to the PO state is lower than the bar corresponding to the FO state, we consider the 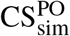 of the PO state to agree better with the 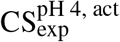 of the activated state. If both signals are below the green shaded area, which indicates the experimental tolerance, a most likely state is not assigned. Most of the 49 distinguishing nuclei identified in this study indicate a better agreement with the PO state (61%) than with the FO state (25%), suggesting that the activated samples prepared for NMR analysis more closely agree with a partial opening of the activation gate. If instead we consider which state the majority of the nuclei of a residue indicate, these percentages change to 59%, 9% and 32% for PO, FO, and undetermined respectively.

**FIG. 4.**
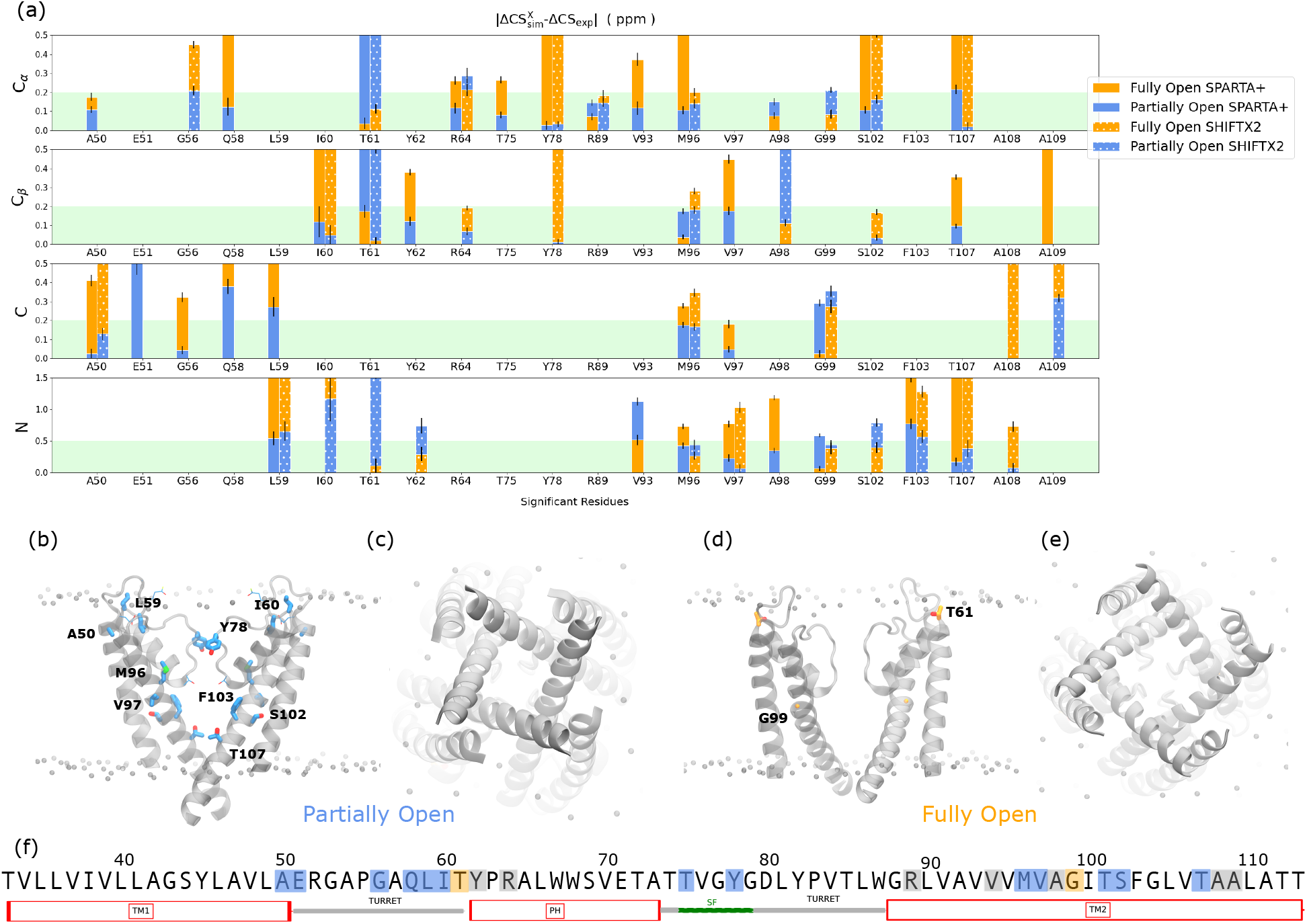
(a) Centers of 94% credible intervals of the difference in relative chemical shifts between experiment and simulation in absolute value. The limits of the credible interval are shown as error bars. Both experiment and simulation use as reference the closed state chemical shifts: 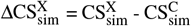 and 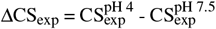, where X represents the FO or PO sate. The residues found to be capable of discriminating between PO and FO using our statistical criteria are represented on the x-axis for the different nuclei and chemical shift prediction methods. The agreement between MD simulations and the NMR experiments is higher the closer the value is to zero. The PO state (blue) has in general a better agreement with experiment than FO state (orange). The green shade depicts the typical experimental uncertainty. If two methods produce statistically identical CS or the signal is missing in the spectrum, the bar is not represented. (b) Side view of the molecular structure of the PO state. Only two subunits are shown. Residues identified using one chemical shift prediction method are shown with a thin “licorice representation”. Residues identified using two chemical shift prediction methods are shown with a thick “licorice representation”. (c) Molecular structure of the PO state viewed from the intracellular side. (d) and (e) show the same representations as (b) and (c) but for the FO state. (f) KcsA sequence, truncated to region of the protein used in the MD simulations. Blue highlights represent residues whose experimental chemical shift agrees with the simulated PO state, orange highlights those in agreement with the FO state and grey highlights inconclusive residues.

The nuclei displaying better agreement with the PO state (highlighted in blue in Figure 4b,c) are located in the turret region linking TM1 and the pore helix (A50, E51, G56, Q58, L59, I60), in the selectivity filter (T75, Y78), and in the transmembrane part of TM2 (M96, V97, S102, F103, T106 and T112). Additional nuclei spread all over the sequence were inconclusive (grey highlights in Figure 4c – Y62, R64, R88, V93, A98, A108 and A109). Only two residues displayed better agreement with the FO state (orange highlights in Figure 4b,c – T61 and G99).

Within the approach of analyzing the difference in predicted and experimental chemical shifts, one can imagine a number of logical ways to compare the various structural states or MD ensembles (FO, PO, and C) to the two experimental states (activated and deactivated). We developed an *ad hoc* classification system summarized briefly in Figure 5 and in more detail in Figure S30. Individual nuclei are first grouped in 5 classes based on the predicted shifts for the three MD ensembles, as follows: (**A**) a very well populated group for which the three structural states are predicted to give rise to indistinguishable shifts (akin to experimental spectators), (**B**) a well populated group for which the FO state is predicted to be distinct from the C and the PO states (which are themselves not distinguishable from each other); (**C**) a poorly populated group for which, curiously, PO is distinct from FO and C (which are themselves not distinguishable from each other); (**D**) an unpopulated hypothetical group for which the C state is distinct but PO and FO are indistinguishable 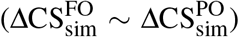, which has been eliminated by our statistical filtering analysis; and (**E**) a fairly well populated group for which all three computed states are expected to give rise to distinct chemical shifts. Each nucleus is further classified in terms of whether (**1**) the two experimental states have indistinguishable chemical shifts, or (**2**) if a change in the chemical shift is observed. Each nucleus is then identified as being category **B1**, **B2**,... up to **E2** (and in some cases is identified as belonging to two classes because the two chemical shift prediction tools that were used were not in agreement).

**FIG. 5.**
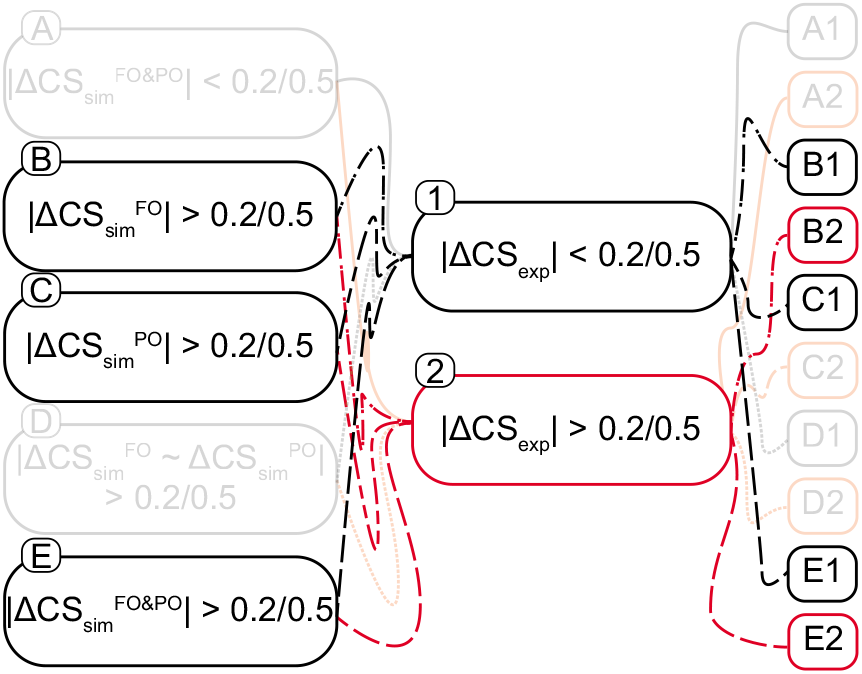
Flow chart demonstrating the combinations of the 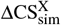 and ΔCS_exp_ that yield various classifications of the state markers identified in this study. A threshold of tolerance of 0.2 ppm for ^13^C and 0.5 ppm for ^15^N was used to determine if the CS change was significant. FO = fully open, PO = partially open, C = closed, 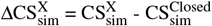, and 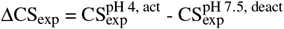.

Two classes of nuclei are of particular note in this study. A sparsely populated type, **E2**, where the predicted CS is different for each state and the experimental data for the nucleus also indicates a shift, has strong ability to address the questions posed in this work. (This class includes A50C, E51C, T107C_*α*_, T107N, A109C, and A109C_*β*_). These markers clearly show a better agreement with the conclusion that the sample is PO rather than FO (Figure 4a). Nuclei classified as **B1**, for which the FO state (but not the PO state) is predicted to show a deviation from C, and for which the experimental data obtained at both pH conditions are indistinguishable, are the most common (apart from the spectators **A**, which are not discussed here). Resonances that fall into the **B1** classification in aggregate offer strong support for the conclusion that the activated samples studied experimentally are closer to PO than to FO (A50C_*α*_, G56C_*α*_, G56C, Q58C_*α*_, L59N, I60C_*β*_, R64C_*β*_ (SPARTA+), T75C_*α*_, Y78C_*α*_, Y78C_*β*_, R89C_*α*_, M96C_*α*_, M96C_*β*_, M96C, M96N, V97C_*β*_, V97C, V97N, G99N, S102C_*α*_, and A108N).

Another group of CS are classified as **B2**, when the FO state is predicted to differ from the PO and C states, and the experimental data do show a shift (Q58C, A97Cβ, G99C, S102N, and A108C) and yet another as **E1** for which the predicted CS differs for each structural state while the data do not reflect a change in CS (L59C, I60N, T61C_*α*_ (SPARTA+), Y62C_*β*_, V93C_*α*_, A98C_*α*_ (SHIFTX2), and F103N). These two types of markers could be inconclusive although in some cases there is a much better match for one hypothesis than the other (typically the PO state being in better agreement).

A handful of nuclei are observed as class **C1** if the predicted PO shifts are different from the FO and C states while the experimental data show no shift. At first sight, these would appear to indicate that the sample is FO. The pattern is difficult to interpret mechanistically and the phenomenon is mainly observed in the turret region for which the connection to opening is unclear and where antibody binding in the crystal structure may have complicated the MD sampling (T61C_*α*_ (SHIFTX2), T61C_*β*_, T61N, Y62N, R64C_*α*_ (SHIFTX2), R64C_*β*_, V93N, A98C_*α*_ (SPARTA+), and G99N).

No resonance is classified into **C2**, where the PO shift is distinct from the other states and there is an observed experimental chemical shift change between the activated and deactivated states. This particular class would suggest that the resonances that are identified within it would favor the sample to be in the PO state. Notably, some additional classifications in this study are populated by no resonances, namely **D1** and **D2**, due to the initial conditions of the analysis that require a resonance be distinguishable between the FO and PO states.

### E. Molecular Basis for KcsA activation

The chemical shift markers identified by the SSNMR experiments reflect conformational gating of the corresponding residue during opening. These sites exhibit a variety of conformational differences in the MD simulations, including sidechain rotamer hops and backbone rearrangements (especially at the TM2 hinge) (Figure 6).

**FIG. 6.**
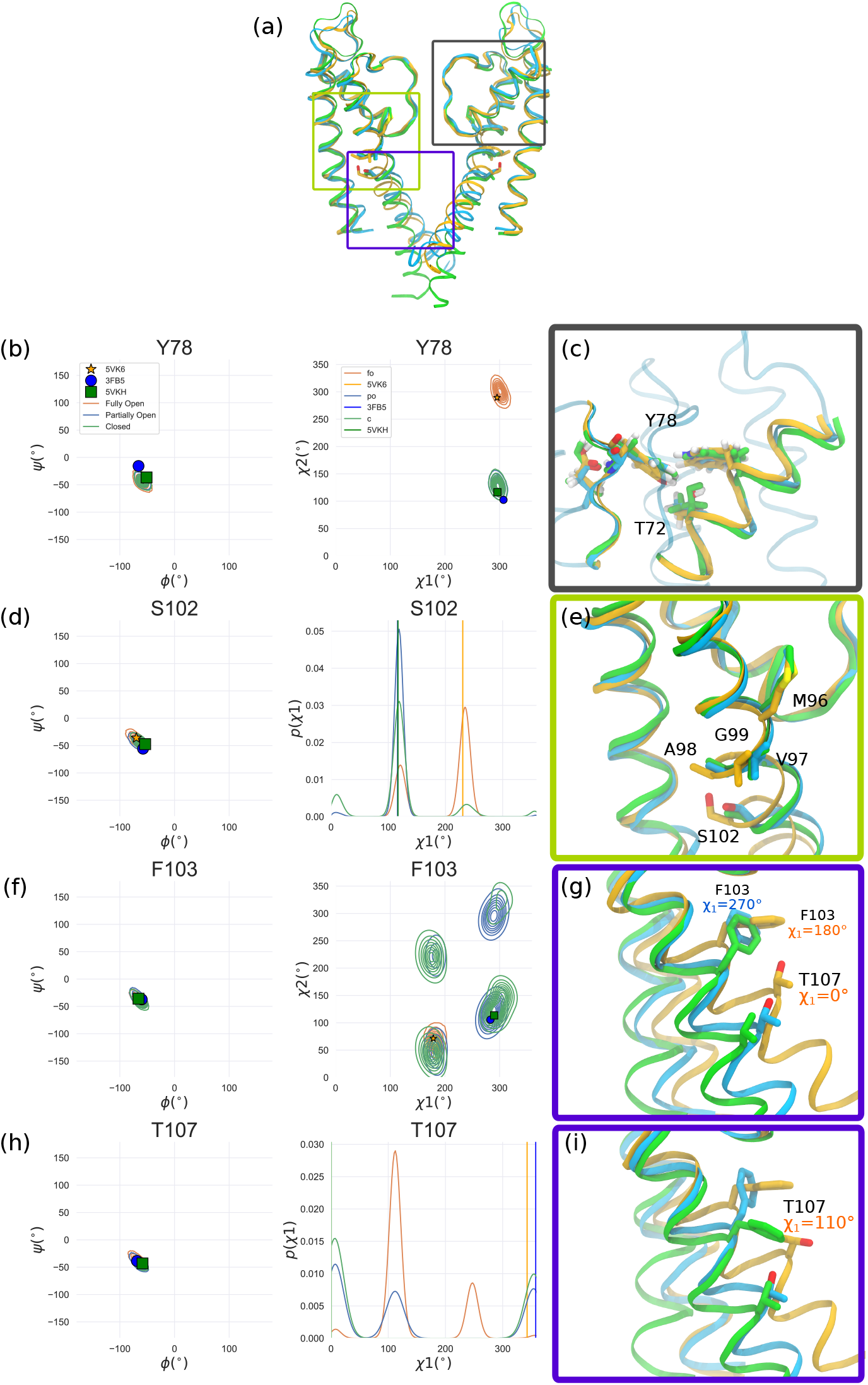
Structural comparison between C, PO, and FO states (a) Side view of 2 opposite subunits in the C (green ribbon), PO (blue) and FO states (orange). The selectivity filter region is highlighted by a grey box, the hinge region by a green box and the cavity facing helices by a purple box. (b) Dihedral (Ramachandran and *χ*_1_-*χ*_2_ plots) angle plots showing the MD simulation distributions (surface contours, coloring is the same as in panels (a) and (c) corresponding molecular models showing details of the environment around Y78 in the three states. Snapshots were aligned on the selectivity filter residues backbone (75-80) of the subunit Y78 belongs to. (d-e) Same as (c) for S102 (f-i) same as (c) for F103 and T107. The molecular views in the purple boxes (g,i) show the rearrangement of the rotameric state of T107 from the crystal structure to the most prevalent rotameric state observed in MD simulations, in the FO state.

Several of the residues that enabled the distinction between the PO and FO states (A50, A98, S102, T107, A108 and A109) also exhibit experimental chemical shift changes when comparing deactivated and activated datasets (Figure 5). S102 and T107, which undergo rotameric changes in their *χ*_1_ dihedral angles, in particular, offer particularly strong support to the conclusion of partial opening (Figure 6d,h).

Scrutinizing additional dihedral angle plots revealed other substantial rearrangements at residues identified to discriminate the PO and the FO states in the selectivity filter (Y78), in the hinge region (M96, V97, A98, and S102) and in the residues facing the central pore cavity (F103 and T107) (Figure 6). The hinge region residues displayed subtle changes in the alpha helical basin, as expected from the hinge motion that is key for gate opening described in numerous studies (Figure 6d-i).^3,4^ F103 has indeed been reported as a key player in transmitting the allosteric signal coupling the activation gate and the inactivation gate at the selectivity filter.^3^ In the C and PO states, several rotameric states are accessible to the *χ*_1_ angle of this residue, while in the FO state the residue is locked in the state found in the XRD structure avoiding exposure to the hydrated cavity. Several studies have found that its clash with T74 leads to SF pinching linked to inactivation^3,70^. Our work seems to indicate that in high [K^+^] and low pH conditions, locking of this residue in the rotameric state compatible with inactivation is not favored. T107 assumes a similar rotameric state in the C, PO and FO state XRD structures (Figure 6h). Intriguingly, our MD simulations show that another χ_1_ dihedral conformation is more prevalent in the FO state. This FO rotameric state appears unfavorable in the activated state condition SSNMR. While the molecular basis for this observation remains unclear, we note that a CS prediction based on a pure XRD structure diffraction (in the absence of MD simulations) would have missed this observation.

The only SF residue with discriminating power which shows an interesting rotameric change is Y78 (Figure 6b). Indeed the other SF residue, T75, is a weak marker, with the C_*α*_ nucleus CS pointing towards the PO state only when predicted by SPARTA+. Y78 is at the center of a hydrogen bonding hub which connects the selectivity filter and the pore helix of adjacent subunits, via hydrogen bonding to the side chains of T72 and W68 and the backbone carbonyl of E71 (Figure 6c). The rotameric state of Y78 *χ*_2_ differs between the XRD structures of the C and PO state, on one hand, and of the FO state, on the other (Figure 6b). The slight change in Y78 orientation in the FO state appears to lead to a motion of the pore helix. Interestingly no pore helix residues, including T72 and W68, showed a change in predicted CS between the PO and the FO states.

Only two residues indicated a better agreement with the FO state, T61 and G99. It would be intriguing if a portion of the channel were in a partially open like state and another portion were fully open, but this seems not likely to be quite correct. For example T61 is a more complex situation without a clear assignment to either state. A comparison of the Ramachandran distributions of T61 in both states reveals that both states explore a common conformer but the PO state samples an additional conformer (Fig. S31). This conformer, which is only present in the PO state simulations, is only explored by one of four subunits. If the chemical shifts produced from this subunit are removed then the difference in CS between PO and FO states becomes statistically insignificant (Fig. S32). We attribute this conformational state to a fluctuation in the dynamics of the flexible turret region which may not be representative of the entire configurational ensemble. This region was indeed in contact with the antibodies used to stabilized the sample in the XRD structure and numerous indications in the literature suggest dynamic plasticity for these residues.^18^ G99 is located in the center of the hinge region, surrounded by residues that exhibit good agreement with the PO state. The fact that its chemical shift agrees better with the FO state appears intriguing. Indeed, the dihedral angles explored by G99 in both states are fairly similar and correspond to the initial angles of the X-ray structure (Fig. S31). Why it is the only residue in better agreement with the FO state in this region remains to be explained.

## IV. CONCLUSIONS

The open conductive state of KcsA has remained elusive to structural studies due to its short lifetime. Mutagenesis and other engineering strategies have led to the determination of several XRD structures, but since they were obtained under somewhat artificial conditions, the physiologically relevant state remains of interest to be fully characterized. Spectroscopy studies have revealed that there is considerable conformational heterogeneity within the open states of KcsA of the degree of opening, making a static characterization not only challenging but also limited in its physiological relevance. SSNMR allows the study of a sample at near-physiological conditions, but interpreting the spectra in terms of structure is challenging. MD simulations can assist with this interpretation since the conditions of the MD simulation and the SSNMR experiment can be nearly equivalent, and the time-scales probed are compatible. This procedure rests on the use of chemical shift prediction software which have been refined over the years to yield robust predictions.

Here, we have designed a computational pipeline to characterize the activated state present in the SSNMR activated state sample. The two candidates from X-ray crystallography that we compared it to were a partially open PO (PDB ID 3FB5) and a fully open FO (PDB ID 5VK6) structure. While a direct comparison between the experimental and computationally predicted CS was noisy, implementing a Bayesian inference scheme in which nuclei were selected when they had the predictive power to statistically discriminate the PO and the FO states, and the use of difference chemical shifts (relative to the deactivated/closed state) for both the experimental and the predicted shifts, proved more successful. With this approach, we were able to conclude that the majority state in the SSNMR resembles the PO state.

Why does the system stall in the partially open state, instead of advancing to form the inactivated state as it does in electrophysiology experiments? Formation of the fully open state may linked to ion expulsion, and so may be prevented in these SSNMR experiments by the relatively high concentration of ambient potassium. In other words, perhaps the FO state is unstable in the presence of bound potassium ions. Indeed, under some conditions the FO state characterized by X-ray crystallography exhibited reduction in the ion occupancy in the selectivity filter.^18^ MD simulations of the opening also appear to indicate a slow process from partial opening to full opening, with an unknown inherent bottleneck, of a possibly entropic nature, or it is possible that the system does not fully open absent other coupled processes such as ion loss. Alternatively, perhaps the various differences between the sample conditions (protein concentration, symmetric vs. asymmetric lipids and pH, etc.) cause the system to stall in one case and advance to inactivation in the other.

The conclusion that the activated state studied by SSNMR is a partially open state seems compatible with a number of earlier studies or hypotheses. It has been speculated in prior studies that the state stabilized for NMR studies by cardiolipin lipids at low pH is partially open.^71^ It is also consistent with the fact that preparation of the FO state for X-ray crystallography experiments required a number of mutations and cross links, without which the partially open state resulted. The results collectively suggest a mechanism in which opening, once triggered by protonation on the pH gate, proceeds to an intermediate point which can be stabilized under appropriate experimental conditions such as high K^+^ concentration. The degree of opening is possibly a balancing act between repulsions in the protonated pH sensor that tend to push the system open, and conformational clashes in the hinge and base of the selectivity filter, which tend to keep the system closed.

This work demonstrates an approach to combining SSNMR and MD simulations to provide a molecular level description of a physiologically important allosteric process in a membrane protein.

## V. SUPPLEMENTARY MATERIALS

See Supplementary Materials for additional data figures and tables.

## VI. DEDICATION

We dedicate this paper to Professor Cynthia Jameson of UIC for her contributions to fundamental understanding of the NMR chemical shift, mapping the NMR Chemical Shift and coupling constants across the periodic table, and providing insights into intermolecular effects, and the importance of considering ensembles in calculating shifts. This dedication recognizes Cynthia’s many contributions to furthering equity and diversity in science, especially for women scientists.

## VII. ACKNOWLEDGEMENTS

E.G.K. was supported by a postdoctoral fellowship from the National Institutes of Health (NIH) (Grant No. GM135350). A.E.M. was supported by Grant No. GM088724 from the NIH and Grant No. 1412253 from the National Science Foundation. The NMR data were collected at the New York Structural Biology Center (NYSBC) with support from the Center on Macromolecular Dynamics by NMR Spectroscopy, a Biomedical Technology Research Resource (NIH Grant No. GM118302). The NYSBC is also supported by the Empire State Division of Science Technology and Innovation and the Office of Research Infrastructure Program (NIH Grant No. CO6RR015495). A.E.M. is a member of the NYSBC. S.P.C. was supported by a postdoctoral scholarship of the Gustafsson foundation. L. D. would like to thank the support of Science for Life Laboratory, the Göran Gustafsson Foundation, and the Swedish research council (Grant No. VR-2018-04905). Simulations were performed on computational resources provided by the Swedish National Infrastructure for Computing (SNIC) at the PDC Centre for High Performance Computing (PDC-HPC). The authors thank Zhiyu Sun for the synthesis of one of the KcsA samples used in this work. The authors thank Dr. Keith Fritzsching for helpful discussions.

## VIII. DATA AVAILABILITY

The data that support the findings of this study are openly available in https://github.com/delemottelab/Informing_NMR_experiments_w_MD and https://osf.io/6h2z5/ with the raw NMR data deposited in BMRbig (https://bmrbig.org) with under deposition ID bmr-big9.

