## Supplementary Material for "Informing NMR experiments with molecular dynamics simulations to characterize the dominant activated state of the KcsA ion channel"

(Dated: 24 February 2021)

[a)] These three authors contributed equally

[b)] Correspondance to:

### I. SUPPLEMENTARY FIGURES

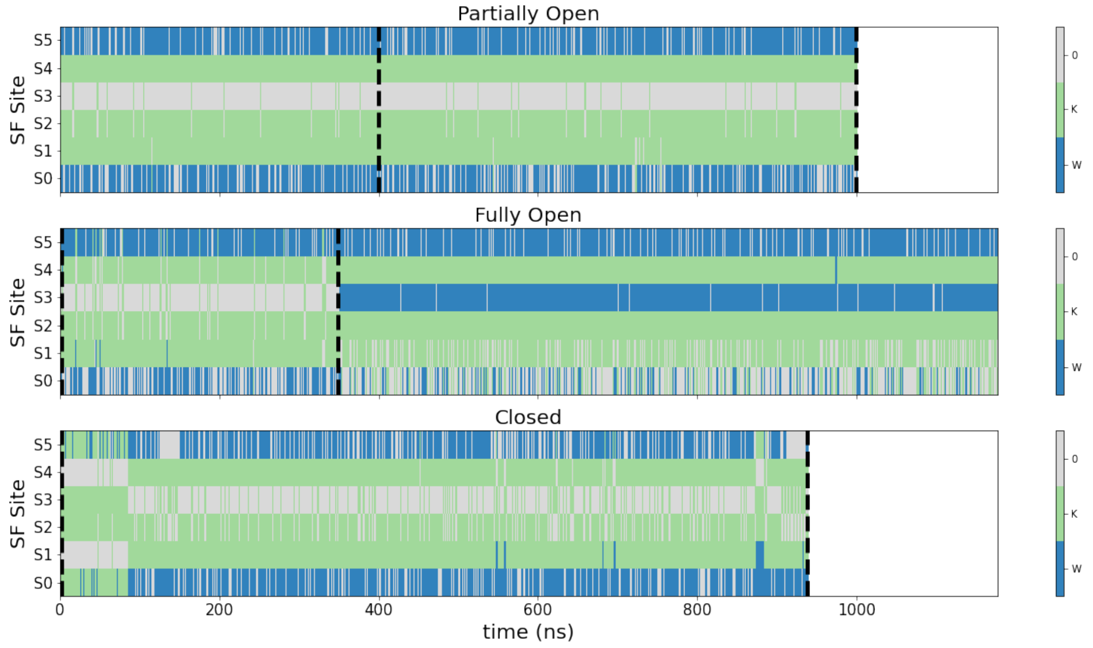


FIG. S1. Occupation of the canonical selectivity filter sites as a function of time in the MD simulations for the study states: Partially Open (top), Fully Open (middle) and Closed (bottom). For the Fully Open state the simulation data from the entry of a water molecule in the S3 site is discarded since this has been identified as a pre-inactivation sign.

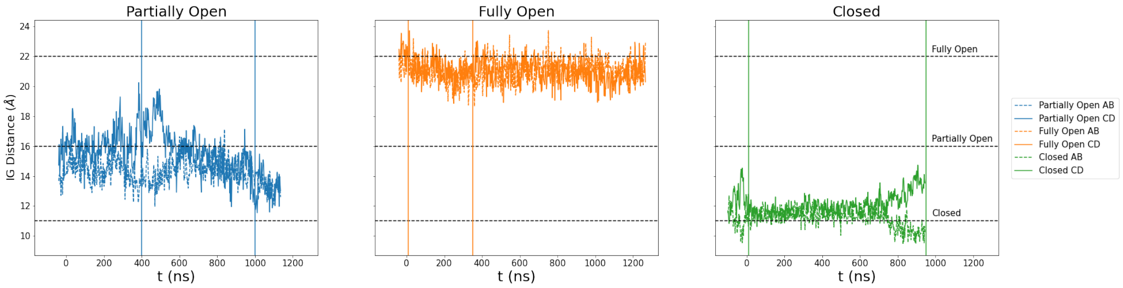


FIG. S2. Inner gate opening measured as the T112 $C_\alpha$ distance of opposing subunits ( subunits AB solid line or subunits CD dashed line). Vertical lines delimit the part of the simulation analyzed. In the case of the partially open state the part with a more stable inner gate is used. For the Fully Open state this choice is based on selectivity filter occupation see Fig. S1. The data is smoothed with 50 point rolling median. Negative values of time are the equilibration period.

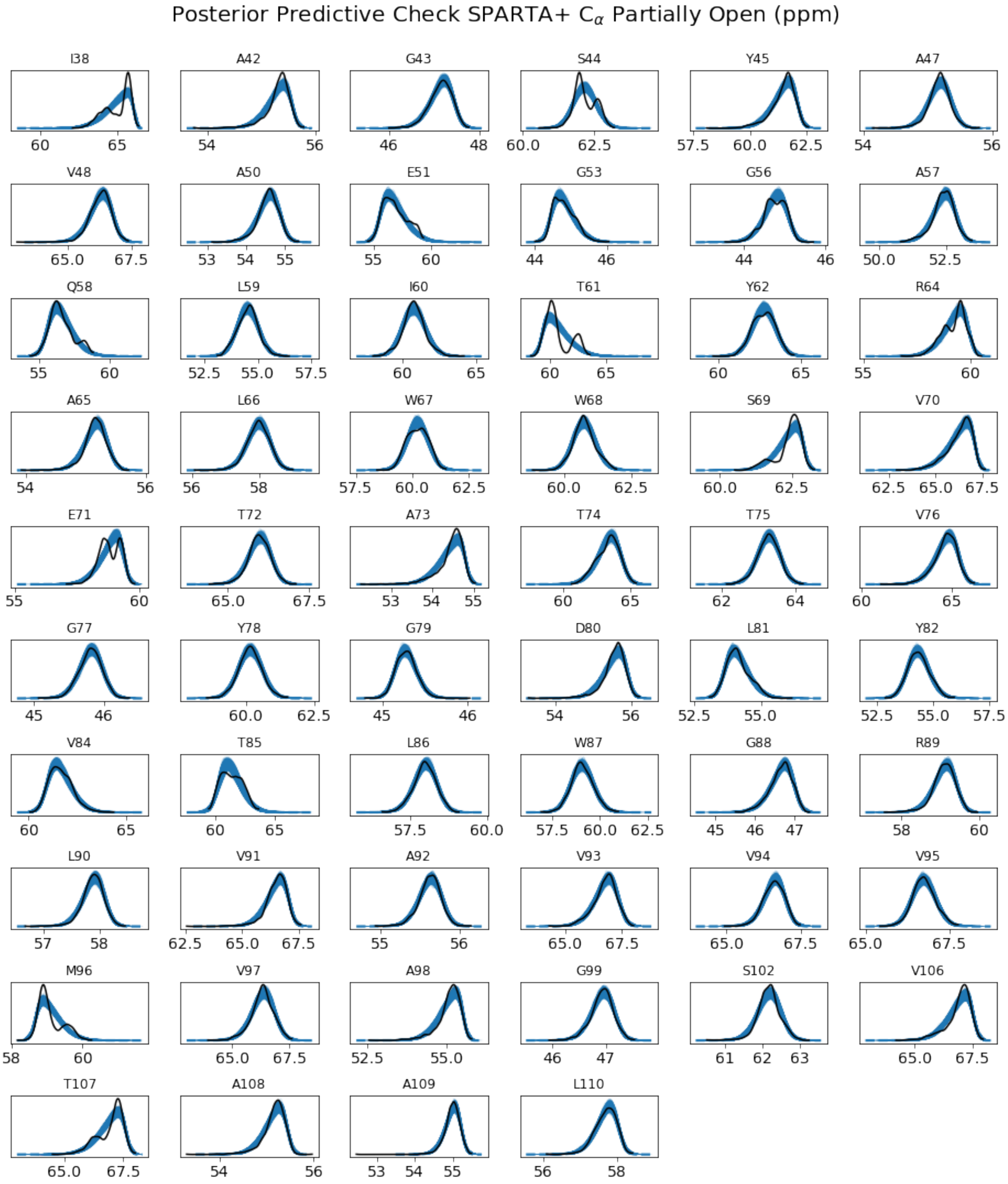


FIG. S3. Posterior predictive checks for $C_\alpha$ chemical shift data of the different residues calculated for the Partially Open state simulation with the CS prediction method SPARTA+. A posterior predictive check consists in sampling the posterior probability distributions ( for our case the posteriors of $\mu$, $\sigma$ and $\alpha$ parameters since the likelihood is a skew-gaussian distribution) to produce an ensemble of distributions (blue lines) that are compared to the data used for the inference (dashed black line). Similar posterior predictive checks have been obtained for the rest of states, methods and nuclei and can be found in `https://github.com/delemottelab/Informing_NMR_experiments_w_MD`

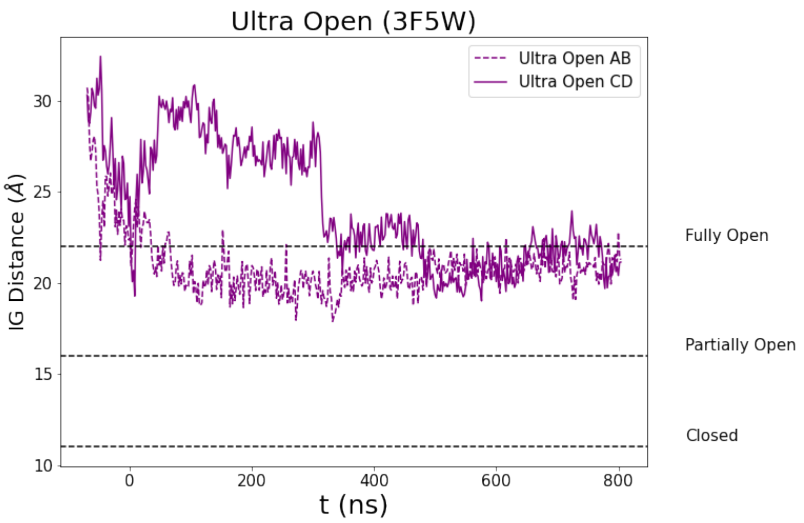


FIG. S4. Inner gate opening measured as the T112 $C_\alpha$ distance of opposing subunits ( subunits AB solid line or subunits CD dashed line) for the "Ultra Open" structure (PDB ID 3F5W). The structure is unstable and the inner gate rapidly decays to openings compatible with the Fully Open state (PDB ID 5VK6). For this reason this simulation was not analyzed in this work.

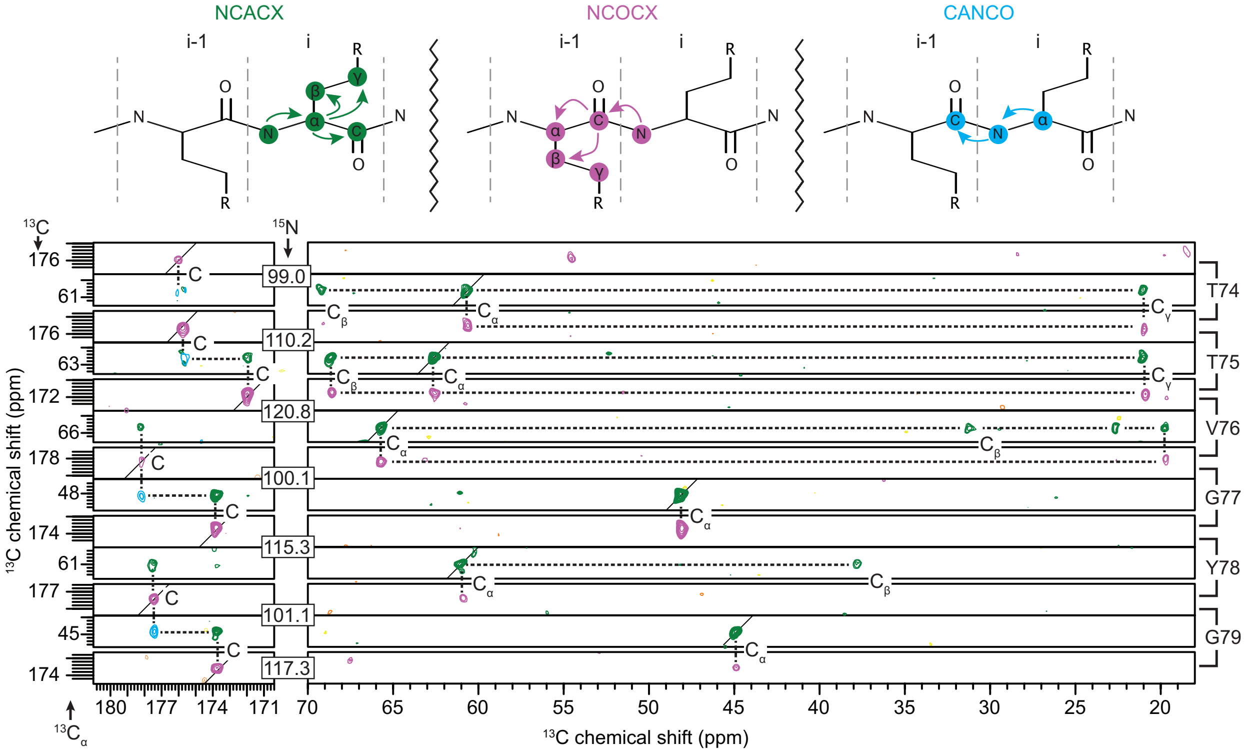


FIG. S5. Representative strip plot of the 3D experiments NCACX (green), NCOCX (magenta), CANCO (cyan) collected at 900 MHz that were used for the backbone walk of assignments of KcsA in the activated state (3:1 DOPE:DOPG, 50 mM KCl, pH 4.0). The polarization pathway of the experiments is shown (top). The backbone walk for residues T74 to G79 is shown.

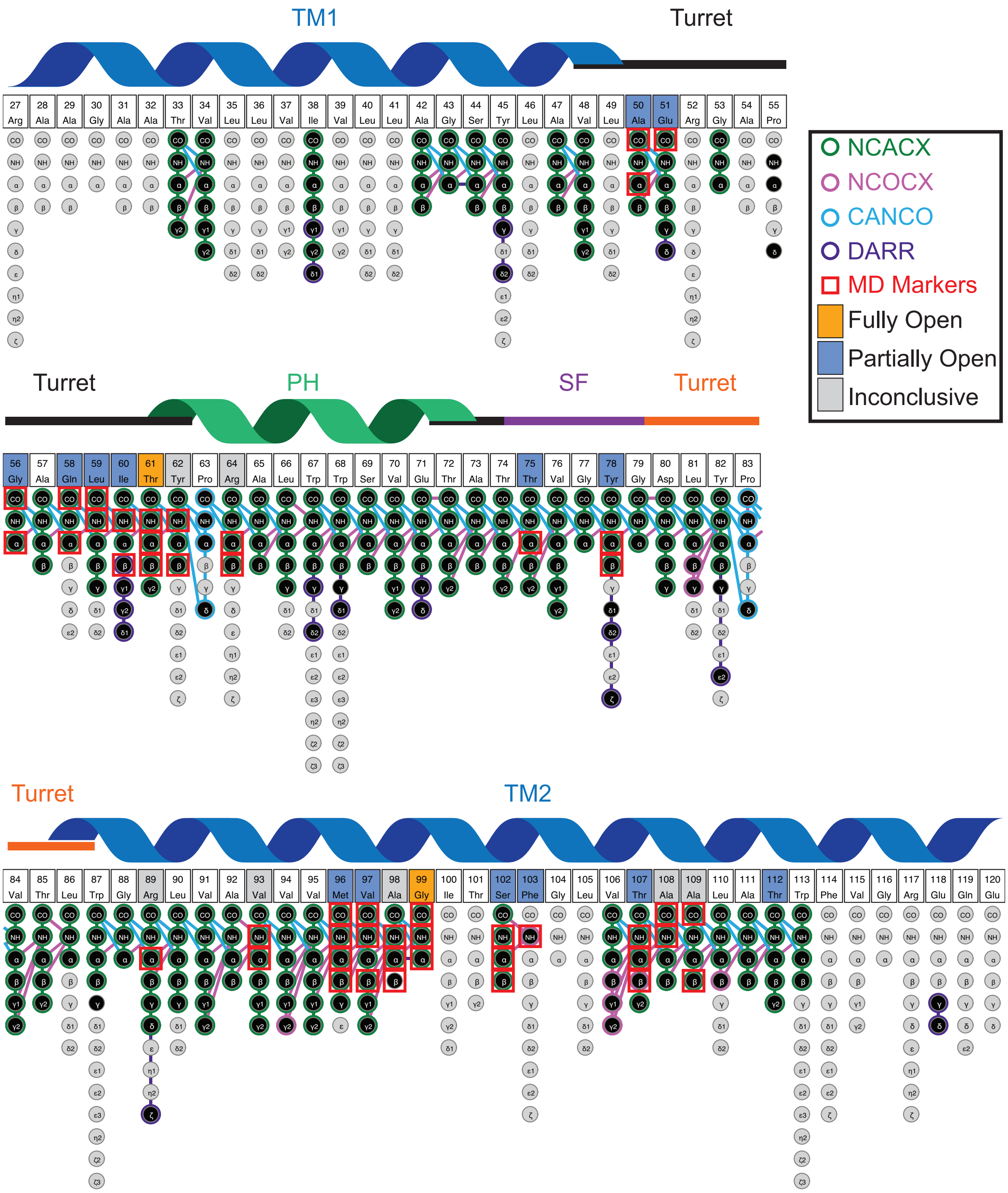


FIG. S6. Schematic indicating the assigned residues of KcsA in the activated state (3:1 DOPE:DOPG, 50 mM KCl, pH 4.0) with black filled in circles. The unassigned resonances are shown with gray filled circles. Open circles are used to indicate in which experiments the resonances are present. The 3D NCACX (green), NCOCX (magenta), and CANCO (cyan) and the 2D $^{13}$C-$^{13}$C DARR (purple) are shown, other assigned peaks are present in other experimental spectra that are not indicated on this figure (ZF-TEDOR (Pro), NcoCX, NcaCX, NCO, NCA). The resonances identified by statistical inference on the MD simulation data to be state markers are indicated with a red square and the agreement with the various states (Fully Open - orange, Partially Open - blue, Inconclusive - gray) are shown on the residue name and number.

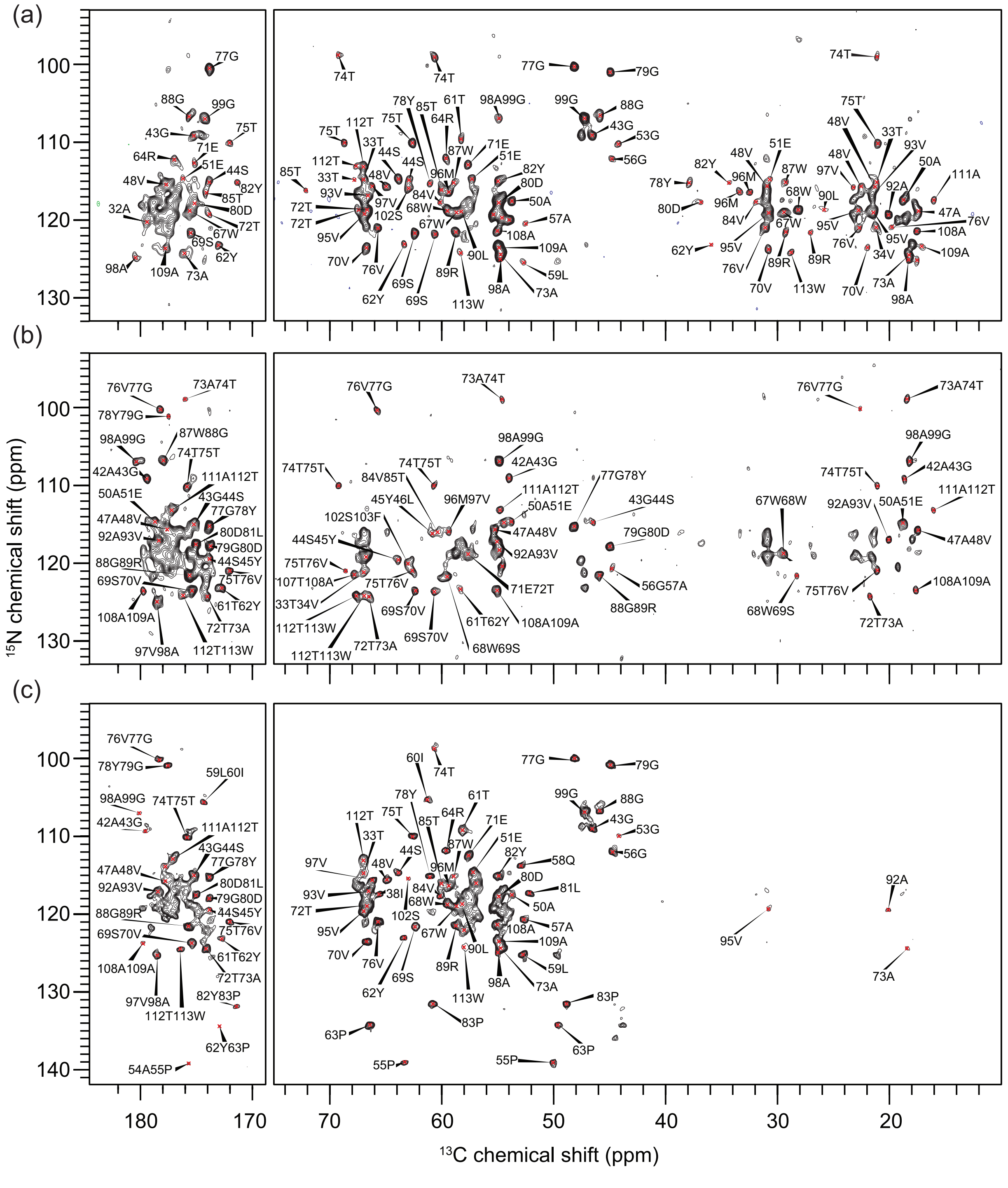


FIG. S7. 2D $^{13}$C-$^{15}$N NcaCX (a), NcoCX (b), and ZF-TEDOR (c) spectra of KcsA in the activated state (3:1 DOPE/DOPG, 50 mM KCl, pH 4.0) spectra collected at 900 MHz (a and b) and 750 MHz (c). Assigned resonances are labeled as either $C^{i}$-$N^{i}$ (NcaCX and ZF-TEDOR) or $C^{i-1}$-$N^{i}$ (NcoCX and ZF-TEDOR).

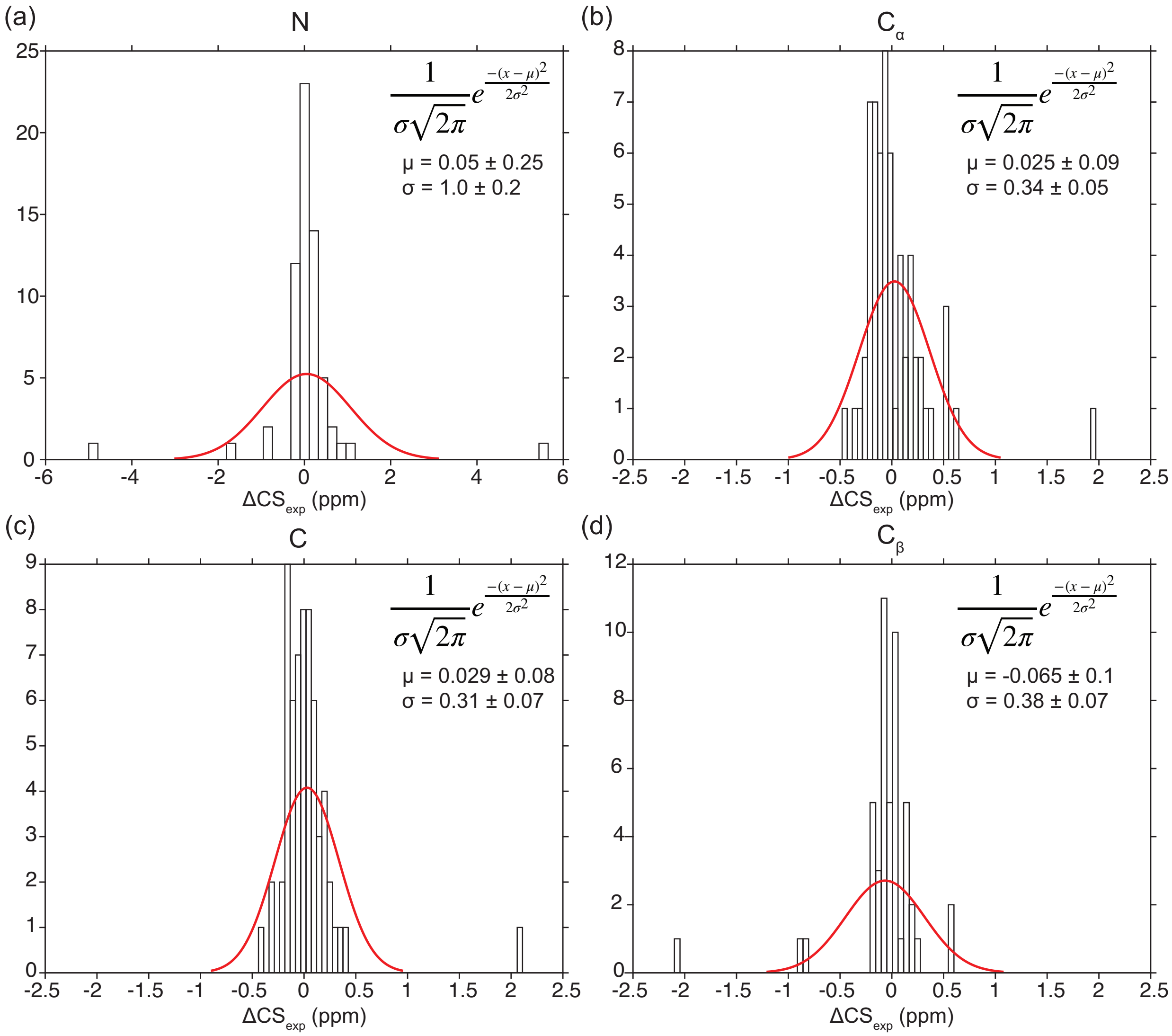


FIG. S8. Histograms of the experimental difference ($\Delta CS_{exp} = CS_{exp}^{pH=4,act} - CS_{exp}^{pH=7.5,deact}$) between the activated and deactivated state assigned chemical shifts for (a) N, (b) $C_\alpha$ , (c) C, and (d) $C_\beta$ . A fitted normal distribution is shown on each histogram (red lines) with the distribution equation and parameters shown, inset, demonstrating the proper referencing of the datasets to one another.

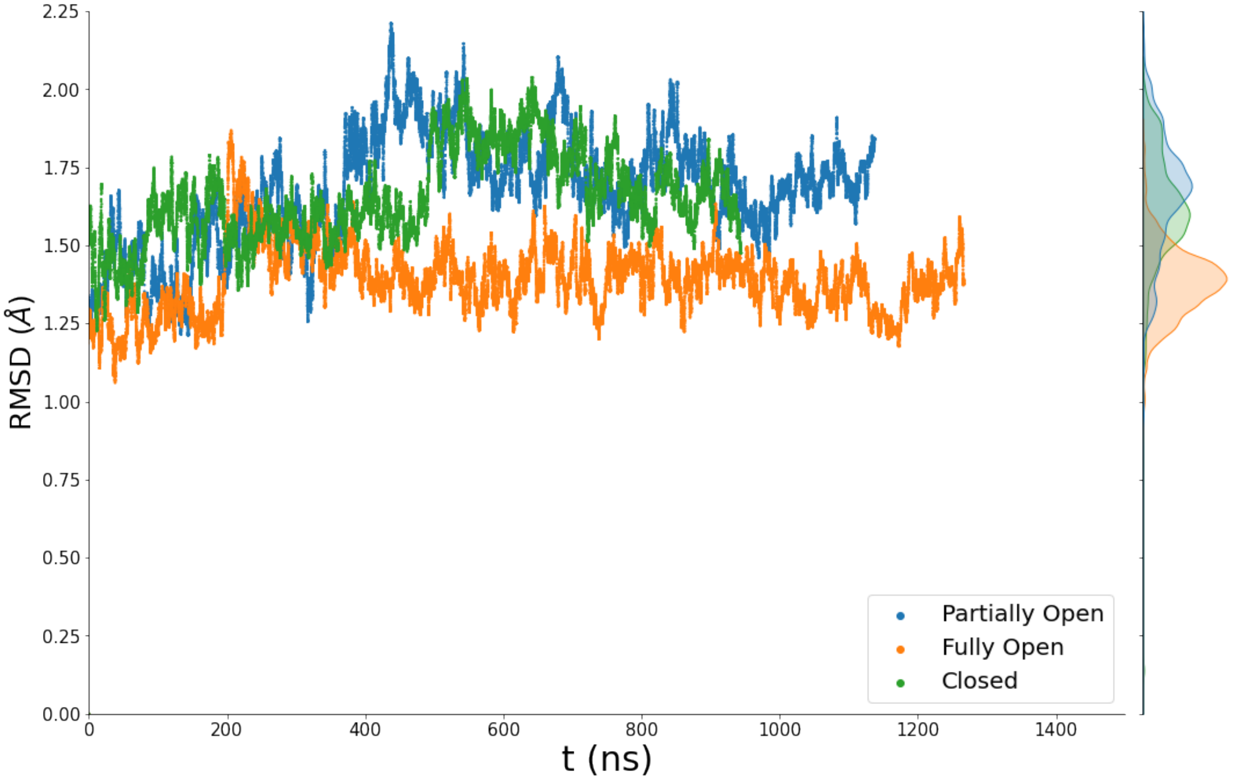


FIG. S9. Root mean square deviation (RMSD) of the $C_\alpha$ atom positions of the MD simulations as a function of time for the Partially Open, Fully Open and Closed states. Some of the noise is smoothed by a 10 step rolling median. Negative values of time are the equilibration period.

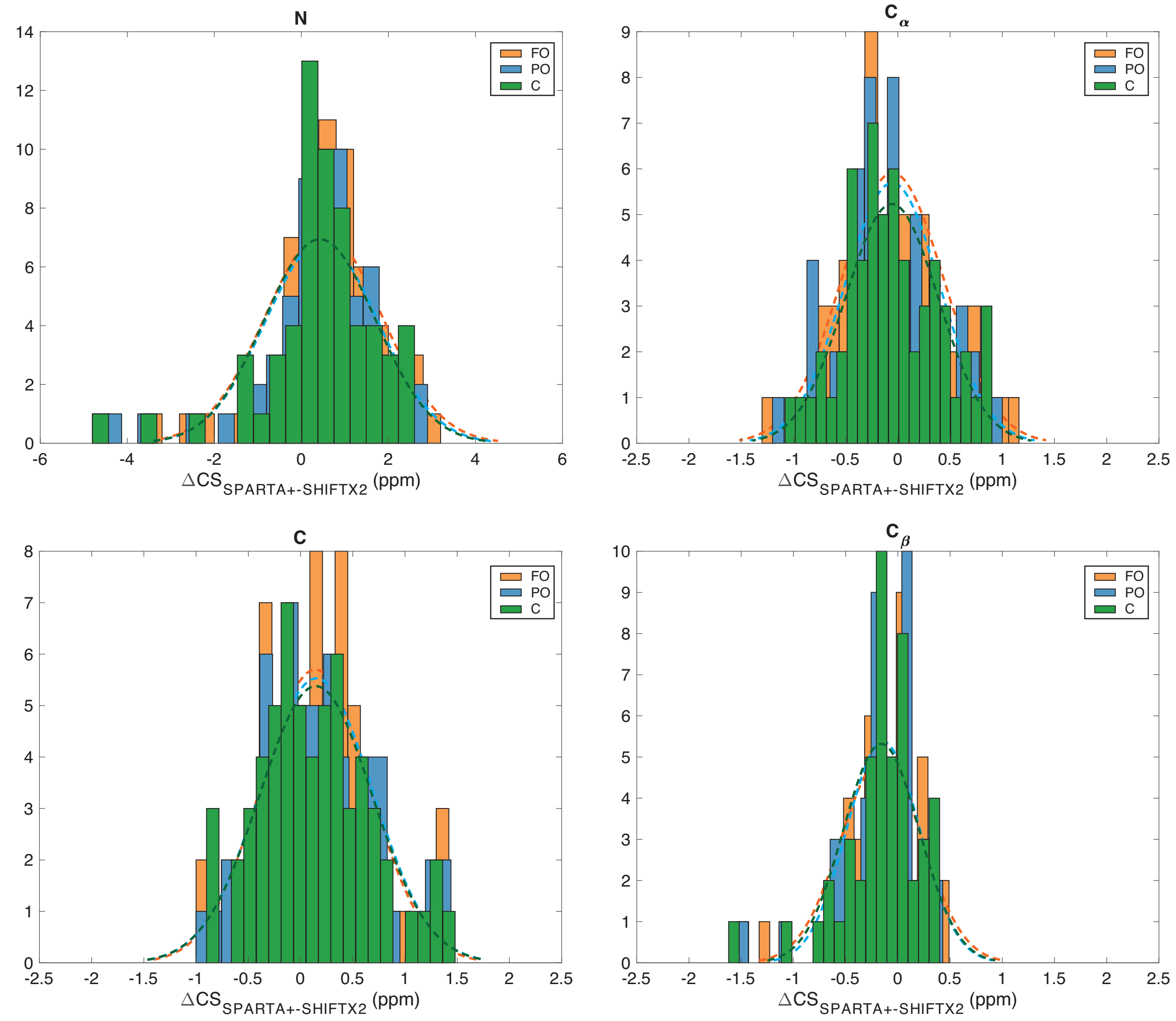


FIG. S10. Comparison of the predicted chemical shifts from the MD trajectories of the various states of KcsA using SPARTA+ and SHIFTX2. $\Delta CS_{\text{SPARTA+-SHIFTX2}} = CS^{X}_{\text{sim, SPARTA+}} - CS^{X}_{\text{sim, SHIFTX2}}$, where X is FO (orange), PO (blue), or Closed (green). The dashed lines represent a normal distribution fit to the histograms.

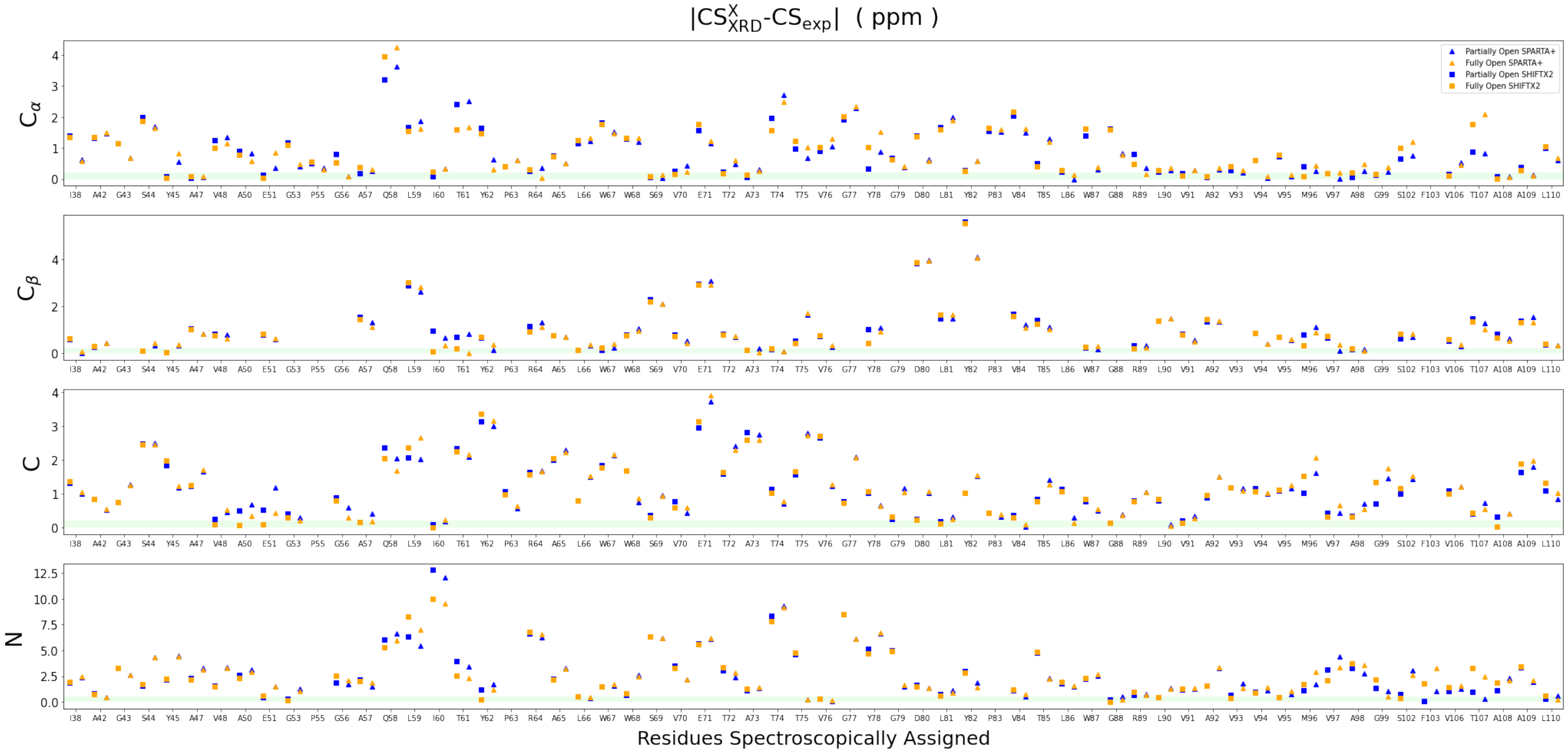


FIG. S11. Chemical shifts difference between experiment and the calculated CS for the XRD structures ($CS^X_{XRD} - CS_{exp}$), where X represents the FO (orange) or PO (blue) sate. All residues on the x-axis for the different nuclei and chemical shift prediction methods. The agreement between the calculation and the NMR experiments is higher the closer the value is to zero. Chemical shifts predicted by SPARTA+ and SHIFTX2 are drawn with triangles and squares respectively. The green shade depicts the typical experimental uncertainty.

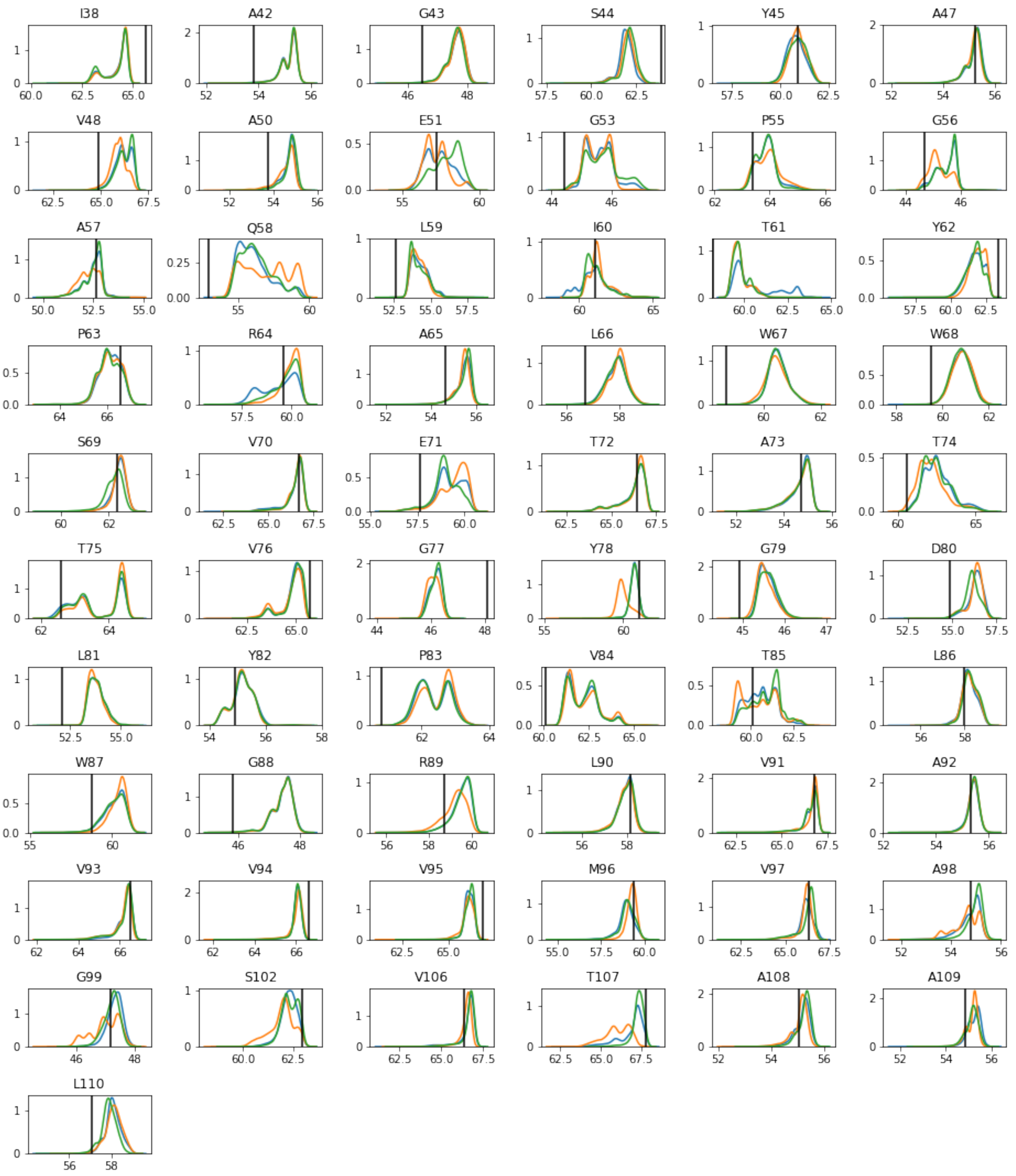


FIG. S12. Probability distributions of the raw simulated CS of the MD trajectory snapshots for the different residues assigned spectroscopically for the $C_\alpha$ nucleus and the CS prediction method SHIFTX2. Three different states are compared: Partially Open (blue), Fully Open (orange) and Closed (green). The vertical line indicates the NMR determined experimental CS for the activated state. The distributions are calculated using kernel density estimation as implemented in the python library Seaborn.

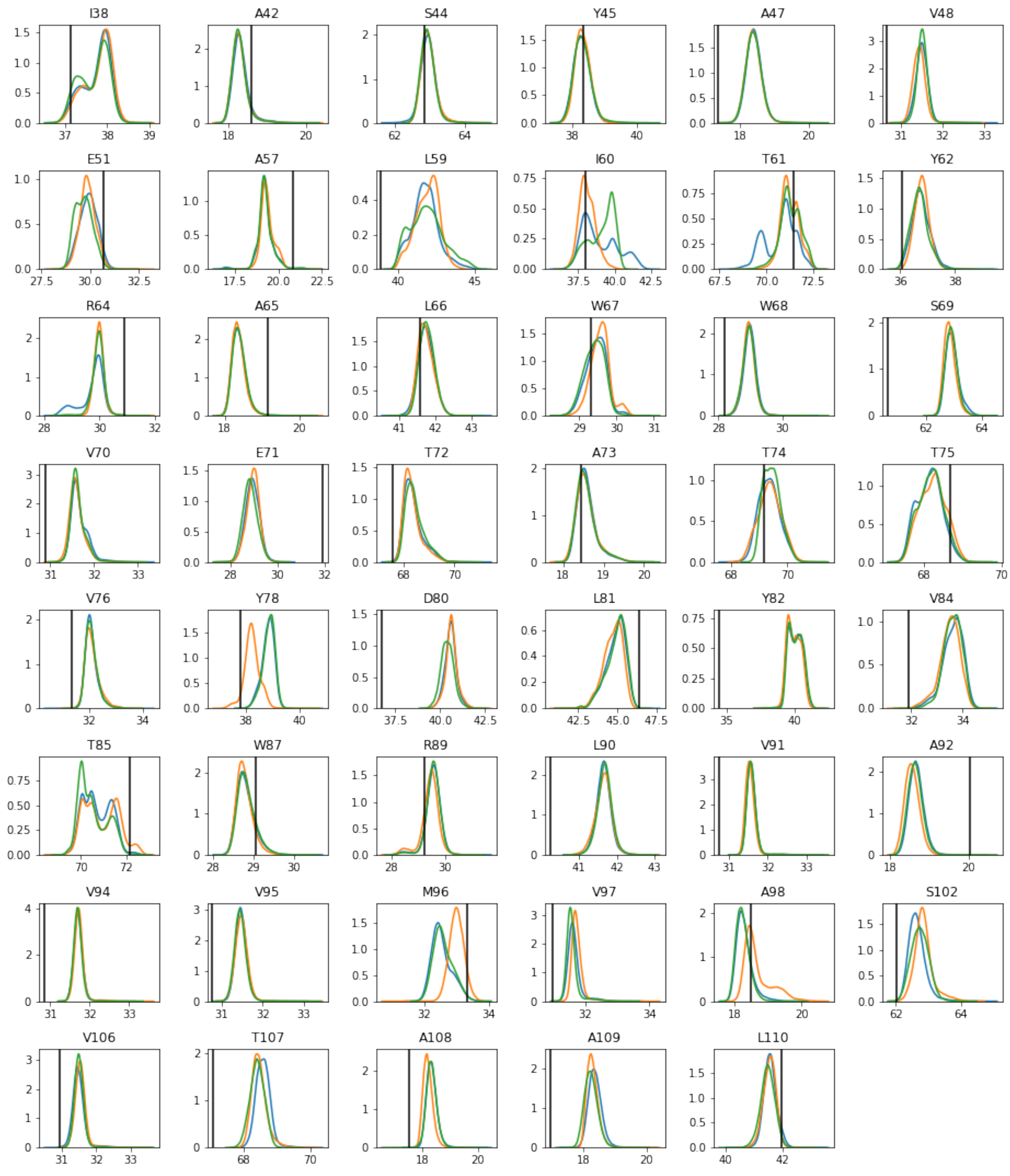


FIG. S13. Probability distributions of the raw simulated CS of the MD trajectory snapshots for the different residues assigned spectroscopically for the $C_\beta$ nucleus and the CS prediction method SHIFTX2. Three different states are compared: Partially Open (blue), Fully Open (orange) and Closed (green). The vertical line indicates the NMR determined experimental CS for the activated state. The distributions are calculated using kernel density estimation as implemented in the python library Seaborn.

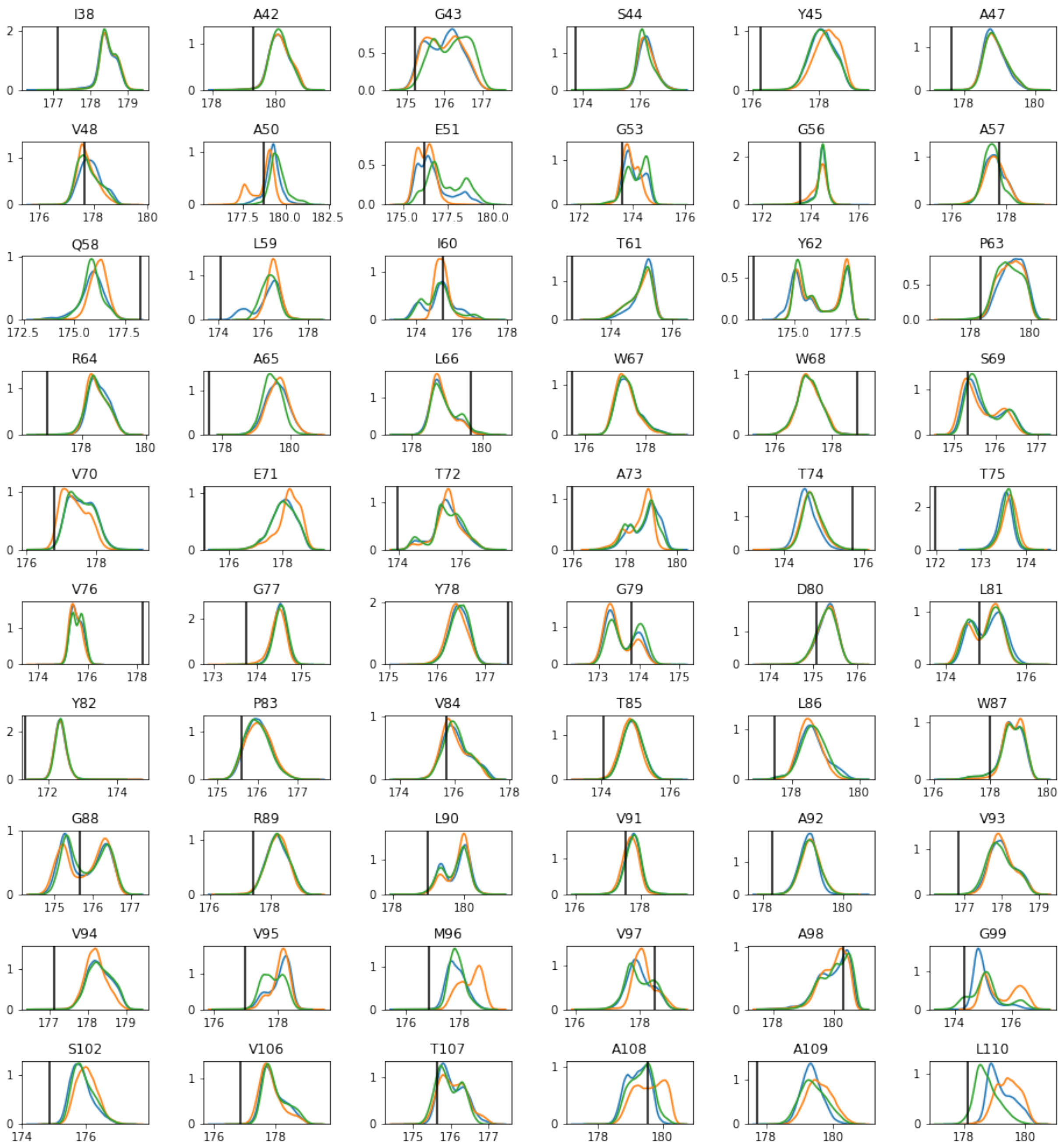


FIG. S14. Probability distributions of the raw simulated CS of the MD trajectory snapshots for the different residues assigned spectroscopically for the C nucleus and the CS prediction method SHIFTX2. Three different states are compared: Partially Open (blue), Fully Open (orange) and Closed (green). The vertical line indicates the NMR determined experimental CS for the activated state. The distributions are calculated using kernel density estimation as implemented in the python library Seaborn.

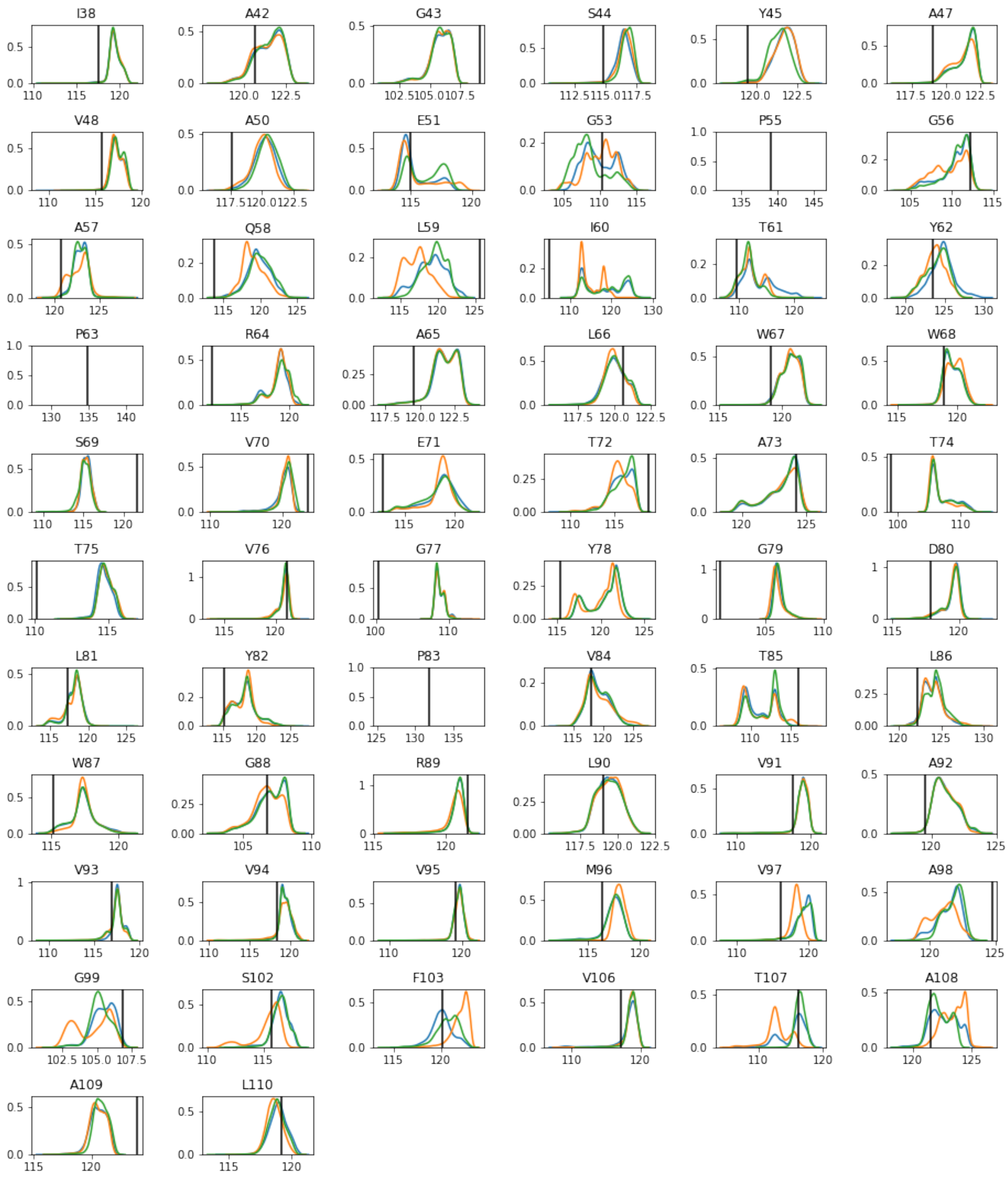


FIG. S15. Probability distributions of the raw simulated CS of the MD trajectory snapshots for the different residues assigned spectroscopically for the N nucleus and the CS prediction method SHIFTX2. Three different states are compared: Partially Open (blue), Fully Open (orange) and Closed (green). The vertical line indicates the NMR determined experimental CS for the activated state. The distributions are calculated using kernel density estimation as implemented in the python library Seaborn.

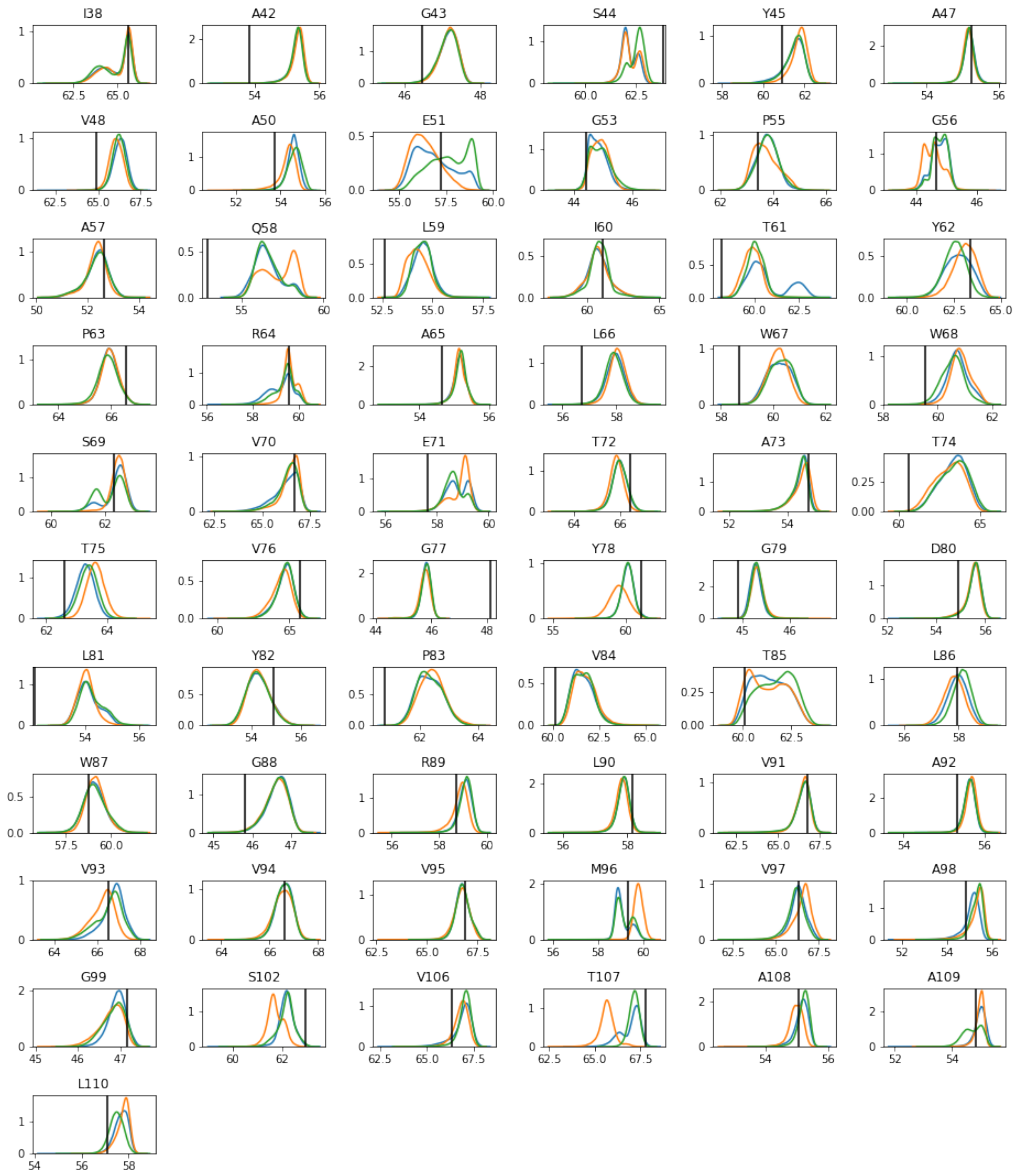


FIG. S16. Probability distributions of the raw simulated CS of the MD trajectory snapshots for the different residues assigned spectroscopically for the $C_\alpha$ nucleus and the CS prediction method SHIFTX2. Three different states are compared: Partially Open (blue), Fully Open (orange) and Closed (green). The vertical line indicates the NMR determined experimental CS for the activated state. The distributions are calculated using kernel density estimation as implemented in the python library Seaborn.

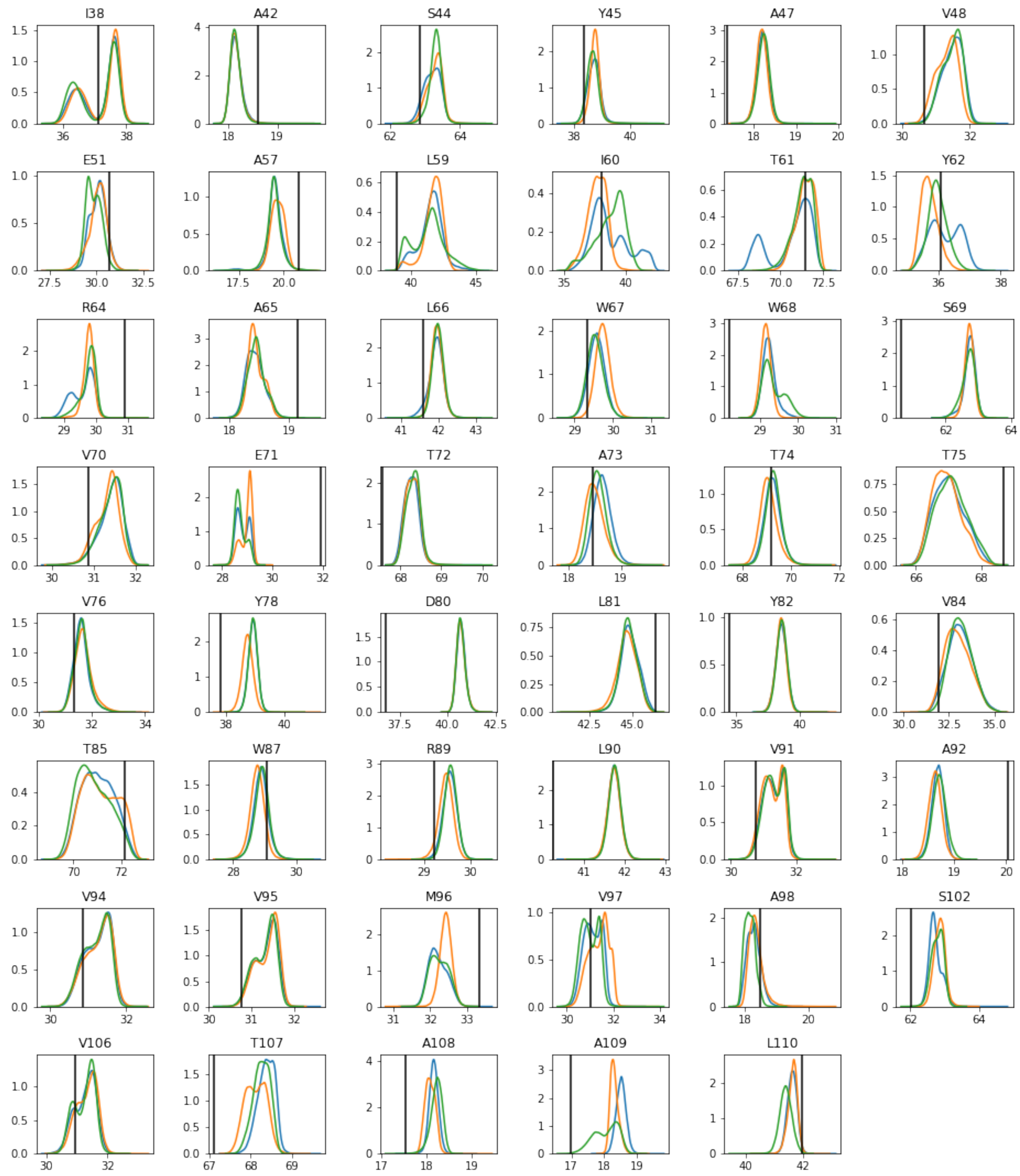


FIG. S17. Probability distributions of the raw simulated CS of the MD trajectory snapshots for the different residues assigned spectroscopically for the $C_\beta$ nucleus and the CS prediction method SHIFTX2. Three different states are compared: Partially Open (blue), Fully Open (orange) and Closed (green). The vertical line indicates the NMR determined experimental CS for the activated state. The distributions are calculated using kernel density estimation as implemented in the python library Seaborn.

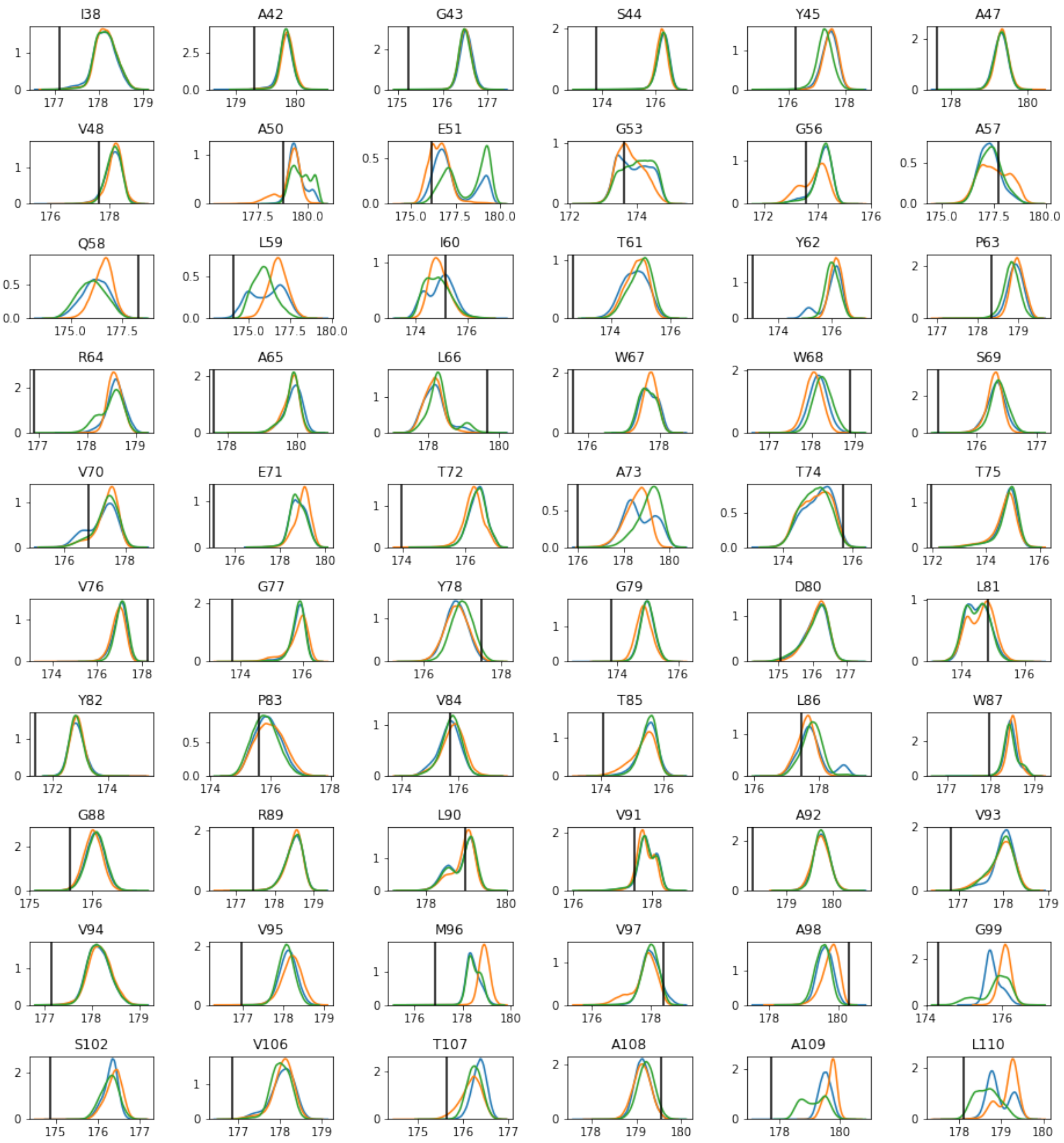


FIG. S18. Probability distributions of the raw simulated CS of the MD trajectory snapshots for the different residues assigned spectroscopically for the C nucleus and the CS prediction method SHIFTX2. Three different states are compared: Partially Open (blue), Fully Open (orange) and Closed (green). The vertical line indicates the NMR determined experimental CS for the activated state. The distributions are calculated using kernel density estimation as implemented in the python library Seaborn.

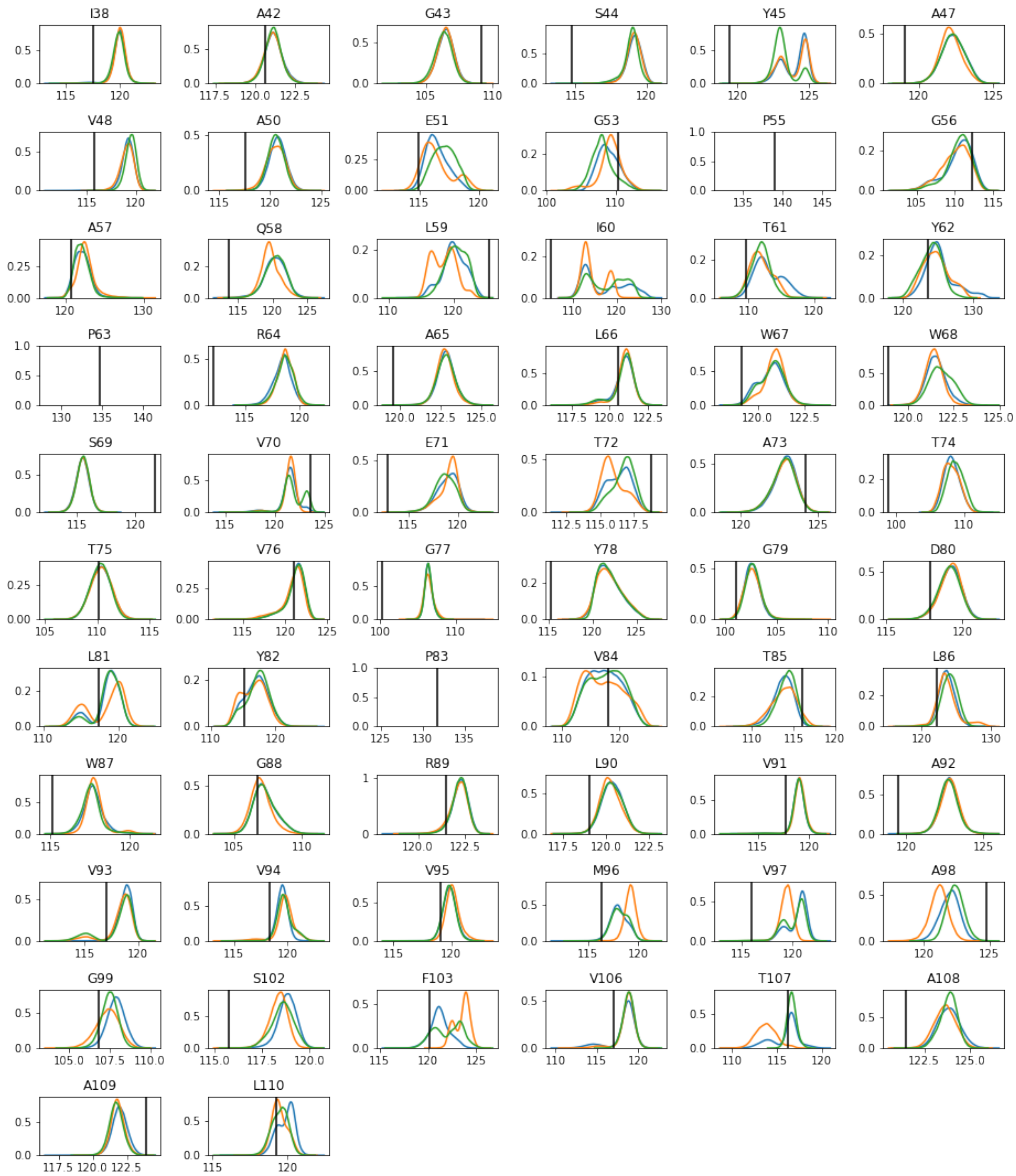


FIG. S19. Probability distributions of the raw simulated CS of the MD trajectory snapshots for the different residues assigned spectroscopically for the N nucleus and the CS prediction method SHIFTX2. Three different states are compared: Partially Open (blue), Fully Open (orange) and Closed (green). The vertical line indicates the NMR determined experimental CS for the activated state. The distributions are calculated using kernel density estimation as implemented in the python library Seaborn.

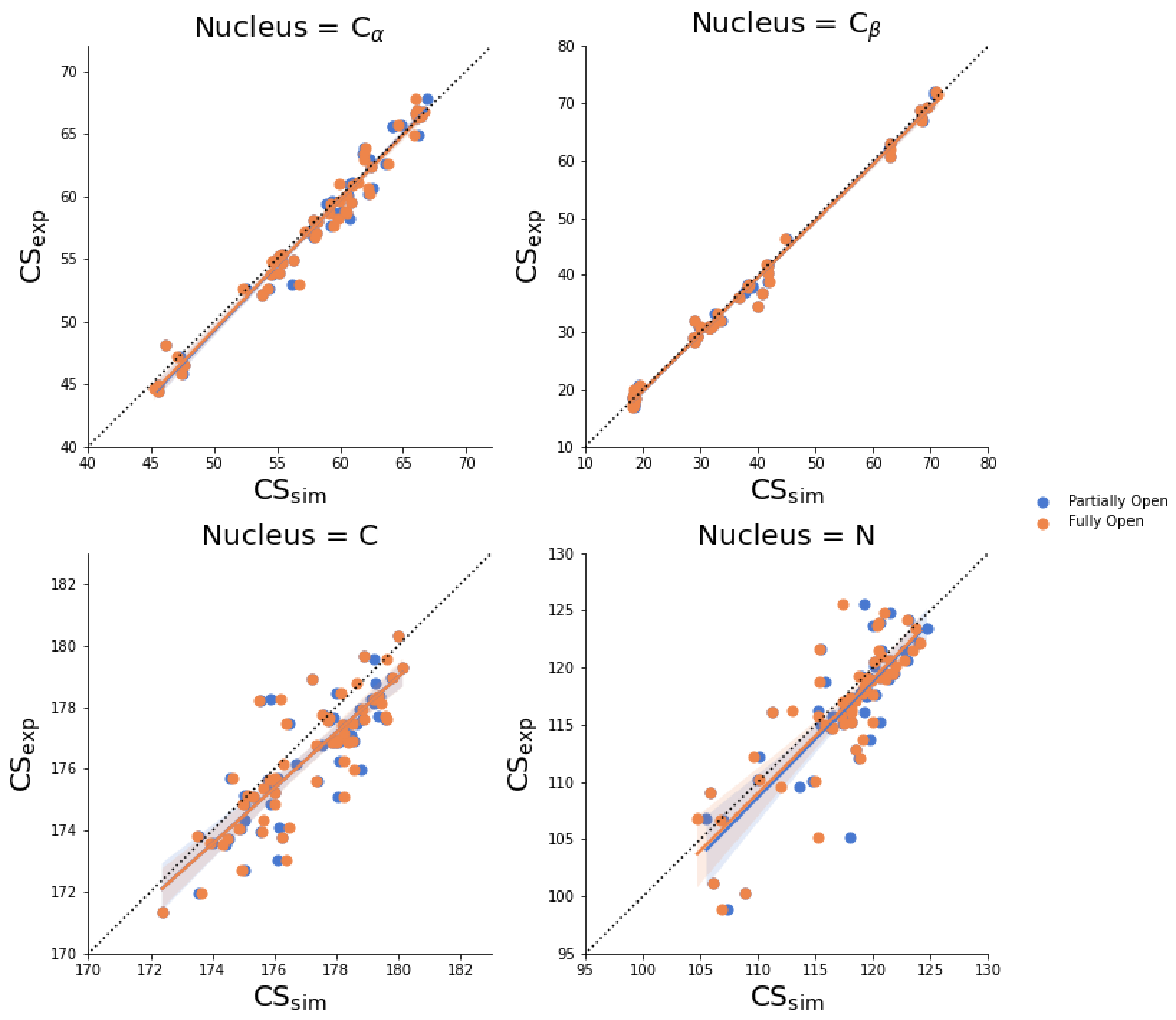


FIG. S20. Correlation diagram between experimental and simulated chemical shifts (shown in Figures S12-S15) for the Partially Open (blue) and Fully Open (orange) states for the nuclei studied and for the chemical shift prediction method SHIFTX2.

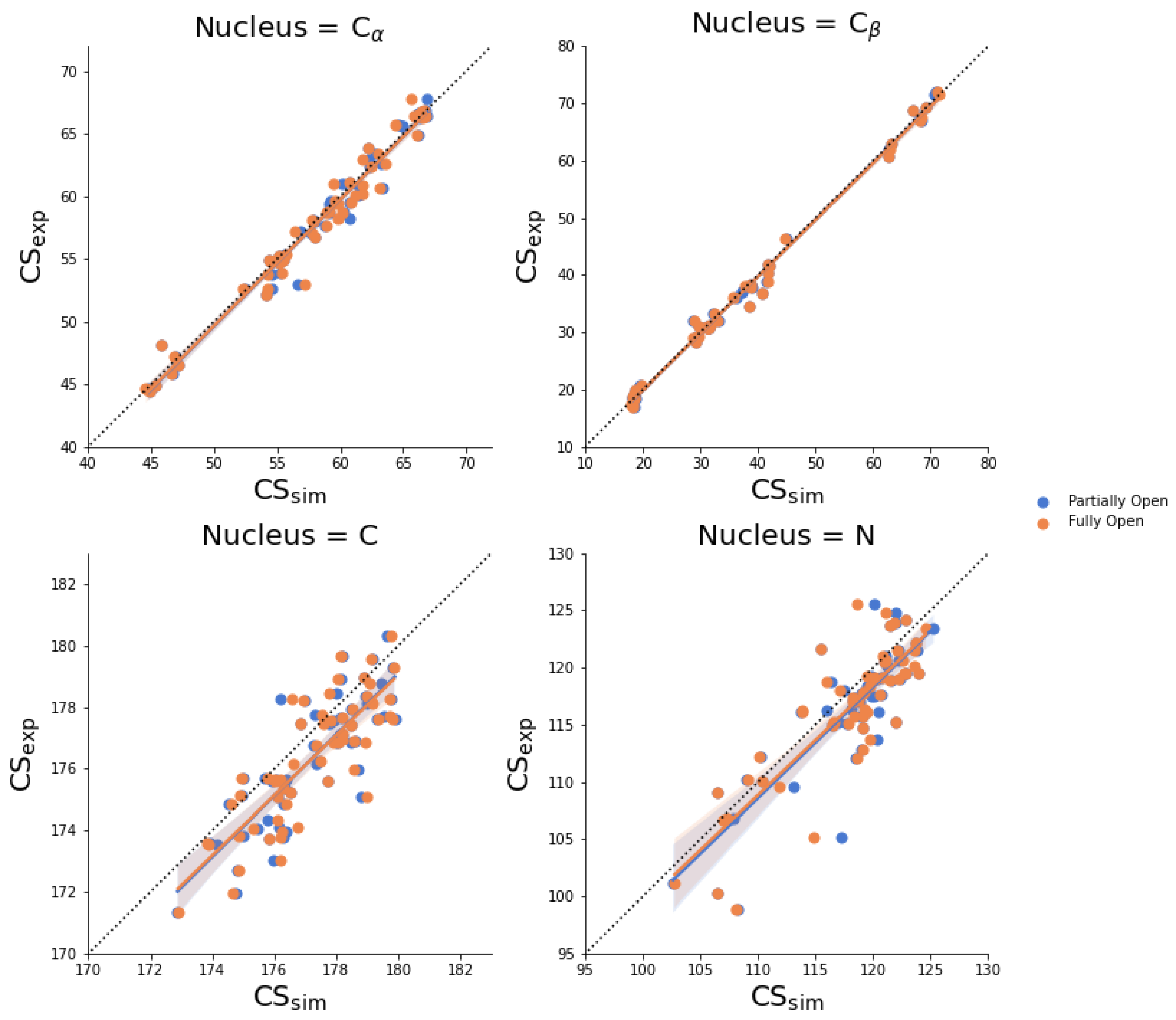


FIG. S21. Correlation diagram between experimental and simulated chemical shifts (shown in Figures S16-S19) for the Partially Open (blue) and Fully Open (orange) states for the nuclei studied and for the chemical shift prediction method SPARTA+.

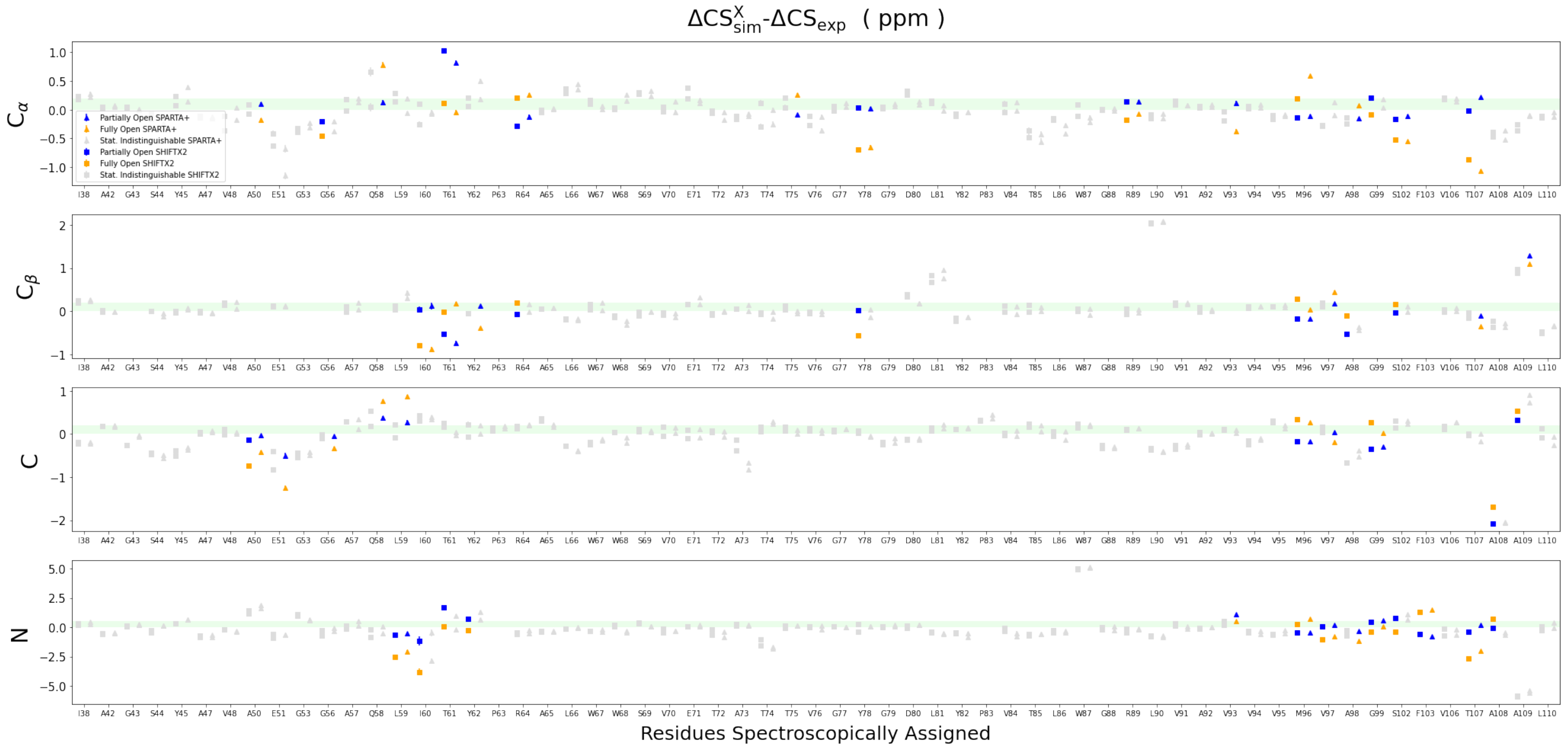


FIG. S22. Centers of 94% credible intervals of the difference in relative chemical shifts between experiment and simulation ($\Delta CS^X_{sim} - \Delta CS_{exp}$). The limits of the credible interval are shown as error bars. Both experiment and simulation use as reference the closed state chemical shifts: $\Delta CS^X_{sim} = CS^X_{sim}$ - $CS^C_{sim}$ and $\Delta CS_{exp} = CS^{pH=4}_{exp}$ - $CS^{pH=7.5}_{exp}$, where X represents the FO or PO sate. All residues on the x-axis for the different nuclei and chemical shift prediction methods. The agreement between MD simulations and the NMR experiments is higher the closer the value is to zero. Chemical shifts predicted by SPARTA+ and SHIFTX2 are drawn with triangles and squares respectively. The PO state (blue) has in general a better agreement with experiment than FO state (orange). The green shade depicts the typical experimental uncertainty. If two methods produce statistically identical CS or the signal is missing in the spectrum, the symbols are represented in gray.

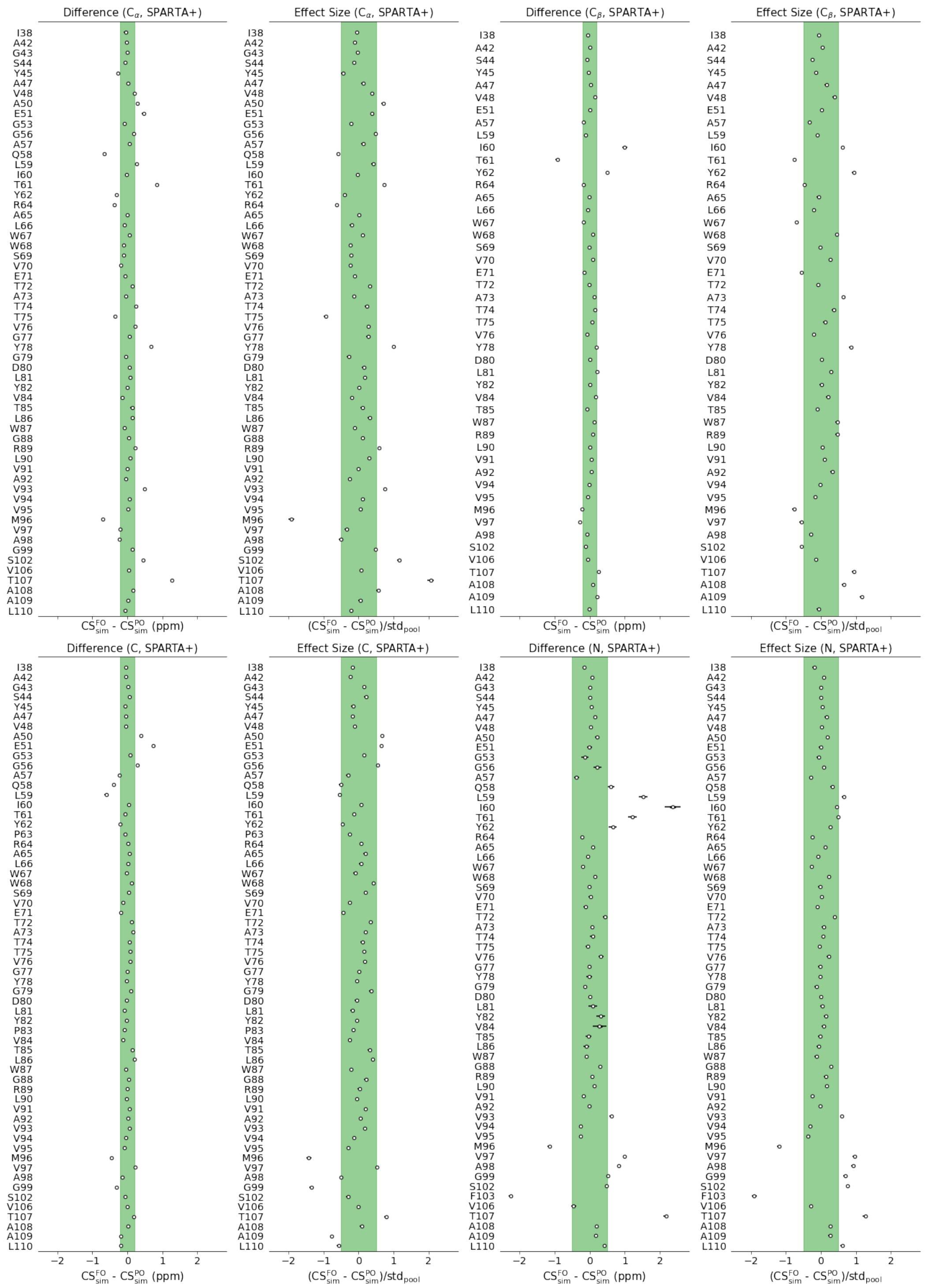


FIG. S23. Statistical filtering of simulated signals to discover discriminating residues using the CS prediction method SPARTA+. Horizontal bars represent the 94% credible interval of the variable distributions and circles represent its center.

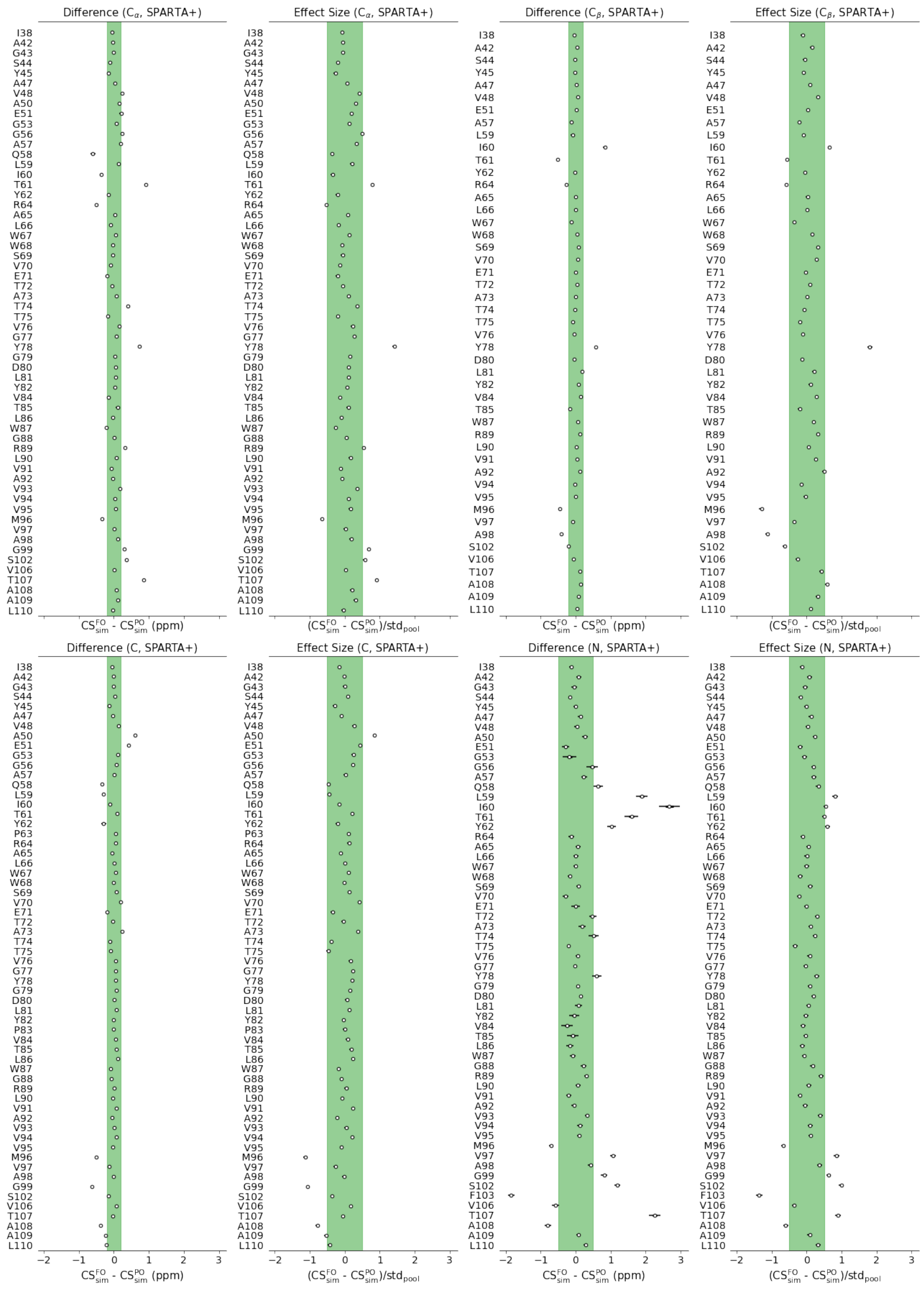


FIG. S24. Statistical filtering of simulated signals to discover discriminating residues using the CS prediction method SHIFTX2. Horizontal bars represent the 94% credible interval of the variable distributions and circles represent its center.

### SI: The dominant activated state of KcsA

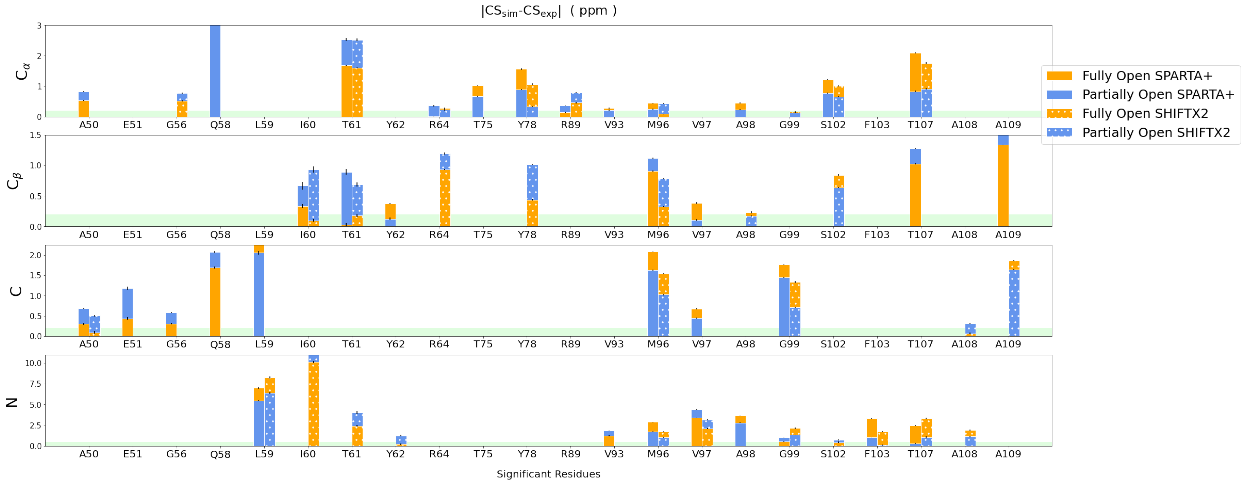


FIG. S25. Centers of 94% credible intervals for the difference in chemical shifts between experiment and simulation in absolute value ($|CS^{X}_{sim} - CS_{exp}|$). The limits of the credible interval are shown as error bars. The residues found to be discriminating residues using our statistical criteria are represented on the x-axis for the different nuclei and chemical shift prediction methods. The agreement to experiment is higher the closer the value is to zero. The green shade depicts the typical experimental uncertainty. The absence of a reference state increases the noise in the data.

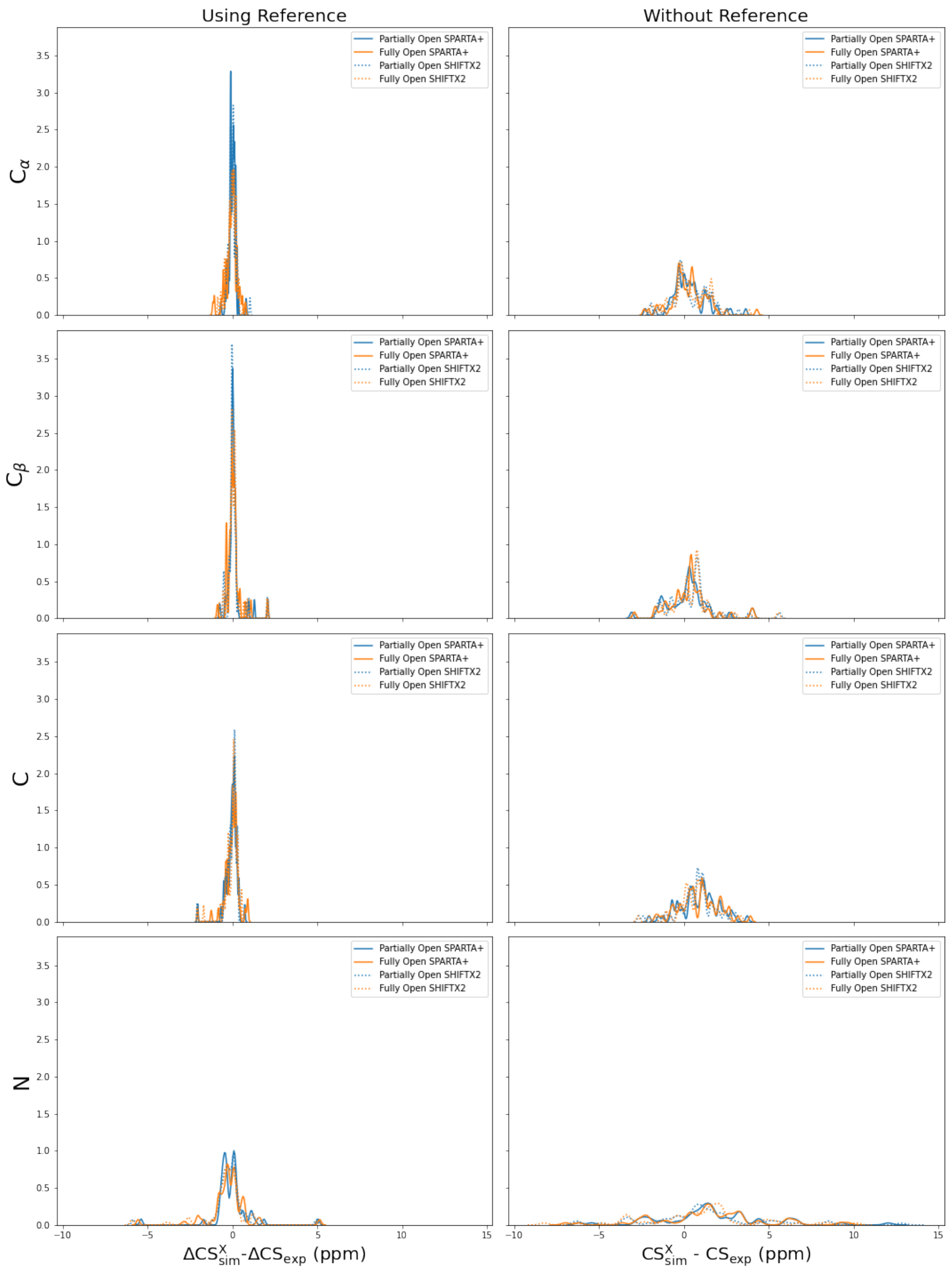


FIG. S26. Distributions of the CS simulation-experiment difference for the different nuclei (rows), for the two CS prediction methods (SPARTA+ solid line and SHIFTX2 dotted line) and for the two conductive states (Partially Open, blue and Fully Open, orange). These differences with experiment can be done using the CS of the closed state as a reference ($\Delta CS^X_{sim} - \Delta CS_{exp}$, left column) or without a reference ($CS^X_{sim} - CS_{exp}$ right column). The distributions without using the closed reference state are sharper and therefore using a reference state cancels random and systematic errors. The distributions are calculated using kernel density estimation as implemented in the python library Seaborn.

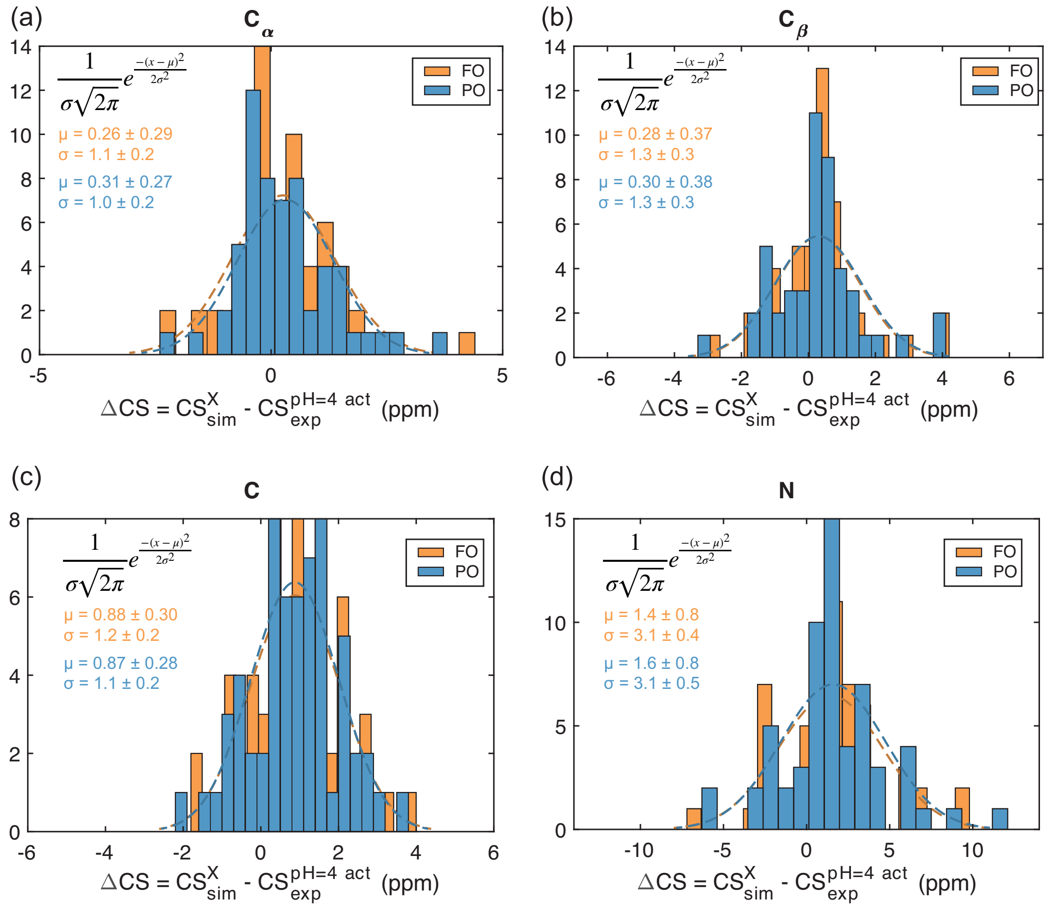


FIG. S27. Distributions of the CS simulation-experiment difference, $CS^{X}_{sim} - CS_{exp}$, for SPARTA+ and for the two conductive states (Partially Open, blue and Fully Open, orange). Each distribution is fit to a normal distribution function, shown inset with fitted values for PO (blue), and FO (orange).

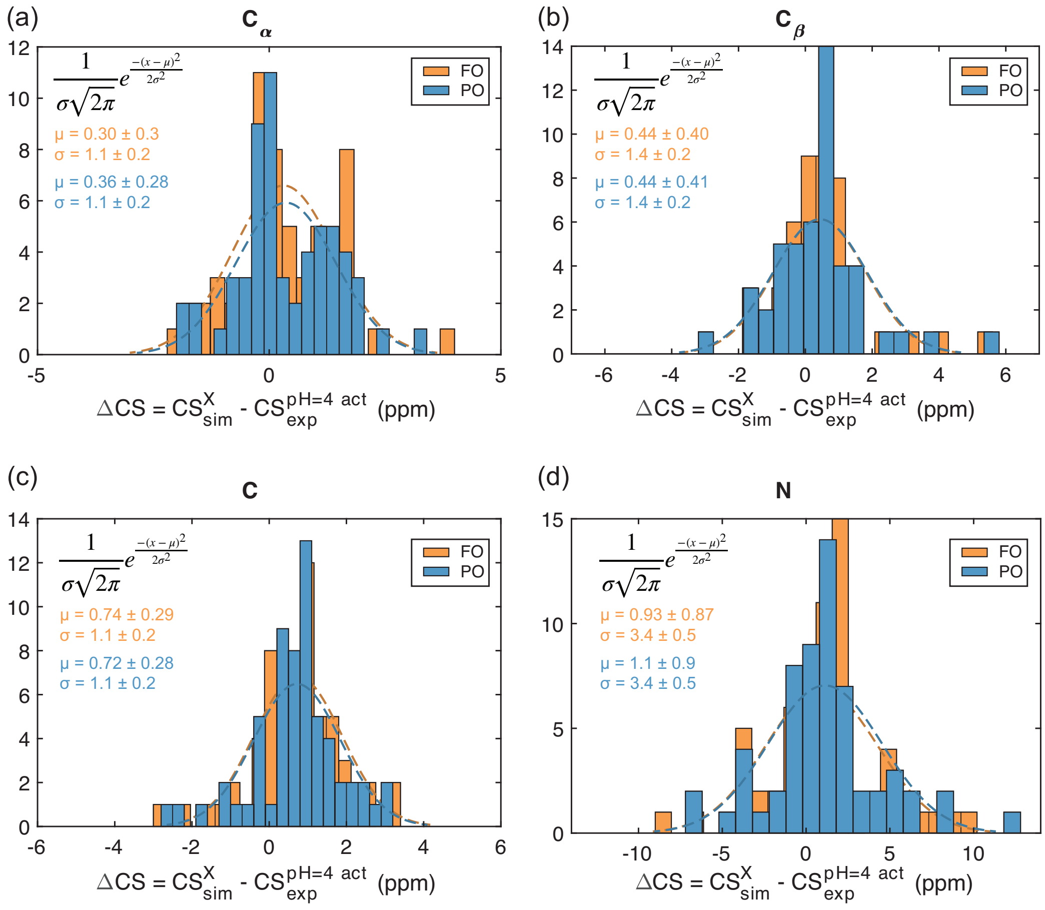


FIG. S28. Distributions of the CS simulation-experiment difference, $CS^X_{sim} - CS_{exp}$, for SHIFTX2 and for the two conductive states (Partially Open, blue and Fully Open, orange). Each distribution is fit to a normal distribution function, shown inset with fitted values for PO (blue), and FO (orange).

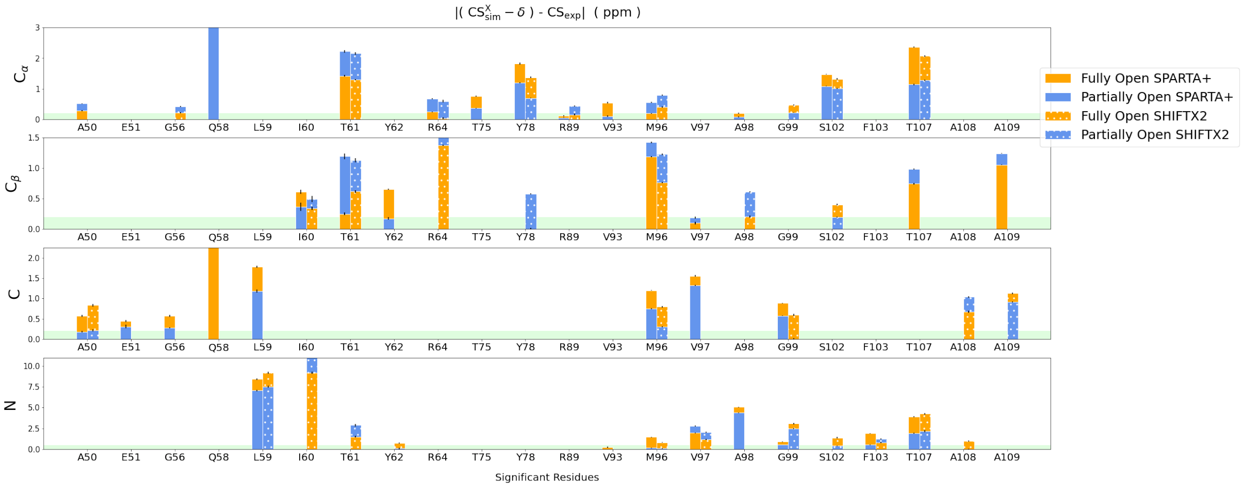


FIG. S29. Centers 94% credible intervals of the difference in chemical shifts between experiment and simulation in absolute value ($|(CS^X_{sim} - \delta) - CS_{exp}|$) where the simulated chemical shift has been corrected by $\delta$. $\delta$ is the position of the a fitted gaussian distribution to the chemical shifts of a nuclei-type (See Figures S28-S27). The limits of the credible interval are shown as error bars. The residues found to be discriminating residues using our statistical criteria are represented on the x-axis for the different nuclei and chemical shift prediction methods. The agreement to experiment is higher the closer the value is to zero. The green shade depicts the typical experimental uncertainty. The absence of a reference state increases the noise in the data.

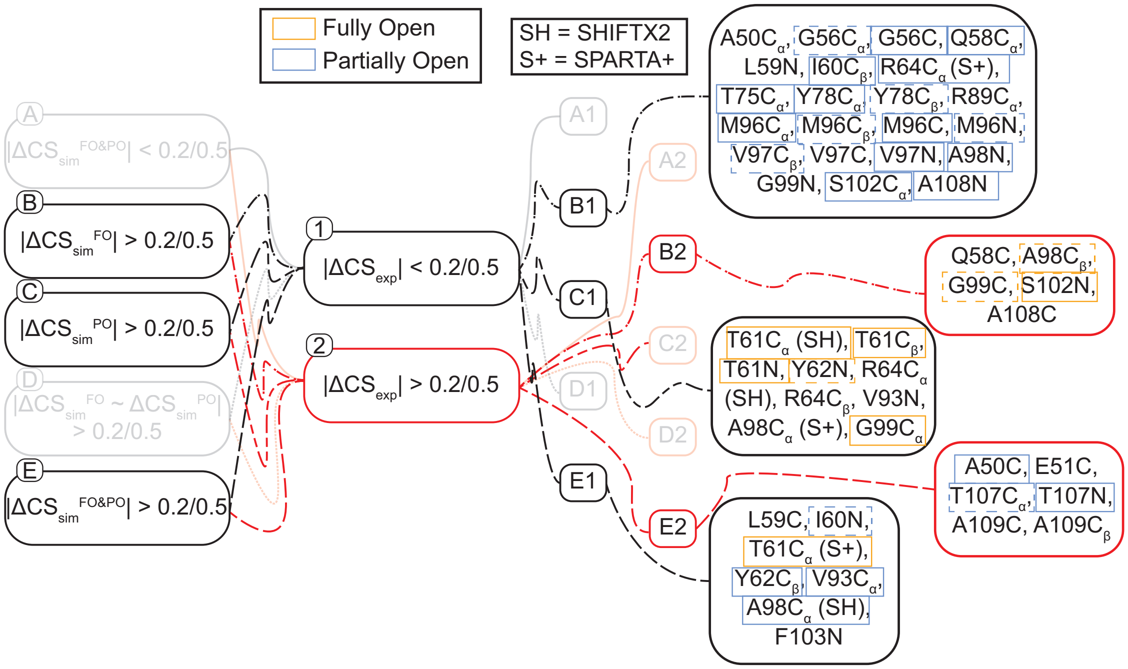


FIG. S30. Flow chart demonstrating the combinations of the $\Delta CS^{X}_{sim}$ and $\Delta CS_{exp}$ that yield various classifications of the state markers identified in this study. A threshold of tolerance of 0.2 ppm for $^{13}$C and 0.5 ppm for $^{15}$N, based on the experimental tolerance, was used to determine if the CS change was significant. The resonances that are identified by Bayesian inference from the MD simulation data as distinguishing between FO and PO are shown, on the right, within the classification that they were determined to belong. Two different classifications for each state marker are possible due to the two chemical shift prediction tools. These are marked with what chemical shift prediction tool was used for each resonance and classification type (SH = SHIFTX2 and S+ = SPARTA+). The state that determined to be likely based on the calculation of $\Delta\Delta CS^{X}$ is shown as an orange (FO) and blue (PO) box. Dashed boxes are used to indicate when one CS prediction tool was inconclusive and the other determined a state. Resonances that were determined to be inconclusive are shown without a thin box around them. FO = fully open, PO = partially open, C = closed, $\Delta CS^{X}_{sim} = CS^{X}_{sim} - CS^{Closed}_{sim}$, and $\Delta CS_{exp} = CS^{pH=4,\,act}_{exp} - CS^{pH=7.5,\,deact}_{exp}$.

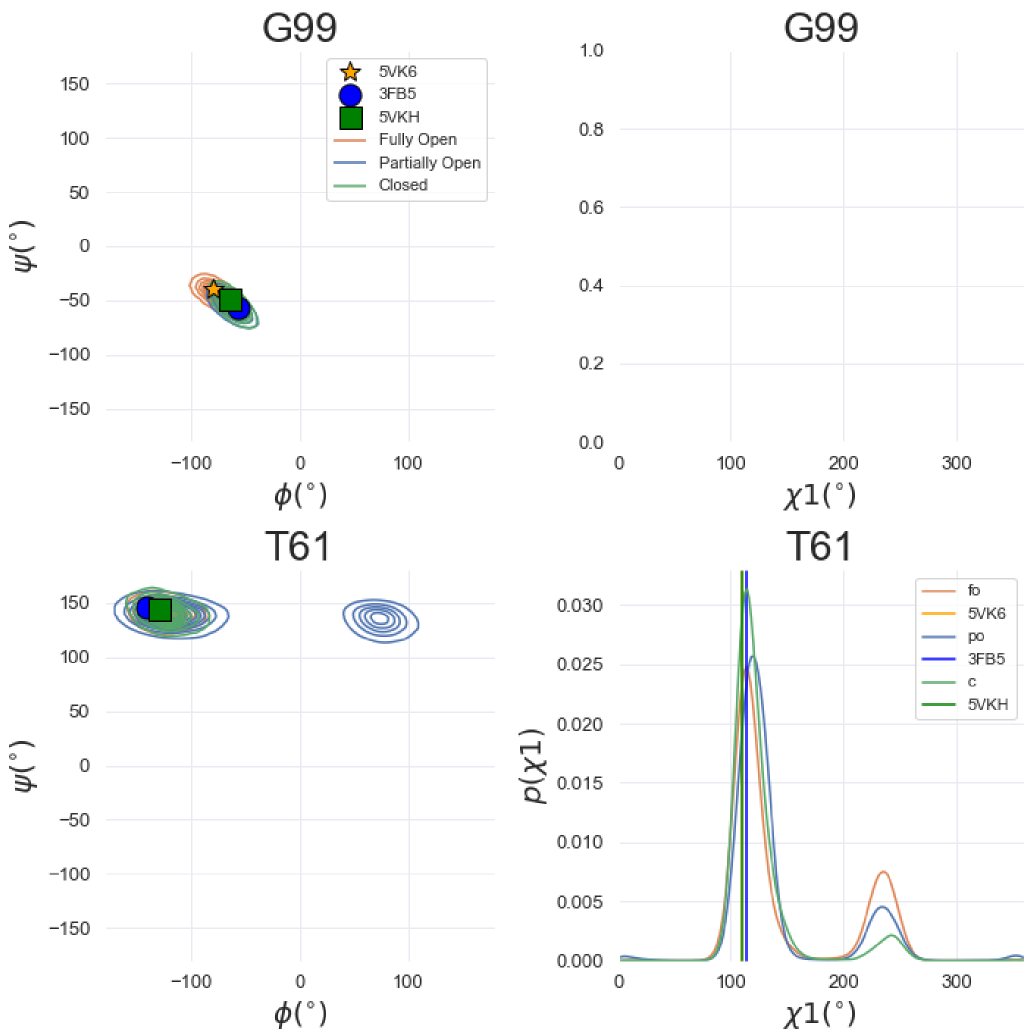


FIG. S31. Ramachandran and Janin diagrams (left and right columns, respectively) of the discriminating amino acids (rows) that are consistent with the Fully Open state as opposed to the Partially Open state consistent with most markers. The distributions are obtained from the dihedral angles obtained from the simulations of the Fully Open, Partially Open and Closed states. Additionally the values of the XRD structures are included as points or vertical lines.. Kernel density estimation is used to calculate the distributions as implemented in the Seaborn python library.

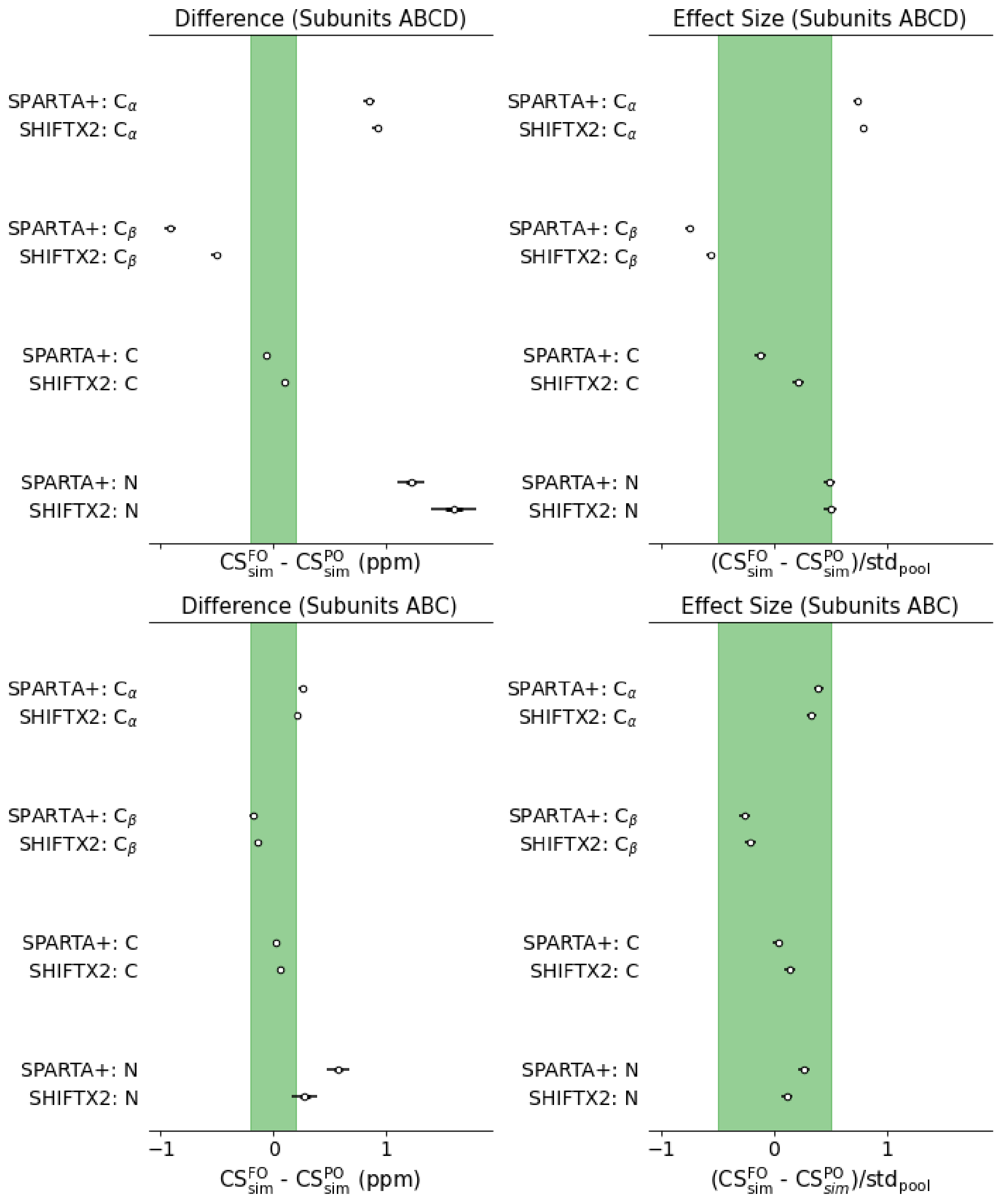


FIG. S32. Statistical filtering variables: difference in means and effect size for residue T61. The top graphs show the data using the four subunits of KcsA and the bottom graphs only subunits ABC. This figure shows that the reason why T61 is a discriminating residue is because one of the subunits has fluctuated away from the average. Horizontal bars represent the 94% credible interval of the variable distributions and circles represent its center.

#### II. SUPPLEMENTARY TABLES

TABLE S1: Experimental parameters for the 3D experiments at 900 MHz. Pulse sequences similar to those used for these experiments are available at `comdnmr.nysbc.org`

| Experiment | | 3D NCACX | 3D NCOCX | 3D CANCO |
|---|---|---|---|---|
| MAS frequency | | 16.666 kHz | 16.666 kHz | 16.666 kHz |
| First transfer | | H-N CP | H-N CP | H-C CP |
| | $\omega_{1,H}/2\pi$ (kHz) | 62 | 62 | 69 |
| | $\omega_{1,N/C}/2\pi$ (kHz) | N: 50 | N: 50 | C: 59 |
| | Pulse shape | $^{1}$H tangential | $^{1}$H tangential | Linear ramp |
| | Contact time (ms) | 0.8 | 0.8 | 0.8 |
| Second transfer | | N-C CP | N-C CP | C-N CP |
| | $\omega_{1,H}/2\pi$ (kHz) | 94 | 91 | 91 |
| | $\omega_{1,N/C}/2\pi$ (kHz) | C: 28<br>N: 42 | C: 41<br>N: 25 | C: 29<br>N: 42 |
| | Shape: nucleus and range | $^{13}$C tangential | $^{13}$C tangential | $^{13}$C tangential |
| | Contact time (ms) | 5 | 5.2 | 5.5 |
| Third transfer | | C-C mixing | C-C mixing | N-C CP |
| | $\omega_{1,H}/2\pi$ (kHz) | ∼17 | ∼17 | 98 |
| | $\omega_{1,N/C}/2\pi$ (kHz) | | | C: 43<br>N: 25 |
| | Shape: nucleus and range | | | $^{13}$C tangential |
| | Contact time (ms) | 50 | 50 | 4.5 |
| | $\omega_{1,H}/2\pi$ | | | 91 |
| Scans | | 16 | 32 | 64 |
| Points | $^{13}$C (direct) | 2048 | 2048 | 2048 |
| | $^{13}$C | 248 | 110 | 50 |
| | $^{15}$N | 60 | 80 | 48 |
| Acquisition time | $^{13}$C (direct) (ms) | 12.3 | 12.3 | 12.3 |
| | $^{13}$C (ms) | 6.6 | 6.6 | 7.2 |
| | $^{15}$N (ms) | 7 | 7.2 | 6 |

| | | | | |
|---|---|---|---|---|
| Sweep width | $^{13}$C (direct) (ppm) | 368 | 368 | 368 |
| | $^{13}$C (ppm) | 74 | 37 | 37 |
| | $^{15}$N (ppm) | 50 | 46 | 46 |
| Carrier Frequency | $^{13}$C (direct) (ppm) | 100.8 | 100.8 | 175.8 |
| | $^{13}$C (ppm) | 60.8 | 175.8 | 60.8 |
| | $^{15}$N (ppm) | 118.1 | 118.1 | 118.1 |
| Total time | | 5 d 12 h | 8 d | 7 d 14 h |

TABLE S2: Experimental parameters for the 3D experiments at 750 MHz. Pulse sequences similar to those used for these experiments are available at `comdnmr.nysbc.org`

| Experiment | | 3D NCACX | 3D NCOCX | 3D CANCO |
|---|---|---|---|---|
| MAS frequency | | 16 kHz | 16 kHz | 33.333 kHz |
| First transfer | | H-N CP | H-N CP | H-C CP |
| | $\omega_{1,\mathrm{H}}/2\pi$ (kHz) | 72 | 72 | 80 |
| | $\omega_{1,\mathrm{N/C}}/2\pi$ (kHz) | N: 58 | N: 58 | C: 56 |
| | Pulse shape | $^{1}$H tangential | $^{1}$H tangential | $^{1}$H tangential |
| | Contact time (ms) | 1 | 1 | 1 |
| Second transfer | | N-C CP | N-C CP | C-N CP |
| | $\omega_{1,\mathrm{H}}/2\pi$ (kHz) | 95 | 95 | 95 |
| | $\omega_{1,\mathrm{N/C}}/2\pi$ (kHz) | C: 24<br>N: 40 | C: 8<br>N: 24 | C: 17<br>N: 48 |
| | Shape: nucleus and range | $^{13}$C tangential | $^{13}$C tangential | $^{13}$C tangential |
| | Contact time (ms) | 3.75 | 5.0 | 3.75 |
| Third transfer | | C-C mixing | C-C mixing | N-C CP |
| | $\omega_{1,\mathrm{H}}/2\pi$ (kHz) | $\sim$16 | $\sim$16 | 95 |
| | $\omega_{1,\mathrm{N/C}}/2\pi$ (kHz) | | | C: 17<br>N: 49 |
| | Shape: nucleus and range | | | $^{13}$C tangential |

| | | | | |
|---|---|---|---|---|
| | Contact time (ms) | 50 | 50 | 5.0 |
| | $\omega_{1,H}/2\pi$ | | | 95 |
| Scans | | 96 | 144 | 64 |
| Points | $^{13}C$ (direct) | 2000 | 2000 | 2100 |
| | $^{13}C$ | 90 | 60 | 130 |
| | $^{15}N$ | 32 | 32 | 36 |
| Acquisition time | $^{13}C$ (direct) (ms) | 12.8 | 12.8 | 13.44 |
| | $^{13}C$ (ms) | 5.625 | 5.625 | 7.02 |
| | $^{15}N$ (ms) | 6 | 6 | 7.8 |
| Sweep width | $^{13}C$ (direct) (ppm) | 414 | 414 | 414 |
| | $^{13}C$ (ppm) | 42.4 | 28.3 | 44.2 |
| | $^{15}N$ (ppm) | 35 | 35 | 33.7 |
| Carrier Frequency | $^{13}C$ (direct) (ppm) | 109.9 | 109.9 | 109.9 |
| | $^{13}C$ (ppm) | 58 | 176.5 | 58 |
| | $^{15}N$ (ppm) | 111.2 | 111.2 | 111.2 |
| Total time | | 5 d 1 h | 5 d 21 h | 5 d 7 h |

TABLE S3: Root mean square errors (RMSE) of the different CS calculations of this work in relation to their experimental counter parts. FO stands for the Fully Open state (5VK6) and PO for the Partially Open state (3FB5). In general these differences are close to the experimental uncertainty of about 0.2 ppm.

| | | RMSE (ppm) | | | |
|---|---|---|---|---|---|
| | | SPARTA+ | | SHIFTX2 | |
| Nuclei | Magnitude | PO | FO | PO | FO |
| all | $CS^{X}_{XRD} - CS_{exp}$ | 4.2 | 4.2 | 4.1 | 4.1 |
| all | $\Delta CS^{X}_{XRD} - \Delta CS_{exp}$ | 0.6 | 0.6 | 0.8 | 0.7 |
| all | $CS^{X}_{sim} - CS_{exp}$ | 4.5 | 4.4 | 4.6 | 4.5 |
| discriminating | $CS^{X}_{sim} - CS_{exp}$ | 3.2 | 4.4 | 8.0 | 7.0 |
| all | $\left(CS^{X}_{sim} - \delta\right) - CS_{exp}$ | 2.4 | 2.4 | 2.9 | 2.9 |
| discriminating | $\left(CS^{X}_{sim} - \delta\right) - CS_{exp}$ | 1.5 | 2.3 | 5.8 | 5.0 |
| all | $\Delta CS^{X}_{sim} - \Delta CS_{exp}$ | 0.4 | 0.5 | 0.4 | 0.6 |
| discriminating | $\Delta CS^{X}_{sim} - \Delta CS_{exp}$ | 0.2 | 0.6 | 0.4 | 1.2 |

TABLE S4: Experimental resonance assignments of KcsA in the activated state (50 mM KCl, pH 4.0, 3:1 DOPE/DOPG)

| **Residue** | **N** | **C** | **C$\alpha$** | **C$\beta$** | **C$\gamma$1** | **C$\gamma$2** | **C$\delta$** | **C$\varepsilon$** | **C$\zeta$** |
|---|---|---|---|---|---|---|---|---|---|
| T33 | 114.86 | 175.04 | 66.88 | 67.65 | 21.15 | | | | |
| V34 | 120.9 | 177.17 | 66.88 | 30.86 | 23.11 | 21.28 | | | |
| L35 | | | | | | | | | |
| L36 | | | | | | | | | |
| V37 | | | | | | | | | |
| I38 | 117.52 | 177.12 | 65.62 | 37.14 | 28.74 | 16.72 | 13.36 | | |
| V39 | | | | | | | | | |
| L40 | | | | | | | | | |
| L41 | | | | | | | | | |
| A42 | 120.63 | 179.3 | 53.82 | 18.6 | | | | | |
| G43 | 109.11 | 175.24 | 46.47 | | | | | | |
| S44 | 114.79 | 173.76 | 63.88 | 62.85 | | | | | |

| **Residue** | **N** | **C** | **C$\alpha$** | **C$\beta$** | **C$\gamma1$** | **C$\gamma2$** | **C$\delta$** | **C$\varepsilon$** | **C$\zeta$** |
|---|---|---|---|---|---|---|---|---|---|
| Y45 | 119.48 | 176.26 | 60.9 | 38.36 | 127.63 | | 131.78 | | |
| L46 | | | | | | | | | |
| A47 | 119.02 | 177.63 | 55.23 | 17.36 | | | | | |
| V48 | 115.78 | 177.66 | 64.91 | 30.67 | 22.54 | 21.11 | | | |
| L49 | | | | | | | | | |
| A50 | 117.62 | 178.76 | 53.73 | 18.81 | | | | | |
| E51 | 114.96 | 176.17 | 57.24 | 30.71 | 38.05 | | 183.14 | | |
| R52 | | | | | | | | | |
| G53 | 110.26 | 173.61 | 44.42 | | | | | | |
| A54 | | | | | | | | | |
| P55 | 139.04 | | 63.4 | | | | 50.05 | | |
| G56 | 112.17 | 173.56 | 44.67 | | | | | | |
| A57 | 120.69 | 177.73 | 52.64 | 20.83 | | | | | |
| Q58 | 113.76 | 178.25 | 52.95 | | | | | | |
| L59 | 125.54 | 174.09 | 52.65 | 38.85 | 24.25 | | | | |
| I60 | 105.13 | 175.14 | 61.12 | 38.03 | 25.37 | 17.49 | 13.18 | | |
| T61 | 109.6 | 172.71 | 58.19 | 71.46 | 22.13 | | | | |
| Y62 | 123.47 | 173.01 | 63.38 | 36.09 | | | | | |
| P63 | 134.74 | 178.35 | 66.55 | | | | 49.65 | | |
| R64 | 112.12 | 176.91 | 59.6 | 30.91 | | | | | |
| A65 | 119.54 | 177.6 | 54.65 | 19.15 | | | | | |
| L66 | 120.55 | 179.68 | 56.73 | 41.58 | 26.4 | | | | |
| W67 | 119.11 | 175.58 | 58.69 | 29.32 | | 110.12 | 128.32 | | |
| W68 | 118.92 | 178.9 | 59.53 | 28.19 | | 112.58 | 124.38 | | |
| S69 | 121.71 | 175.35 | 62.35 | 60.63 | | | | | |
| V70 | 123.63 | 176.78 | 66.68 | 30.88 | 21.85 | 23.03 | | | |
| E71 | 112.88 | 175.08 | 57.64 | 31.91 | 31.98 | | | 181.74 | |
| T72 | 118.8 | 173.96 | 66.44 | 67.56 | 21.7 | | | | |
| A73 | 124.23 | 175.96 | 54.74 | 18.45 | | | | | |

| Residue | N | C | Cα | Cβ | Cγ1 | Cγ2 | Cδ | Cε | Cζ |
|---|---|---|---|---|---|---|---|---|---|
| T74 | 98.91 | 175.71 | 60.61 | 69.18 | | 21.03 | | | |
| T75 | 110.12 | 171.97 | 62.6 | 68.68 | | 21.05 | | | |
| V76 | 121.06 | 178.22 | 65.74 | 31.34 | 22.65 | 19.76 | | | |
| G77 | 100.32 | 173.75 | 48.1 | | | | | | |
| Y78 | 115.3 | 177.49 | 61.04 | 37.81 | | | 130.03 | 117.55 | 157.46 |
| G79 | 101.11 | 173.82 | 44.91 | | | | | | |
| D80 | 117.85 | 175.07 | 54.9 | 36.75 | 178.96 | | | | |
| L81 | 117.33 | 174.85 | 52.15 | 46.35 | 24.35 | | | | |
| Y82 | 115.19 | 171.35 | 54.9 | 34.44 | | 128.52 | | 117.63 | |
| P83 | 131.79 | 175.6 | 60.82 | | | | 48.85 | | |
| V84 | 118.03 | 175.71 | 60.14 | 31.91 | 20.71 | 17.63 | | | |
| T85 | 116.09 | 174.05 | 60.1 | 72.15 | | 22.14 | | | |
| L86 | 122.16 | 177.47 | 57.97 | | | | | | |
| W87 | 115.14 | 177.96 | 58.77 | 29.05 | | 109.81 | | | |
| G88 | 106.74 | 175.65 | 45.82 | | | | | | |
| R89 | 121.49 | 177.44 | 58.72 | 29.22 | | 27.22 | 43.68 | | 158.38 |
| L90 | 119.02 | 178.97 | 58.16 | 40.26 | 25.68 | | | | |
| V91 | 117.73 | 177.55 | 66.78 | 30.75 | 23.11 | 21.35 | | | |
| A92 | 119.48 | 178.25 | 55.32 | 20.03 | | | | | |
| V93 | 117.01 | 176.84 | 66.54 | 31.08 | 22.95 | 21.69 | | | |
| V94 | 118.34 | 177.14 | 66.63 | 30.86 | | 21.14 | | | |
| V95 | 119.08 | 176.95 | 66.86 | 30.77 | 23.26 | 21.4 | | | |
| M96 | 116.37 | 176.84 | 59.35 | 33.33 | 32.46 | | | | |
| V97 | 116.06 | 178.45 | 66.33 | 31.01 | 23.01 | 21.63 | | | |
| A98 | 124.77 | 180.33 | 54.82 | 18.48 | | | | | |
| G99 | 106.86 | 174.32 | 47.17 | | | | | | |
| I100 | | | | | | | | | |
| T101 | | | | | | | | | |
| S102 | 115.73 | 174.86 | 62.94 | 62.02 | | | | | |

| **Residue** | **N** | **C** | **C$\alpha$** | **C$\beta$** | **C$\gamma1$** | **C$\gamma2$** | **C$\delta$** | **C$\varepsilon$** | **C$\zeta$** |
|---|---|---|---|---|---|---|---|---|---|
| F103 | 120.2 | | | | | | | | |
| G104 | | | | | | | | | |
| L105 | | | | | | | | | |
| V106 | 117.11 | 176.85 | 66.37 | 30.95 | 23.17 | 21.42 | | | |
| T107 | 116.27 | 175.63 | 67.77 | 67.1 | | 19.64 | | | |
| A108 | 121.53 | 179.55 | 55.06 | 17.53 | | | | | |
| A109 | 123.94 | 177.72 | 54.85 | 16.97 | | | | | |
| L110 | 119.23 | 178.12 | 57.08 | 41.95 | | | | | |
| A111 | 117.65 | 177.21 | 54.94 | 16.05 | | | | | |
| T112 | 113.24 | 176.32 | 67.02 | 67.86 | 19.7 | | | | |
| W113 | 124.26 | 178.15 | 58.14 | 28.67 | | | | | |
| F114 | | | | | | | | | |
| V115 | | | | | | | | | |
| G116 | | | | | | | | | |
| R117 | | | | | | | | | |
| E118 | | | | | 34.31 | | 181.4 | | |

TABLE S5: Experimental resonance assignments of KcsA in the deactivated state (50 mM KCl, pH 7.5, 9:1 DOPE/DOPS) taken from previous data[1] with re-adjustment of the chemical shift referencing.

| Residue | N | C | C$\alpha$ | C$\beta$ | C$\gamma 1$ | C$\gamma 2$ | C$\delta$ | C$\varepsilon$ | C$\zeta$ |
|---|---|---|---|---|---|---|---|---|---|
| R27 | | 177.13 | 59.47 | 28.84 | | | | | |
| A28 | 119.6 | 179.78 | 54.855 | 17.74 | | | | | |
| A29 | 123.8 | 179.78 | 54.78 | 17.85 | | | | | |
| G30 | 106.75 | 174.13 | 47.1 | | | | | | |
| A31 | 122.8 | 178.18 | 55.025 | 18.22 | | | | | |
| A32 | | | | | | | | | |
| T33 | | | | | | | | | |
| V34 | 120.45 | 177.63 | 66.48 | 30.8 | | | | | |
| L35 | 118.2 | 177.38 | 57.82 | 41.27 | | | | | |
| L36 | 117.85 | 177.13 | 57.92 | 40.57 | | | | | |
| V37 | 117.6 | 177.33 | 67.085 | 30.92 | | | | | |
| I38 | 117.8 | 176.93 | 65.725 | 37.29 | 28.78 | 16.78 | 13.58 | | |
| V39 | 119.75 | 177.43 | 66.74 | 30.81 | | | | | |
| L40 | 118.3 | 179.28 | 58.135 | 41.33 | | | | | |
| L41 | 118.05 | 179.73 | 57.305 | 42.28 | | | | | |
| A42 | 120.15 | 179.48 | 53.855 | 18.58 | | | | | |
| G43 | 109.2 | 175.18 | 46.45 | | | | | | |
| S44 | 114.7 | 173.23 | 63.915 | 62.83 | | | | | |
| Y45 | 119.5 | 175.73 | 61.07 | 38.35 | | | | | |
| L46 | 116.55 | 177.18 | 56.9 | 41.11 | | | | | |
| A47 | 118.35 | 177.68 | 55.095 | 17.35 | | | | | |
| V48 | 115.85 | 177.68 | 64.895 | 30.88 | | | | | |
| L49 | 115.9 | 179.48 | 56.87 | 41.81 | | | | | |
| A50 | 119.3 | 178.98 | 53.92 | | | | | | |
| E51 | 114.95 | 176.43 | 57.405 | 30.59 | 38.38 | | 183.28 | | |
| R52 | 115.75 | | 58.57 | | | | | | |
| G53 | 110 | 173.28 | 44.205 | | | | | | |

| **Residue** | **N** | **C** | **C$\alpha$** | **C$\beta$** | **C$\gamma1$** | **C$\gamma2$** | **C$\delta$** | **C$\varepsilon$** | **C$\zeta$** |
|---|---|---|---|---|---|---|---|---|---|
| A54 | 123.55 | | 49.665 | 17.85 | | | | | |
| P55 | 139.3 | 174.98 | 63.2 | 30.97 | | | 49.78 | | |
| G56 | 112.3 | 173.53 | 44.495 | | | | | | |
| A57 | 120.8 | 177.93 | 52.8 | 20.83 | | | | | |
| Q58 | 113.95 | 178.48 | 53.135 | 31.26 | | | | | |
| L59 | 125.3 | 174.13 | 52.825 | 38.99 | | | 24.58 | | |
| I60 | 105.25 | 175.43 | 61.12 | 38.19 | 25.48 | 17.48 | 13.38 | | |
| T61 | 109.4 | 172.88 | 58.36 | 71.45 | | 22.28 | | | |
| Y62 | 123.5 | 173.08 | 63.355 | 35.97 | | | | | |
| P63 | 134.6 | 178.38 | 66.33 | 30.93 | | | 49.63 | | |
| R64 | 111.9 | 177.03 | 59.655 | 31.07 | | | | | |
| A65 | 119.2 | 177.78 | 54.69 | 19.23 | | | | | |
| L66 | 120.45 | 179.48 | 57.04 | 41.43 | | | | | |
| W67 | 118.85 | 175.38 | 58.825 | 29.31 | 110.48 | | | | |
| W68 | 118.9 | 178.93 | 59.505 | 28.07 | 113.08 | | | | |
| S69 | 121.8 | 175.48 | 62.42 | 60.59 | | | | | |
| V70 | 123.75 | 176.93 | 66.75 | 30.82 | 23.08 | 21.58 | | | |
| E71 | 112.75 | 174.98 | 57.635 | 31.98 | 26.58 | | 181.98 | | |
| T72 | 118.75 | 174.03 | 66.415 | 67.58 | | | | | |
| A73 | 124.35 | 175.73 | 54.6 | 18.5 | | | | | |
| T74 | 97.82 | 175.93 | 60.715 | 69.2 | 21.01 | | | | |
| T75 | 110.15 | 172.08 | 62.62 | 68.76 | 21.15 | | | | |
| V76 | 121.2 | 178.38 | 65.66 | 31.3 | 22.68 | 19.78 | | | |
| G77 | 100.3 | 173.88 | 48.2 | | | | | | |
| Y78 | 115.4 | 177.58 | 61.045 | 37.86 | | | | | |
| G79 | 101.33 | 173.73 | 45 | | | | | | |
| D80 | 117.95 | 174.93 | 55.045 | 36.92 | 178.98 | | | | |
| L81 | 117 | 174.93 | 52.335 | 47.22 | 27.08 | | | | |
| Y82 | 114.95 | 171.48 | 54.8 | 34.35 | | | | | |

| **Residue** | **N** | **C** | **C$\alpha$** | **C$\beta$** | **C$\gamma 1$** | **C$\gamma 2$** | **C$\delta$** | **C$\varepsilon$** | **C$\zeta$** |
|---|---|---|---|---|---|---|---|---|---|
| P83 | 131.7 | 175.88 | 58.75 | 31.98 | | | 48.79 | | |
| V84 | 117.8 | 175.73 | 60.115 | 32.04 | 20.78 | 17.78 | | | |
| T85 | 115.95 | 174.28 | 60.055 | 71.95 | | 22.38 | | | |
| L86 | 122.15 | 177.53 | 57.87 | 40.79 | | | | | |
| W87 | | 178.13 | 58.595 | 29.11 | | | | | |
| G88 | 106.85 | 175.38 | 45.825 | | | | | | |
| R89 | 121.35 | 177.58 | 58.89 | 29.28 | | | | | |
| L90 | 118.25 | 178.58 | 58.085 | 42.36 | | | | | |
| V91 | 117.65 | 177.28 | 66.8 | 30.96 | | | | | |
| A92 | 119.4 | 178.28 | 55.37 | 20.11 | | | | | |
| V93 | 117.25 | 176.88 | 66.43 | | | | | | |
| V94 | 118.1 | 177.03 | 66.7 | 30.92 | | | | | |
| V95 | 118.5 | 177.03 | 66.795 | 30.85 | | | | | |
| M96 | 116.05 | 176.68 | 59.305 | 33.18 | 32.48 | | | | |
| V97 | 115.95 | 178.48 | 66.085 | 31.05 | | | | | |
| A98 | 124.85 | 179.73 | 54.93 | 17.91 | | | | | |
| G99 | 107.05 | 174.08 | 47.28 | | | | | | |
| I100 | 120.8 | 177.13 | 65.5 | 38.79 | 25.48 | 17.78 | 13.78 | | |
| T101 | 120 | 176.88 | 67.1 | 67.71 | | | | | |
| S102 | 116.6 | 175.08 | 62.825 | 62.07 | | | | | |
| F103 | 120.25 | 178.68 | 60.175 | 37.07 | | | | | |
| G104 | 108 | 174.53 | 47.34 | | | | | | |
| L105 | 119.4 | 178.73 | 58.11 | 41.13 | | | | | |
| V106 | 117.1 | 177.08 | 66.705 | 30.95 | | | | | |
| T107 | 117.15 | 175.53 | 68.19 | 66.89 | | | | | |
| A108 | 121.05 | 177.58 | 54.78 | 17.3 | | | | | |
| A109 | 118.3 | 178.08 | 54.51 | 17.81 | | | | | |
| L110 | 119.2 | 177.58 | 56.76 | 41.37 | | | | | |
| A111 | | | | | | | | | |

| **Residue** | **N** | **C** | **C$\alpha$** | **C$\beta$** | **C$\gamma$1** | **C$\gamma$2** | **C$\delta$** | **C$\varepsilon$** | **C$\zeta$** |
|---|---|---|---|---|---|---|---|---|---|
| T112 | | | | | | | | | |
| W113 | | | | | | | | | |
| F114 | | | | | | | | | |
| V115 | 114.5 | 178.58 | 65.98 | 31.22 | | | | | |
| G116 | 109.8 | 174.58 | 47.065 | | | | | | |
| R117 | 120.6 | 174.38 | 58.34 | | | | | | |
| E118 | 116.6 | | 58.66 | 29.08 | 35.48 | | 182.28 | | |
| Q119 | | | | | | | | | |
| E120 | | | 58.96 | 27.48 | 35.68 | | 182.28 | | |

TABLE S6: Chemical shift differences between the activated and deactivated states ($\Delta CS_{exp} = CS_{exp}^{pH=4,act} - CS_{exp}^{pH=7.5,deact}$)

| **Residue** | **N** | **C** | **C$\alpha$** | **C$\beta$** |
|---|---|---|---|---|
| T33 | 1.37 | -0.29 | -0.22 | 0.03 |
| V34 | 0.45 | -0.46 | 0.4 | 0.06 |
| L35 | | | | |
| L36 | | | | |
| V37 | | | | |
| I38 | -0.27 | 0.19 | -0.1 | -0.15 |
| V39 | | | | |
| L40 | | | | |
| L41 | | | | |
| A42 | 0.48 | -0.18 | -0.03 | 0.02 |
| G43 | -0.09 | 0.06 | 0.02 | |
| S44 | 0.09 | 0.53 | -0.04 | 0.02 |
| Y45 | -0.02 | 0.53 | -0.17 | 0.01 |
| L46 | | | | |
| A47 | 0.67 | -0.05 | 0.14 | 0.01 |

| Residue | N | C | $C_\alpha$ | $C_\beta$ |
|---|---|---|---|---|
| V48 | -0.07 | -0.02 | 0.01 | -0.21 |
| L49 | | | | |
| A50 | -1.68 | -0.22 | -0.19 | |
| E51 | 0.01 | -0.26 | -0.16 | 0.12 |
| R52 | | | | |
| G53 | 0.26 | 0.33 | 0.22 | |
| A54 | | | | |
| P55 | -0.26 | | 0.2 | |
| G56 | -0.13 | 0.03 | 0.18 | |
| A57 | -0.11 | -0.2 | -0.16 | 0 |
| Q58 | -0.19 | -0.23 | -0.18 | |
| L59 | 0.24 | -0.04 | -0.18 | -0.14 |
| I60 | -0.12 | -0.29 | 0 | -0.16 |
| T61 | 0.21 | -0.17 | -0.17 | 0.01 |
| Y62 | -0.02 | -0.07 | 0.02 | 0.12 |
| P63 | 0.14 | -0.03 | 0.22 | |
| R64 | 0.22 | -0.12 | -0.05 | -0.16 |
| A65 | 0.34 | -0.18 | -0.04 | -0.08 |
| L66 | 0.1 | 0.2 | -0.31 | 0.15 |
| W67 | 0.26 | 0.19 | -0.14 | 0.01 |
| W68 | 0.02 | -0.03 | 0.02 | 0.12 |
| S69 | -0.09 | -0.13 | -0.07 | 0.04 |
| V70 | -0.12 | -0.15 | -0.07 | 0.06 |
| E71 | 0.13 | 0.09 | 0 | -0.07 |
| T72 | 0.05 | -0.07 | 0.03 | -0.02 |
| A73 | -0.12 | 0.23 | 0.14 | -0.05 |
| T74 | 1.09 | -0.22 | -0.1 | -0.02 |
| T75 | -0.02 | -0.11 | -0.02 | -0.08 |
| V76 | -0.14 | -0.16 | 0.08 | 0.04 |

| Residue | N | C | $C_\alpha$ | $C_\beta$ |
|---|---|---|---|---|
| G77 | 0.03 | -0.13 | -0.1 | |
| Y78 | -0.1 | -0.09 | -0.01 | -0.05 |
| G79 | -0.22 | 0.09 | -0.09 | |
| D80 | -0.1 | 0.14 | -0.14 | -0.17 |
| L81 | 0.33 | -0.08 | -0.19 | -0.87 |
| Y82 | 0.24 | -0.13 | 0.1 | 0.09 |
| P83 | 0.09 | -0.28 | 2.07 | |
| V84 | 0.23 | -0.02 | 0.03 | -0.13 |
| T85 | 0.14 | -0.23 | 0.05 | 0.19 |
| L86 | 0.01 | -0.06 | 0.1 | |
| W87 | | -0.17 | 0.17 | -0.06 |
| G88 | -0.1 | 0.27 | -0.01 | |
| R89 | 0.15 | -0.14 | -0.17 | -0.06 |
| L90 | 0.77 | 0.39 | 0.08 | -2.1 |
| V91 | 0.08 | 0.27 | -0.02 | -0.21 |
| A92 | 0.08 | -0.03 | -0.05 | -0.08 |
| V93 | -0.23 | -0.04 | 0.11 | |
| V94 | 0.24 | 0.11 | -0.07 | -0.06 |
| V95 | 0.58 | -0.08 | 0.06 | -0.09 |
| M96 | 0.32 | 0.16 | 0.04 | 0.15 |
| V97 | 0.12 | -0.03 | 0.25 | -0.04 |
| A98 | -0.08 | 0.6 | -0.11 | 0.57 |
| G99 | -0.19 | 0.24 | -0.11 | |
| I100 | | | | |
| T101 | | | | |
| S102 | -0.87 | -0.22 | 0.11 | -0.05 |
| F103 | -0.05 | | | |
| G104 | | | | |
| L105 | | | | |

| Residue | N | C | Cα | Cβ |
|---|---|---|---|---|
| V106 | 0.01 | -0.23 | -0.33 | 0 |
| T107 | -0.88 | 0.1 | -0.42 | 0.21 |
| A108 | 0.48 | 1.97 | 0.27 | 0.23 |
| A109 | 5.64 | -0.37 | 0.34 | -0.84 |
| L110 | 0.03 | 0.54 | 0.32 | 0.58 |
| A111 | 0.11 | -0.32 | 0.09 | 0.2 |
| T112 | -0.41 | -0.16 | 0.11 | 0.27 |

TABLE S7: Simulated chemical shifts for $C_\alpha$ for the two chemical shift prediction methods and the three channel states. The data represented corresponds to the credible intervals of the chemical shifts that are inferred from the simulation data ($CS^{X}_{sim}$). The credible intervals are expressed as the center of the interval plus or minus the distance to the upper and lower bounds of the interval. This interval measures our uncertainty of the resonance position in the spectrum and not the peak width which is related to the standard deviation of the skew-normal distribution with which the data is modeled.

| Residue | Fully Open, SPARTA+ | Partially Open, SPARTA+ | Closed, SPARTA+ | Fully Open, SHIFTX2 | Partially Open, SHIFTX2 | Closed, SHIFTX2 |
|---|---|---|---|---|---|---|
| I38 | 65.01 ± 0.03 | 64.96 ± 0.03 | 64.84 ± 0.03 | 64.21 ± 0.02 | 64.16 ± 0.02 | 64.09 ± 0.02 |
| A42 | 55.32 ± 0.01 | 55.29 ± 0.01 | 55.27 ± 0.01 | 55.14 ± 0.02 | 55.12 ± 0.01 | 55.13 ± 0.01 |
| G43 | 47.15 ± 0.01 | 47.14 ± 0.01 | 47.12 ± 0.01 | 47.63 ± 0.01 | 47.61 ± 0.01 | 47.56 ± 0.02 |
| S44 | 62.23 ± 0.02 | 62.17 ± 0.02 | 62.51 ± 0.02 | 62.00 ± 0.02 | 61.89 ± 0.02 | 62.09 ± 0.02 |
| Y45 | 61.72 ± 0.02 | 61.46 ± 0.02 | 61.49 ± 0.02 | 60.95 ± 0.02 | 60.80 ± 0.02 | 60.88 ± 0.02 |
| A47 | 55.14 ± 0.01 | 55.17 ± 0.01 | 55.16 ± 0.01 | 55.14 ± 0.01 | 55.16 ± 0.01 | 55.14 ± 0.01 |
| V48 | 66.06 ± 0.02 | 66.26 ± 0.02 | 66.22 ± 0.02 | 65.91 ± 0.02 | 66.16 ± 0.02 | 66.25 ± 0.02 |
| A50 | 54.27 ± 0.02 | 54.55 ± 0.01 | 54.64 ± 0.01 | 54.47 ± 0.02 | 54.62 ± 0.01 | 54.73 ± 0.01 |
| E51 | 56.39 ± 0.03 | 56.85 ± 0.04 | 57.70 ± 0.04 | 57.19 ± 0.03 | 57.40 ± 0.04 | 57.98 ± 0.04 |
| G53 | 44.90 ± 0.01 | 44.83 ± 0.01 | 44.92 ± 0.02 | 45.52 ± 0.02 | 45.60 ± 0.02 | 45.70 ± 0.03 |
| G56 | 44.57 ± 0.01 | 44.75 ± 0.01 | 44.77 ± 0.01 | 45.19 ± 0.02 | 45.44 ± 0.02 | 45.47 ± 0.02 |
| A57 | 52.32 ± 0.02 | 52.38 ± 0.02 | 52.35 ± 0.02 | 52.26 ± 0.02 | 52.46 ± 0.02 | 52.44 ± 0.02 |
| Q58 | 57.23 ± 0.04 | 56.59 ± 0.04 | 56.65 ± 0.03 | 56.77 ± 0.06 | 56.16 ± 0.05 | 56.29 ± 0.05 |
| L59 | 54.27 ± 0.02 | 54.53 ± 0.02 | 54.50 ± 0.02 | 54.18 ± 0.02 | 54.32 ± 0.03 | 54.21 ± 0.02 |

| Residue | Fully Open, SPARTA+ | Partially Open, SPARTA+ | Closed, SPARTA+ | Fully Open, SHIFTX2 | Partially Open, SHIFTX2 | Closed, SHIFTX2 |
|---|---|---|---|---|---|---|
| I60 | 60.79 ± 0.04 | 60.77 ± 0.03 | 60.84 ± 0.03 | 61.37 ± 0.03 | 61.01 ± 0.04 | 61.27 ± 0.03 |
| T61 | 59.87 ± 0.02 | 60.72 ± 0.04 | 60.08 ± 0.02 | 59.79 ± 0.02 | 60.71 ± 0.04 | 59.85 ± 0.02 |
| Y62 | 63.06 ± 0.03 | 62.75 ± 0.03 | 62.54 ± 0.03 | 61.91 ± 0.02 | 61.76 ± 0.03 | 61.67 ± 0.02 |
| R64 | 59.62 ± 0.01 | 59.24 ± 0.02 | 59.41 ± 0.02 | 59.88 ± 0.02 | 59.37 ± 0.04 | 59.72 ± 0.02 |
| A65 | 55.15 ± 0.01 | 55.15 ± 0.01 | 55.18 ± 0.01 | 55.37 ± 0.01 | 55.40 ± 0.01 | 55.45 ± 0.01 |
| L66 | 58.05 ± 0.01 | 57.97 ± 0.01 | 57.92 ± 0.01 | 57.97 ± 0.01 | 57.89 ± 0.01 | 57.91 ± 0.02 |
| W67 | 60.16 ± 0.02 | 60.22 ± 0.02 | 60.29 ± 0.02 | 60.45 ± 0.02 | 60.51 ± 0.01 | 60.49 ± 0.01 |
| W68 | 60.86 ± 0.02 | 60.74 ± 0.02 | 60.57 ± 0.02 | 60.87 ± 0.02 | 60.83 ± 0.02 | 60.80 ± 0.02 |
| S69 | 62.48 ± 0.01 | 62.38 ± 0.02 | 62.22 ± 0.02 | 62.43 ± 0.01 | 62.41 ± 0.02 | 62.20 ± 0.02 |
| V70 | 66.42 ± 0.02 | 66.24 ± 0.03 | 66.35 ± 0.02 | 66.46 ± 0.02 | 66.38 ± 0.02 | 66.48 ± 0.02 |
| E71 | 58.87 ± 0.02 | 58.81 ± 0.02 | 58.69 ± 0.02 | 59.43 ± 0.03 | 59.24 ± 0.03 | 59.04 ± 0.03 |
| T72 | 65.83 ± 0.01 | 65.96 ± 0.01 | 65.98 ± 0.02 | 66.21 ± 0.02 | 66.17 ± 0.03 | 66.20 ± 0.03 |
| A73 | 54.47 ± 0.01 | 54.42 ± 0.01 | 54.40 ± 0.01 | 54.59 ± 0.02 | 54.65 ± 0.02 | 54.62 ± 0.02 |
| T74 | 63.10 ± 0.04 | 63.35 ± 0.04 | 63.47 ± 0.04 | 62.18 ± 0.04 | 62.59 ± 0.04 | 62.58 ± 0.04 |
| T75 | 63.62 ± 0.01 | 63.27 ± 0.01 | 63.38 ± 0.01 | 63.77 ± 0.02 | 63.60 ± 0.03 | 63.58 ± 0.03 |
| V76 | 64.43 ± 0.03 | 64.66 ± 0.03 | 64.70 ± 0.03 | 64.65 ± 0.02 | 64.80 ± 0.02 | 64.84 ± 0.02 |
| G77 | 45.75 ± 0.01 | 45.81 ± 0.01 | 45.80 ± 0.01 | 46.09 ± 0.01 | 46.16 ± 0.01 | 46.20 ± 0.01 |
| Y78 | 59.48 ± 0.03 | 60.15 ± 0.02 | 60.13 ± 0.02 | 59.98 ± 0.02 | 60.70 ± 0.01 | 60.68 ± 0.01 |
| G79 | 45.33 ± 0.01 | 45.28 ± 0.00 | 45.31 ± 0.01 | 45.55 ± 0.01 | 45.59 ± 0.01 | 45.62 ± 0.01 |
| D80 | 55.47 ± 0.01 | 55.53 ± 0.01 | 55.53 ± 0.01 | 56.24 ± 0.02 | 56.30 ± 0.02 | 56.12 ± 0.02 |
| L81 | 54.05 ± 0.01 | 54.14 ± 0.02 | 54.26 ± 0.02 | 53.76 ± 0.01 | 53.82 ± 0.02 | 53.84 ± 0.02 |
| Y82 | 54.33 ± 0.02 | 54.34 ± 0.02 | 54.28 ± 0.02 | 55.17 ± 0.02 | 55.21 ± 0.02 | 55.17 ± 0.02 |
| V84 | 61.79 ± 0.03 | 61.65 ± 0.03 | 61.64 ± 0.02 | 62.32 ± 0.04 | 62.18 ± 0.03 | 62.19 ± 0.03 |
| T85 | 61.24 ± 0.04 | 61.37 ± 0.04 | 61.75 ± 0.04 | 60.47 ± 0.04 | 60.59 ± 0.04 | 60.91 ± 0.04 |
| L86 | 57.82 ± 0.02 | 57.97 ± 0.02 | 58.13 ± 0.01 | 58.25 ± 0.02 | 58.21 ± 0.01 | 58.29 ± 0.01 |
| W87 | 59.15 ± 0.02 | 59.07 ± 0.02 | 59.10 ± 0.03 | 60.38 ± 0.02 | 60.17 ± 0.02 | 60.11 ± 0.03 |
| G88 | 46.61 ± 0.01 | 46.65 ± 0.01 | 46.63 ± 0.01 | 47.41 ± 0.01 | 47.43 ± 0.01 | 47.42 ± 0.01 |
| R89 | 58.87 ± 0.01 | 59.08 ± 0.01 | 59.11 ± 0.01 | 59.18 ± 0.02 | 59.51 ± 0.02 | 59.53 ± 0.02 |

| Residue | Fully Open, SPARTA+ | Partially Open, SPARTA+ | Closed, SPARTA+ | Fully Open, SHIFTX2 | Partially Open, SHIFTX2 | Closed, SHIFTX2 |
|---|---|---|---|---|---|---|
| L90 | 57.80 ± 0.01 | 57.87 ± 0.01 | 57.88 ± 0.01 | 57.86 ± 0.02 | 57.93 ± 0.01 | 57.94 ± 0.02 |
| V91 | 66.49 ± 0.02 | 66.49 ± 0.02 | 66.43 ± 0.02 | 66.62 ± 0.01 | 66.56 ± 0.02 | 66.49 ± 0.02 |
| A92 | 55.67 ± 0.01 | 55.62 ± 0.01 | 55.62 ± 0.01 | 55.41 ± 0.01 | 55.39 ± 0.01 | 55.40 ± 0.01 |
| V93 | 66.26 ± 0.02 | 66.75 ± 0.02 | 66.52 ± 0.03 | 66.07 ± 0.02 | 66.23 ± 0.01 | 66.15 ± 0.02 |
| V94 | 66.54 ± 0.02 | 66.59 ± 0.02 | 66.58 ± 0.02 | 65.98 ± 0.01 | 66.01 ± 0.01 | 66.02 ± 0.01 |
| V95 | 66.72 ± 0.02 | 66.74 ± 0.01 | 66.77 ± 0.02 | 66.05 ± 0.01 | 66.11 ± 0.01 | 66.15 ± 0.01 |
| M96 | 59.80 ± 0.01 | 59.10 ± 0.01 | 59.16 ± 0.01 | 59.26 ± 0.01 | 58.92 ± 0.02 | 59.01 ± 0.02 |
| V97 | 66.53 ± 0.02 | 66.31 ± 0.02 | 66.16 ± 0.02 | 66.12 ± 0.02 | 66.12 ± 0.02 | 66.15 ± 0.02 |
| A98 | 55.28 ± 0.02 | 55.05 ± 0.02 | 55.31 ± 0.01 | 54.62 ± 0.02 | 54.72 ± 0.02 | 54.97 ± 0.01 |
| G99 | 46.78 ± 0.01 | 46.93 ± 0.01 | 46.86 ± 0.01 | 47.01 ± 0.02 | 47.31 ± 0.01 | 47.21 ± 0.01 |
| S102 | 61.73 ± 0.01 | 62.17 ± 0.01 | 62.16 ± 0.01 | 61.93 ± 0.02 | 62.29 ± 0.02 | 62.33 ± 0.02 |
| V106 | 66.83 ± 0.02 | 66.88 ± 0.02 | 67.02 ± 0.02 | 66.44 ± 0.01 | 66.45 ± 0.02 | 66.58 ± 0.02 |
| T107 | 65.67 ± 0.02 | 66.94 ± 0.02 | 67.15 ± 0.01 | 66.01 ± 0.03 | 66.86 ± 0.03 | 67.29 ± 0.01 |
| A108 | 54.98 ± 0.01 | 55.14 ± 0.01 | 55.23 ± 0.01 | 55.05 ± 0.01 | 55.14 ± 0.02 | 55.24 ± 0.01 |
| A109 | 54.97 ± 0.01 | 54.98 ± 0.01 | 54.73 ± 0.01 | 55.11 ± 0.01 | 55.22 ± 0.01 | 55.13 ± 0.01 |
| L110 | 57.76 ± 0.01 | 57.69 ± 0.01 | 57.49 ± 0.01 | 58.12 ± 0.02 | 58.10 ± 0.02 | 57.91 ± 0.01 |

TABLE S8: Simulated chemical shifts for $C_\beta$ for the two chemical shift prediction methods and the three channel states. The data represented corresponds to the credible intervals of the chemical shifts that are inferred from the simulation data ($CS^{X}_{sim}$). The credible intervals are expressed as the center of the interval plus or minus the distance to the upper and lower bounds of the interval. This interval measures our uncertainty of the resonance position in the spectrum and not the peak width which is related to the standard deviation of the skew-normal distribution with which the data is modeled.

| Residue | Fully Open, SPARTA+ | Partially Open, SPARTA+ | Closed, SPARTA+ | Fully Open, SHIFTX2 | Partially Open, SHIFTX2 | Closed, SHIFTX2 |
|---|---|---|---|---|---|---|
| I38 | 37.19 ± 0.02 | 37.14 ± 0.02 | 37.07 ± 0.03 | 37.79 ± 0.01 | 37.74 ± 0.01 | 37.69 ± 0.01 |
| A42 | 18.15 ± 0.00 | 18.16 ± 0.01 | 18.15 ± 0.00 | 18.29 ± 0.01 | 18.33 ± 0.01 | 18.29 ± 0.01 |
| S44 | 63.27 ± 0.01 | 63.20 ± 0.01 | 63.30 ± 0.01 | 62.97 ± 0.01 | 62.95 ± 0.01 | 62.94 ± 0.01 |
| Y45 | 38.75 ± 0.01 | 38.71 ± 0.01 | 38.66 ± 0.01 | 38.32 ± 0.01 | 38.30 ± 0.01 | 38.32 ± 0.01 |

| Residue | Fully Open, SPARTA+ | Partially Open, SPARTA+ | Closed, SPARTA+ | Fully Open, SHIFTX2 | Partially Open, SHIFTX2 | Closed, SHIFTX2 |
|---|---|---|---|---|---|---|
| A47 | 18.18 ± 0.01 | 18.21 ± 0.01 | 18.23 ± 0.01 | 18.39 ± 0.01 | 18.41 ± 0.01 | 18.40 ± 0.01 |
| V48 | 31.33 ± 0.01 | 31.48 ± 0.01 | 31.48 ± 0.01 | 31.45 ± 0.01 | 31.51 ± 0.01 | 31.52 ± 0.00 |
| E51 | 30.09 ± 0.02 | 30.11 ± 0.02 | 29.86 ± 0.02 | 29.90 ± 0.02 | 29.92 ± 0.02 | 29.67 ± 0.02 |
| A57 | 19.68 ± 0.02 | 19.51 ± 0.02 | 19.48 ± 0.02 | 19.38 ± 0.02 | 19.26 ± 0.02 | 19.27 ± 0.02 |
| L59 | 41.63 ± 0.03 | 41.52 ± 0.04 | 41.35 ± 0.05 | 41.86 ± 0.03 | 41.77 ± 0.04 | 41.87 ± 0.05 |
| I60 | 37.70 ± 0.03 | 38.70 ± 0.06 | 38.74 ± 0.05 | 38.13 ± 0.02 | 38.96 ± 0.05 | 39.08 ± 0.04 |
| T61 | 71.49 ± 0.02 | 70.57 ± 0.05 | 71.30 ± 0.02 | 71.28 ± 0.02 | 70.77 ± 0.04 | 71.28 ± 0.02 |
| Y62 | 35.71 ± 0.01 | 36.21 ± 0.02 | 35.97 ± 0.01 | 36.79 ± 0.01 | 36.78 ± 0.01 | 36.72 ± 0.01 |
| R64 | 29.77 ± 0.01 | 29.59 ± 0.01 | 29.77 ± 0.01 | 29.98 ± 0.01 | 29.72 ± 0.02 | 29.95 ± 0.01 |
| A65 | 18.45 ± 0.01 | 18.43 ± 0.01 | 18.45 ± 0.01 | 18.37 ± 0.01 | 18.38 ± 0.01 | 18.40 ± 0.01 |
| L66 | 41.96 ± 0.01 | 41.92 ± 0.01 | 41.98 ± 0.01 | 41.73 ± 0.01 | 41.73 ± 0.01 | 41.77 ± 0.01 |
| W67 | 29.75 ± 0.01 | 29.57 ± 0.01 | 29.54 ± 0.01 | 29.57 ± 0.01 | 29.46 ± 0.01 | 29.41 ± 0.01 |
| W68 | 29.15 ± 0.01 | 29.24 ± 0.01 | 29.35 ± 0.01 | 28.95 ± 0.01 | 28.99 ± 0.01 | 28.97 ± 0.01 |
| S69 | 62.72 ± 0.01 | 62.72 ± 0.01 | 62.70 ± 0.01 | 62.85 ± 0.01 | 62.94 ± 0.01 | 62.91 ± 0.01 |
| V70 | 31.31 ± 0.01 | 31.41 ± 0.01 | 31.39 ± 0.01 | 31.60 ± 0.01 | 31.66 ± 0.01 | 31.64 ± 0.01 |
| E71 | 28.98 ± 0.01 | 28.82 ± 0.01 | 28.73 ± 0.01 | 28.95 ± 0.01 | 28.95 ± 0.01 | 28.86 ± 0.01 |
| T72 | 68.29 ± 0.01 | 68.28 ± 0.01 | 68.32 ± 0.01 | 68.35 ± 0.01 | 68.39 ± 0.02 | 68.46 ± 0.02 |
| A73 | 18.50 ± 0.01 | 18.64 ± 0.01 | 18.56 ± 0.01 | 18.59 ± 0.01 | 18.59 ± 0.01 | 18.59 ± 0.01 |
| T74 | 69.11 ± 0.02 | 69.27 ± 0.01 | 69.30 ± 0.01 | 69.39 ± 0.02 | 69.37 ± 0.02 | 69.45 ± 0.01 |
| T75 | 66.95 ± 0.02 | 67.02 ± 0.02 | 67.09 ± 0.02 | 68.23 ± 0.01 | 68.15 ± 0.01 | 68.19 ± 0.01 |
| V76 | 31.69 ± 0.01 | 31.61 ± 0.01 | 31.65 ± 0.01 | 32.12 ± 0.01 | 32.09 ± 0.01 | 32.10 ± 0.01 |
| Y78 | 38.73 ± 0.01 | 38.92 ± 0.01 | 38.92 ± 0.01 | 38.25 ± 0.01 | 38.83 ± 0.01 | 38.86 ± 0.01 |
| D80 | 40.71 ± 0.01 | 40.71 ± 0.01 | 40.70 ± 0.01 | 40.63 ± 0.01 | 40.58 ± 0.01 | 40.41 ± 0.02 |
| L81 | 44.69 ± 0.02 | 44.89 ± 0.02 | 44.80 ± 0.02 | 44.70 ± 0.03 | 44.87 ± 0.03 | 44.90 ± 0.03 |
| Y82 | 38.52 ± 0.02 | 38.53 ± 0.02 | 38.57 ± 0.02 | 39.98 ± 0.02 | 40.05 ± 0.02 | 40.12 ± 0.02 |
| V84 | 32.97 ± 0.03 | 33.14 ± 0.03 | 33.17 ± 0.03 | 33.47 ± 0.02 | 33.61 ± 0.02 | 33.61 ± 0.02 |
| T85 | 71.12 ± 0.03 | 71.04 ± 0.03 | 70.83 ± 0.03 | 70.87 ± 0.03 | 70.70 ± 0.03 | 70.53 ± 0.03 |
| W87 | 28.75 ± 0.01 | 28.89 ± 0.01 | 28.90 ± 0.01 | 28.78 ± 0.01 | 28.83 ± 0.01 | 28.84 ± 0.01 |

| Residue | Fully Open, SPARTA+ | Partially Open, SPARTA+ | Closed, SPARTA+ | Fully Open, SHIFTX2 | Partially Open, SHIFTX2 | Closed, SHIFTX2 |
|---|---|---|---|---|---|---|
| R89 | 29.47 ± 0.01 | 29.56 ± 0.01 | 29.58 ± 0.01 | 29.44 ± 0.01 | 29.56 ± 0.01 | 29.56 ± 0.01 |
| L90 | 41.74 ± 0.01 | 41.75 ± 0.01 | 41.76 ± 0.01 | 41.63 ± 0.01 | 41.65 ± 0.01 | 41.69 ± 0.01 |
| V91 | 31.27 ± 0.02 | 31.32 ± 0.01 | 31.33 ± 0.01 | 31.55 ± 0.01 | 31.59 ± 0.01 | 31.61 ± 0.01 |
| A92 | 18.63 ± 0.01 | 18.68 ± 0.00 | 18.71 ± 0.00 | 18.56 ± 0.01 | 18.67 ± 0.01 | 18.66 ± 0.01 |
| V94 | 31.27 ± 0.02 | 31.26 ± 0.01 | 31.22 ± 0.01 | 31.75 ± 0.01 | 31.72 ± 0.00 | 31.70 ± 0.00 |
| V95 | 31.38 ± 0.01 | 31.33 ± 0.01 | 31.31 ± 0.01 | 31.49 ± 0.01 | 31.48 ± 0.01 | 31.46 ± 0.01 |
| M96 | 32.42 ± 0.01 | 32.21 ± 0.01 | 32.23 ± 0.01 | 33.01 ± 0.01 | 32.54 ± 0.01 | 32.57 ± 0.01 |
| V97 | 31.39 ± 0.02 | 31.12 ± 0.02 | 30.98 ± 0.02 | 31.76 ± 0.01 | 31.68 ± 0.01 | 31.61 ± 0.01 |
| A98 | 18.37 ± 0.01 | 18.29 ± 0.01 | 18.16 ± 0.01 | 18.71 ± 0.02 | 18.30 ± 0.01 | 18.26 ± 0.01 |
| S102 | 62.86 ± 0.01 | 62.74 ± 0.01 | 62.80 ± 0.01 | 62.86 ± 0.01 | 62.66 ± 0.01 | 62.74 ± 0.01 |
| V106 | 31.33 ± 0.02 | 31.27 ± 0.02 | 31.26 ± 0.01 | 31.56 ± 0.01 | 31.51 ± 0.01 | 31.52 ± 0.01 |
| T107 | 68.13 ± 0.01 | 68.38 ± 0.01 | 68.27 ± 0.01 | 68.47 ± 0.01 | 68.58 ± 0.01 | 68.42 ± 0.01 |
| A108 | 18.08 ± 0.00 | 18.16 ± 0.00 | 18.22 ± 0.01 | 18.21 ± 0.01 | 18.35 ± 0.01 | 18.34 ± 0.01 |
| A109 | 18.31 ± 0.01 | 18.51 ± 0.01 | 18.06 ± 0.02 | 18.30 ± 0.01 | 18.38 ± 0.01 | 18.26 ± 0.01 |
| L110 | 41.62 ± 0.01 | 41.61 ± 0.01 | 41.38 ± 0.01 | 41.54 ± 0.01 | 41.57 ± 0.01 | 41.47 ± 0.01 |

TABLE S9: Simulated chemical shifts for C for the two chemical shift prediction methods and the three channel states. The data represented corresponds to the credible intervals of the chemical shifts that are inferred from the simulation data ($CS^{X}_{sim}$). The credible intervals are expressed as the center of the interval plus or minus the distance to the upper and lower bounds of the interval. This interval measures our uncertainty of the resonance position in the spectrum and not the peak width which is related to the standard deviation of the skew-normal distribution with which the data is modeled.

| Residue | Fully Open, SPARTA+ | Partially Open, SPARTA+ | Closed, SPARTA+ | Fully Open, SHIFTX2 | Partially Open, SHIFTX2 | Closed, SHIFTX2 |
|---|---|---|---|---|---|---|
| I38 | 178.17 ± 0.01 | 178.12 ± 0.01 | 178.16 ± 0.01 | 178.48 ± 0.01 | 178.44 ± 0.01 | 178.48 ± 0.01 |
| A42 | 179.86 ± 0.00 | 179.83 ± 0.00 | 179.84 ± 0.00 | 180.15 ± 0.01 | 180.15 ± 0.01 | 180.15 ± 0.01 |
| G43 | 176.50 ± 0.01 | 176.53 ± 0.01 | 176.49 ± 0.01 | 175.99 ± 0.02 | 175.99 ± 0.02 | 176.18 ± 0.02 |
| S44 | 176.21 ± 0.01 | 176.27 ± 0.01 | 176.23 ± 0.01 | 176.22 ± 0.01 | 176.26 ± 0.01 | 176.17 ± 0.01 |
| Y45 | 177.51 ± 0.01 | 177.45 ± 0.01 | 177.29 ± 0.01 | 178.24 ± 0.02 | 178.10 ± 0.02 | 178.09 ± 0.02 |

| Residue | Fully Open, SPARTA+ | Partially Open, SPARTA+ | Closed, SPARTA+ | Fully Open, SHIFTX2 | Partially Open, SHIFTX2 | Closed, SHIFTX2 |
|---|---|---|---|---|---|---|
| A47 | 179.33 ± 0.01 | 179.30 ± 0.01 | 179.31 ± 0.01 | 178.89 ± 0.01 | 178.86 ± 0.01 | 178.91 ± 0.01 |
| V48 | 178.17 ± 0.01 | 178.13 ± 0.01 | 178.15 ± 0.01 | 177.78 ± 0.02 | 177.91 ± 0.02 | 177.81 ± 0.02 |
| A50 | 179.07 ± 0.02 | 179.45 ± 0.02 | 179.70 ± 0.02 | 178.67 ± 0.03 | 179.26 ± 0.02 | 179.61 ± 0.02 |
| E51 | 176.61 ± 0.03 | 177.35 ± 0.04 | 178.11 ± 0.05 | 176.28 ± 0.02 | 176.70 ± 0.04 | 177.36 ± 0.04 |
| G53 | 173.82 ± 0.02 | 173.91 ± 0.02 | 173.99 ± 0.02 | 173.92 ± 0.01 | 174.03 ± 0.02 | 174.13 ± 0.02 |
| G56 | 173.87 ± 0.02 | 174.15 ± 0.02 | 174.16 ± 0.01 | 174.36 ± 0.01 | 174.44 ± 0.01 | 174.43 ± 0.01 |
| A57 | 177.53 ± 0.03 | 177.30 ± 0.02 | 177.39 ± 0.03 | 177.56 ± 0.02 | 177.57 ± 0.02 | 177.48 ± 0.01 |
| Q58 | 176.56 ± 0.02 | 176.18 ± 0.03 | 176.03 ± 0.03 | 176.20 ± 0.02 | 175.86 ± 0.03 | 175.89 ± 0.02 |
| L59 | 176.75 ± 0.03 | 176.14 ± 0.04 | 175.91 ± 0.03 | 176.47 ± 0.01 | 176.16 ± 0.03 | 176.29 ± 0.02 |
| I60 | 174.91 ± 0.02 | 174.95 ± 0.02 | 174.85 ± 0.02 | 175.14 ± 0.01 | 175.03 ± 0.03 | 175.01 ± 0.03 |
| T61 | 174.87 ± 0.02 | 174.81 ± 0.02 | 175.00 ± 0.02 | 174.95 ± 0.02 | 175.04 ± 0.01 | 174.96 ± 0.02 |
| Y62 | 176.18 ± 0.01 | 175.98 ± 0.02 | 176.04 ± 0.01 | 176.39 ± 0.05 | 176.10 ± 0.05 | 176.24 ± 0.04 |
| P63 | 178.99 ± 0.01 | 178.93 ± 0.01 | 178.83 ± 0.01 | 179.35 ± 0.02 | 179.41 ± 0.02 | 179.30 ± 0.02 |
| R64 | 178.58 ± 0.01 | 178.59 ± 0.01 | 178.49 ± 0.01 | 178.49 ± 0.01 | 178.54 ± 0.01 | 178.48 ± 0.01 |
| A65 | 179.83 ± 0.01 | 179.89 ± 0.01 | 179.85 ± 0.01 | 179.65 ± 0.01 | 179.61 ± 0.01 | 179.48 ± 0.01 |
| L66 | 178.15 ± 0.01 | 178.18 ± 0.01 | 178.35 ± 0.02 | 178.88 ± 0.01 | 178.88 ± 0.01 | 178.96 ± 0.02 |
| W67 | 177.75 ± 0.01 | 177.72 ± 0.01 | 177.67 ± 0.01 | 177.37 ± 0.01 | 177.42 ± 0.02 | 177.41 ± 0.02 |
| W68 | 178.03 ± 0.01 | 178.15 ± 0.01 | 178.27 ± 0.01 | 177.21 ± 0.02 | 177.20 ± 0.02 | 177.20 ± 0.02 |
| S69 | 176.29 ± 0.01 | 176.32 ± 0.01 | 176.37 ± 0.01 | 175.65 ± 0.02 | 175.72 ± 0.02 | 175.73 ± 0.02 |
| V70 | 177.38 ± 0.02 | 177.24 ± 0.02 | 177.37 ± 0.02 | 177.36 ± 0.02 | 177.55 ± 0.02 | 177.54 ± 0.02 |
| E71 | 178.99 ± 0.01 | 178.80 ± 0.01 | 178.78 ± 0.01 | 178.23 ± 0.02 | 178.04 ± 0.02 | 178.03 ± 0.02 |
| T72 | 176.24 ± 0.01 | 176.37 ± 0.01 | 176.38 ± 0.01 | 175.58 ± 0.02 | 175.56 ± 0.02 | 175.58 ± 0.02 |
| A73 | 178.56 ± 0.02 | 178.72 ± 0.03 | 179.16 ± 0.02 | 178.56 ± 0.02 | 178.80 ± 0.02 | 178.71 ± 0.02 |
| T74 | 174.95 ± 0.02 | 175.00 ± 0.02 | 174.94 ± 0.02 | 174.68 ± 0.01 | 174.57 ± 0.01 | 174.73 ± 0.01 |
| T75 | 174.68 ± 0.02 | 174.76 ± 0.02 | 174.77 ± 0.02 | 173.63 ± 0.01 | 173.54 ± 0.01 | 173.57 ± 0.01 |
| V76 | 176.92 ± 0.01 | 176.99 ± 0.01 | 177.06 ± 0.01 | 175.51 ± 0.01 | 175.56 ± 0.01 | 175.59 ± 0.01 |
| G77 | 175.82 ± 0.02 | 175.83 ± 0.01 | 175.84 ± 0.01 | 174.47 ± 0.01 | 174.52 ± 0.01 | 174.55 ± 0.01 |
| Y78 | 176.84 ± 0.01 | 176.83 ± 0.01 | 176.98 ± 0.01 | 176.40 ± 0.01 | 176.46 ± 0.01 | 176.46 ± 0.01 |

| Residue | Fully Open, SPARTA+ | Partially Open, SPARTA+ | Closed, SPARTA+ | Fully Open, SHIFTX2 | Partially Open, SHIFTX2 | Closed, SHIFTX2 |
|---|---|---|---|---|---|---|
| G79 | 174.87 ± 0.01 | 174.98 ± 0.01 | 174.99 ± 0.01 | 173.50 ± 0.01 | 173.56 ± 0.02 | 173.65 ± 0.02 |
| D80 | 176.12 ± 0.01 | 176.10 ± 0.01 | 176.09 ± 0.02 | 175.31 ± 0.01 | 175.33 ± 0.01 | 175.30 ± 0.01 |
| L81 | 174.60 ± 0.02 | 174.52 ± 0.02 | 174.47 ± 0.02 | 174.99 ± 0.02 | 175.05 ± 0.02 | 174.99 ± 0.02 |
| Y82 | 172.89 ± 0.01 | 172.87 ± 0.01 | 172.87 ± 0.01 | 172.38 ± 0.01 | 172.37 ± 0.01 | 172.39 ± 0.01 |
| P83 | 176.00 ± 0.02 | 175.91 ± 0.02 | 175.82 ± 0.02 | 176.03 ± 0.01 | 176.03 ± 0.01 | 175.99 ± 0.01 |
| V84 | 175.80 ± 0.02 | 175.68 ± 0.02 | 175.74 ± 0.02 | 176.03 ± 0.02 | 176.08 ± 0.02 | 176.09 ± 0.02 |
| T85 | 175.33 ± 0.02 | 175.47 ± 0.01 | 175.50 ± 0.01 | 174.85 ± 0.01 | 174.91 ± 0.01 | 174.91 ± 0.01 |
| L86 | 177.59 ± 0.01 | 177.78 ± 0.02 | 177.78 ± 0.01 | 178.52 ± 0.01 | 178.63 ± 0.02 | 178.63 ± 0.02 |
| W87 | 178.51 ± 0.01 | 178.47 ± 0.01 | 178.46 ± 0.01 | 178.82 ± 0.01 | 178.72 ± 0.02 | 178.74 ± 0.02 |
| G88 | 176.01 ± 0.01 | 176.05 ± 0.01 | 176.07 ± 0.01 | 175.83 ± 0.02 | 175.76 ± 0.02 | 175.82 ± 0.02 |
| R89 | 178.48 ± 0.01 | 178.49 ± 0.01 | 178.50 ± 0.01 | 178.21 ± 0.02 | 178.23 ± 0.02 | 178.25 ± 0.02 |
| L90 | 178.91 ± 0.01 | 178.88 ± 0.01 | 178.91 ± 0.01 | 179.82 ± 0.01 | 179.79 ± 0.01 | 179.76 ± 0.01 |
| V91 | 177.84 ± 0.01 | 177.89 ± 0.01 | 177.87 ± 0.01 | 177.71 ± 0.01 | 177.78 ± 0.01 | 177.78 ± 0.01 |
| A92 | 179.74 ± 0.01 | 179.76 ± 0.01 | 179.76 ± 0.01 | 179.20 ± 0.01 | 179.14 ± 0.01 | 179.18 ± 0.01 |
| V93 | 177.93 ± 0.01 | 177.99 ± 0.01 | 177.94 ± 0.01 | 178.01 ± 0.01 | 178.03 ± 0.01 | 177.95 ± 0.02 |
| V94 | 178.17 ± 0.01 | 178.13 ± 0.01 | 178.15 ± 0.01 | 178.22 ± 0.01 | 178.30 ± 0.01 | 178.34 ± 0.01 |
| V95 | 178.19 ± 0.01 | 178.12 ± 0.01 | 178.06 ± 0.01 | 178.06 ± 0.01 | 178.03 ± 0.01 | 177.83 ± 0.01 |
| M96 | 178.92 ± 0.01 | 178.46 ± 0.01 | 178.48 ± 0.01 | 178.38 ± 0.02 | 177.86 ± 0.02 | 177.87 ± 0.01 |
| V97 | 177.77 ± 0.02 | 178.00 ± 0.01 | 177.99 ± 0.01 | 178.13 ± 0.01 | 178.01 ± 0.02 | 178.00 ± 0.02 |
| A98 | 179.77 ± 0.01 | 179.62 ± 0.01 | 179.55 ± 0.01 | 180.00 ± 0.02 | 179.98 ± 0.02 | 180.06 ± 0.02 |
| G99 | 176.08 ± 0.01 | 175.77 ± 0.01 | 175.82 ± 0.02 | 175.66 ± 0.03 | 175.03 ± 0.02 | 175.15 ± 0.03 |
| S102 | 176.38 ± 0.01 | 176.31 ± 0.01 | 176.28 ± 0.01 | 176.02 ± 0.01 | 175.87 ± 0.01 | 175.94 ± 0.01 |
| V106 | 178.06 ± 0.01 | 178.06 ± 0.01 | 178.01 ± 0.01 | 177.87 ± 0.02 | 177.95 ± 0.02 | 177.99 ± 0.02 |
| T107 | 176.18 ± 0.01 | 176.36 ± 0.01 | 176.26 ± 0.01 | 176.06 ± 0.02 | 176.04 ± 0.01 | 175.96 ± 0.01 |
| A108 | 179.13 ± 0.01 | 179.15 ± 0.01 | 179.22 ± 0.01 | 179.62 ± 0.02 | 179.23 ± 0.01 | 179.34 ± 0.02 |
| A109 | 179.71 ± 0.01 | 179.52 ± 0.01 | 179.15 ± 0.02 | 179.58 ± 0.02 | 179.36 ± 0.01 | 179.40 ± 0.02 |
| L110 | 179.15 ± 0.01 | 178.96 ± 0.01 | 178.67 ± 0.01 | 179.44 ± 0.02 | 179.22 ± 0.02 | 178.77 ± 0.02 |

TABLE S10: Simulated chemical shifts for N for the two chemical shift prediction methods and the three channel states. The data represented corresponds to the credible intervals of the chemical shifts that are inferred from the simulation data ($CS^{X}_{sim}$). The credible intervals are expressed as the center of the interval plus or minus the distance to the upper and lower bounds of the interval. This interval measures our uncertainty of the resonance position in the spectrum and not the peak width which is related to the standard deviation of the skew-normal distribution with which the data is modeled.

| Residue | Fully Open, SPARTA+ | Partially Open, SPARTA+ | Closed, SPARTA+ | Fully Open, SHIFTX2 | Partially Open, SHIFTX2 | Closed, SHIFTX2 |
|---|---|---|---|---|---|---|
| I38 | 120.04 ± 0.02 | 119.88 ± 0.03 | 119.88 ± 0.03 | 119.50 ± 0.04 | 119.38 ± 0.05 | 119.48 ± 0.04 |
| A42 | 121.09 ± 0.02 | 121.15 ± 0.03 | 121.10 ± 0.02 | 121.41 ± 0.05 | 121.49 ± 0.05 | 121.52 ± 0.05 |
| G43 | 106.46 ± 0.03 | 106.47 ± 0.03 | 106.32 ± 0.03 | 105.87 ± 0.05 | 105.82 ± 0.06 | 105.84 ± 0.06 |
| S44 | 119.13 ± 0.02 | 119.13 ± 0.02 | 118.93 ± 0.02 | 116.45 ± 0.04 | 116.29 ± 0.04 | 116.64 ± 0.04 |
| Y45 | 123.91 ± 0.04 | 123.97 ± 0.04 | 123.31 ± 0.03 | 121.70 ± 0.04 | 121.69 ± 0.04 | 121.38 ± 0.04 |
| A47 | 122.15 ± 0.03 | 122.30 ± 0.03 | 122.29 ± 0.03 | 121.14 ± 0.05 | 121.28 ± 0.05 | 121.32 ± 0.05 |
| V48 | 119.05 ± 0.03 | 119.07 ± 0.03 | 119.49 ± 0.02 | 117.27 ± 0.05 | 117.30 ± 0.05 | 117.55 ± 0.04 |
| A50 | 120.53 ± 0.04 | 120.74 ± 0.03 | 120.56 ± 0.03 | 119.94 ± 0.05 | 120.21 ± 0.05 | 120.44 ± 0.05 |
| E51 | 116.49 ± 0.06 | 116.49 ± 0.04 | 117.13 ± 0.04 | 115.80 ± 0.09 | 115.50 ± 0.07 | 116.40 ± 0.09 |
| G53 | 109.13 ± 0.08 | 109.00 ± 0.07 | 108.18 ± 0.07 | 110.11 ± 0.13 | 109.94 ± 0.14 | 108.72 ± 0.14 |
| G56 | 110.07 ± 0.08 | 110.28 ± 0.08 | 110.50 ± 0.07 | 109.65 ± 0.12 | 110.13 ± 0.11 | 110.48 ± 0.09 |
| A57 | 122.54 ± 0.05 | 122.16 ± 0.05 | 122.11 ± 0.04 | 122.66 ± 0.07 | 122.90 ± 0.05 | 122.89 ± 0.05 |
| Q58 | 119.73 ± 0.06 | 120.33 ± 0.07 | 120.43 ± 0.06 | 119.11 ± 0.09 | 119.74 ± 0.09 | 120.12 ± 0.09 |
| L59 | 118.56 ± 0.09 | 120.09 ± 0.08 | 120.40 ± 0.07 | 117.30 ± 0.10 | 119.18 ± 0.12 | 119.59 ± 0.09 |
| I60 | 114.79 ± 0.11 | 117.17 ± 0.20 | 117.76 ± 0.18 | 115.24 ± 0.14 | 117.91 ± 0.25 | 119.20 ± 0.26 |
| T61 | 111.83 ± 0.07 | 113.05 ± 0.09 | 111.85 ± 0.06 | 112.03 ± 0.11 | 113.62 ± 0.17 | 111.73 ± 0.10 |
| Y62 | 124.60 ± 0.08 | 125.25 ± 0.09 | 123.96 ± 0.07 | 123.68 ± 0.08 | 124.70 ± 0.09 | 124.00 ± 0.09 |
| R64 | 118.64 ± 0.03 | 118.42 ± 0.03 | 118.70 ± 0.03 | 118.87 ± 0.05 | 118.74 ± 0.06 | 119.08 ± 0.06 |
| A65 | 122.76 ± 0.02 | 122.85 ± 0.02 | 122.81 ± 0.02 | 121.75 ± 0.05 | 121.82 ± 0.05 | 121.78 ± 0.05 |
| L66 | 120.96 ± 0.02 | 120.90 ± 0.03 | 120.85 ± 0.03 | 120.04 ± 0.04 | 120.04 ± 0.05 | 120.07 ± 0.05 |
| W67 | 120.87 ± 0.02 | 120.66 ± 0.03 | 120.78 ± 0.03 | 120.62 ± 0.04 | 120.62 ± 0.04 | 120.67 ± 0.04 |
| W68 | 121.41 ± 0.02 | 121.56 ± 0.02 | 121.85 ± 0.03 | 119.78 ± 0.05 | 119.62 ± 0.04 | 119.56 ± 0.04 |
| S69 | 115.53 ± 0.02 | 115.51 ± 0.02 | 115.53 ± 0.02 | 115.34 ± 0.04 | 115.42 ± 0.04 | 115.10 ± 0.05 |
| V70 | 121.42 ± 0.03 | 121.44 ± 0.04 | 121.87 ± 0.05 | 120.28 ± 0.05 | 120.00 ± 0.07 | 120.55 ± 0.06 |
| E71 | 119.07 ± 0.04 | 118.95 ± 0.05 | 118.77 ± 0.04 | 118.47 ± 0.07 | 118.47 ± 0.09 | 118.29 ± 0.09 |

| Residue | Fully Open, SPARTA+ | Partially Open, SPARTA+ | Closed, SPARTA+ | Fully Open, SHIFTX2 | Partially Open, SHIFTX2 | Closed, SHIFTX2 |
|---|---|---|---|---|---|---|
| T72 | 115.95 ± 0.04 | 116.39 ± 0.04 | 116.71 ± 0.04 | 115.35 ± 0.08 | 115.83 ± 0.08 | 115.95 ± 0.07 |
| A73 | 122.82 ± 0.03 | 122.89 ± 0.03 | 122.82 ± 0.03 | 122.91 ± 0.08 | 123.09 ± 0.07 | 122.92 ± 0.07 |
| T74 | 108.09 ± 0.05 | 108.18 ± 0.05 | 108.79 ± 0.05 | 106.89 ± 0.09 | 107.41 ± 0.11 | 107.34 ± 0.10 |
| T75 | 110.40 ± 0.04 | 110.36 ± 0.04 | 110.28 ± 0.04 | 114.93 ± 0.03 | 114.72 ± 0.03 | 114.83 ± 0.03 |
| V76 | 120.81 ± 0.05 | 121.13 ± 0.04 | 121.16 ± 0.05 | 120.66 ± 0.04 | 120.73 ± 0.03 | 120.79 ± 0.03 |
| G77 | 106.44 ± 0.03 | 106.42 ± 0.02 | 106.42 ± 0.02 | 108.85 ± 0.04 | 108.83 ± 0.04 | 108.73 ± 0.03 |
| Y78 | 121.98 ± 0.06 | 121.96 ± 0.06 | 121.99 ± 0.06 | 119.92 ± 0.10 | 120.53 ± 0.10 | 120.39 ± 0.11 |
| G79 | 102.76 ± 0.04 | 102.63 ± 0.03 | 102.77 ± 0.03 | 106.09 ± 0.03 | 106.15 ± 0.03 | 106.27 ± 0.03 |
| D80 | 119.19 ± 0.03 | 119.19 ± 0.03 | 119.09 ± 0.03 | 119.27 ± 0.04 | 119.43 ± 0.04 | 119.41 ± 0.04 |
| L81 | 118.27 ± 0.10 | 118.35 ± 0.08 | 118.52 ± 0.07 | 117.99 ± 0.07 | 118.07 ± 0.07 | 118.09 ± 0.06 |
| Y82 | 116.64 ± 0.08 | 116.96 ± 0.08 | 117.23 ± 0.08 | 118.12 ± 0.10 | 118.08 ± 0.10 | 118.35 ± 0.10 |
| V84 | 117.15 ± 0.15 | 117.43 ± 0.12 | 117.69 ± 0.12 | 119.33 ± 0.13 | 119.08 ± 0.11 | 119.16 ± 0.11 |
| T85 | 113.79 ± 0.06 | 113.74 ± 0.05 | 114.20 ± 0.05 | 111.26 ± 0.12 | 111.18 ± 0.11 | 111.73 ± 0.10 |
| L86 | 123.75 ± 0.07 | 123.65 ± 0.05 | 124.07 ± 0.05 | 124.13 ± 0.07 | 123.95 ± 0.07 | 124.37 ± 0.06 |
| W87 | 117.81 ± 0.03 | 117.71 ± 0.03 | 117.57 ± 0.03 | 117.41 ± 0.05 | 117.33 ± 0.06 | 117.30 ± 0.05 |
| G88 | 106.96 ± 0.03 | 107.25 ± 0.03 | 107.32 ± 0.04 | 106.76 ± 0.07 | 106.98 ± 0.06 | 107.07 ± 0.06 |
| R89 | 122.13 ± 0.02 | 122.21 ± 0.02 | 122.20 ± 0.02 | 120.42 ± 0.04 | 120.72 ± 0.03 | 120.73 ± 0.03 |
| L90 | 120.25 ± 0.03 | 120.37 ± 0.03 | 120.34 ± 0.03 | 119.40 ± 0.05 | 119.46 ± 0.05 | 119.41 ± 0.05 |
| V91 | 119.11 ± 0.02 | 118.93 ± 0.03 | 118.95 ± 0.03 | 119.03 ± 0.04 | 118.83 ± 0.05 | 118.61 ± 0.07 |
| A92 | 122.81 ± 0.03 | 122.79 ± 0.02 | 122.73 ± 0.03 | 121.10 ± 0.05 | 121.05 ± 0.05 | 121.08 ± 0.06 |
| V93 | 118.22 ± 0.05 | 118.84 ± 0.03 | 117.95 ± 0.06 | 117.34 ± 0.05 | 117.66 ± 0.04 | 117.36 ± 0.06 |
| V94 | 119.72 ± 0.04 | 119.46 ± 0.02 | 119.77 ± 0.03 | 119.23 ± 0.07 | 119.34 ± 0.05 | 119.41 ± 0.04 |
| V95 | 120.11 ± 0.03 | 119.85 ± 0.02 | 119.81 ± 0.02 | 119.48 ± 0.05 | 119.58 ± 0.04 | 119.55 ± 0.04 |
| M96 | 119.27 ± 0.02 | 118.11 ± 0.04 | 118.22 ± 0.04 | 118.11 ± 0.03 | 117.42 ± 0.06 | 117.53 ± 0.05 |
| V97 | 119.45 ± 0.02 | 120.45 ± 0.04 | 120.11 ± 0.05 | 118.14 ± 0.05 | 119.21 ± 0.06 | 119.06 ± 0.07 |
| A98 | 121.15 ± 0.03 | 121.98 ± 0.03 | 122.40 ± 0.03 | 121.01 ± 0.06 | 121.44 ± 0.05 | 121.79 ± 0.05 |
| G99 | 107.39 ± 0.03 | 107.91 ± 0.02 | 107.51 ± 0.02 | 104.73 ± 0.08 | 105.54 ± 0.05 | 105.28 ± 0.04 |
| S102 | 118.34 ± 0.02 | 118.80 ± 0.02 | 118.54 ± 0.03 | 115.28 ± 0.07 | 116.46 ± 0.05 | 116.55 ± 0.04 |

| Residue | Fully Open, SPARTA+ | Partially Open, SPARTA+ | Closed, SPARTA+ | Fully Open, SHIFTX2 | Partially Open, SHIFTX2 | Closed, SHIFTX2 |
|---|---|---|---|---|---|---|
| F103 | 123.52 ± 0.03 | 121.27 ± 0.04 | 122.08 ± 0.07 | 121.95 ± 0.05 | 120.11 ± 0.07 | 120.72 ± 0.08 |
| V106 | 118.68 ± 0.04 | 118.22 ± 0.06 | 118.87 ± 0.03 | 118.51 ± 0.06 | 117.92 ± 0.09 | 118.61 ± 0.05 |
| T107 | 113.80 ± 0.05 | 115.97 ± 0.06 | 116.68 ± 0.02 | 112.95 ± 0.11 | 115.22 ± 0.12 | 116.47 ± 0.05 |
| A108 | 123.63 ± 0.02 | 123.83 ± 0.03 | 123.81 ± 0.02 | 123.47 ± 0.06 | 122.67 ± 0.07 | 122.26 ± 0.05 |
| A109 | 121.81 ± 0.02 | 121.99 ± 0.02 | 121.73 ± 0.02 | 120.50 ± 0.04 | 120.58 ± 0.04 | 120.79 ± 0.04 |
| L110 | 119.47 ± 0.02 | 119.89 ± 0.02 | 119.48 ± 0.02 | 118.66 ± 0.04 | 118.94 ± 0.04 | 118.86 ± 0.04 |

TABLE S11: Difference between simulated and experimental chemical shifts for $C_\alpha$ for the two chemical shift prediction methods and the Fully Open and Fully Partially states ($CS_{sim}^{FO}$,$CS_{sim}^{PO}$). The last 4 columns show the relative chemical shifts using the closed state as a reference.

| | $CS_{sim}^{X} - CS_{exp}^{pH = 4, act}$ | | | | $\Delta\Delta CS_{sim}^{X}$ | | | |
|---|---|---|---|---|---|---|---|---|
| Residue | Fully Open, SPARTA+ | Partially Open, SPARTA+ | Fully Open, SHIFTX2 | Partially Open, SHIFTX2 | Fully Open, SPARTA+ | Partially Open, SPARTA+ | Fully Open, SHIFTX2 | Partially Open, SHIFTX2 |
| A50 | 0.5±0.0 | 0.8±0.0 | | | -0.17±0.02 | 0.11±0.02 | | |
| G56 | | | 0.5±0.0 | 0.8±0.0 | | | -0.45±0.02 | -0.21±0.02 |
| Q58 | 4.3±0.0 | 3.6±0.0 | | | 0.77±0.05 | 0.12±0.05 | | |
| T61 | 1.7±0.0 | 2.5±0.0 | 1.6±0.0 | 2.5±0.0 | -0.04±0.03 | 0.81±0.05 | 0.11±0.03 | 1.03±0.05 |
| R64 | 0.0±0.0 | -0.4±0.0 | 0.3±0.0 | -0.2±0.0 | 0.26±0.02 | -0.12±0.03 | 0.21±0.03 | -0.29±0.04 |
| T75 | 1.0±0.0 | 0.7±0.0 | | | 0.26±0.02 | -0.08±0.02 | | |
| Y78 | -1.6±0.0 | -0.9±0.0 | -1.1±0.0 | -0.3±0.0 | -0.65±0.04 | 0.03±0.03 | -0.69±0.03 | 0.03±0.02 |
| R89 | 0.1±0.0 | 0.4±0.0 | 0.5±0.0 | 0.8±0.0 | -0.07±0.02 | 0.15±0.02 | -0.18±0.03 | 0.15±0.02 |
| V93 | -0.3±0.0 | 0.2±0.0 | | | -0.37±0.04 | 0.12±0.04 | | |
| M96 | 0.4±0.0 | -0.2±0.0 | -0.1±0.0 | -0.4±0.0 | 0.59±0.02 | -0.11±0.02 | 0.20±0.02 | -0.14±0.02 |
| A98 | 0.5±0.0 | 0.2±0.0 | | | 0.08±0.02 | -0.15±0.02 | | |
| G99 | | | -0.2±0.0 | 0.1±0.0 | | | -0.08±0.02 | 0.21±0.02 |
| S102 | -1.2±0.0 | -0.8±0.0 | -1.0±0.0 | -0.7±0.0 | -0.55±0.02 | -0.11±0.02 | -0.52±0.03 | -0.16±0.02 |
| T107 | -2.1±0.0 | -0.8±0.0 | -1.8±0.0 | -0.9±0.0 | -1.06±0.02 | 0.21±0.02 | -0.86±0.04 | -0.02±0.03 |

TABLE S12: Difference between simulated and experimental chemical shifts for $C_\beta$ for the two chemical shift prediction methods and the Partially Open and Fully Open states ($CS_{sim}^{FO} - CS_{sim}^{PO}$). The last 4 columns show the relative chemical shifts using the closed state as a reference.

| | $CS_{sim}^{X} - CS_{exp}^{pH = 4, act}$ | | | | $\Delta\Delta CS_{sim}^{X}$ | | | |
|---|---|---|---|---|---|---|---|---|
| **Residue** | **Fully Open, SPARTA+** | **Partially Open, SPARTA+** | **Fully Open, SHIFTX2** | **Partially Open, SHIFTX2** | **Fully Open, SPARTA+** | **Partially Open, SPARTA+** | **Fully Open, SHIFTX2** | **Partially Open, SHIFTX2** |
| I60 | -0.3±0.0 | 0.7±0.1 | 0.1±0.0 | 0.9±0.0 | -0.88±0.06 | 0.12±0.08 | -0.79±0.04 | 0.05±0.06 |
| T61 | 0.0±0.0 | -0.9±0.0 | -0.2±0.0 | -0.7±0.0 | 0.17±0.03 | -0.74±0.05 | -0.01±0.03 | -0.52±0.04 |
| Y62 | -0.4±0.0 | 0.1±0.0 | | | -0.38±0.02 | 0.12±0.02 | | |
| R64 | | | -0.9±0.0 | -1.2±0.0 | | | 0.19±0.01 | -0.07±0.02 |
| Y78 | | | 0.4±0.0 | 1.0±0.0 | | | -0.57±0.02 | 0.01±0.01 |
| M96 | -0.9±0.0 | -1.1±0.0 | -0.3±0.0 | -0.8±0.0 | 0.04±0.01 | -0.17±0.01 | 0.28±0.02 | -0.18±0.02 |
| V97 | 0.4±0.0 | 0.1±0.0 | | | 0.45±0.02 | 0.17±0.02 | | |
| A98 | | | 0.2±0.0 | -0.2±0.0 | | | -0.11±0.02 | -0.52±0.01 |
| S102 | | | 0.8±0.0 | 0.6±0.0 | | | 0.17±0.02 | -0.04±0.02 |
| T107 | 1.0±0.0 | 1.3±0.0 | | | -0.35±0.01 | -0.10±0.01 | | |
| A109 | 1.3±0.0 | 1.5±0.0 | | | 1.09±0.02 | 1.29±0.02 | | |

TABLE S13: Difference between simulated and experimental chemical shifts for C for the two chemical shift prediction methods and the Partially Open and Fully Open states ($CS_{sim}^{FO}$,$CS_{sim}^{PO}$). The last 4 columns show the relative chemical shifts using the closed state as a reference.

| | $CS_{sim}^{X} - CS_{exp}^{pH = 4, act}$ | | | | $\Delta\Delta CS_{sim}^{X}$ | | | |
|---|---|---|---|---|---|---|---|---|
| **Residue** | **Fully Open, SPARTA+** | **Partially Open, SPARTA+** | **Fully Open, SHIFTX2** | **Partially Open, SHIFTX2** | **Fully Open, SPARTA+** | **Partially Open, SPARTA+** | **Fully Open, SHIFTX2** | **Partially Open, SHIFTX2** |
| A50 | 0.3±0.0 | 0.7±0.0 | -0.1±0.0 | 0.5±0.0 | -0.41±0.03 | -0.03±0.03 | -0.73±0.04 | -0.13±0.03 |
| E51 | 0.4±0.0 | 1.2±0.0 | | | -1.25±0.05 | -0.50±0.06 | | |
| G56 | 0.3±0.0 | 0.6±0.0 | | | -0.32±0.03 | -0.04±0.02 | | |
| Q58 | -1.7±0.0 | -2.1±0.0 | | | 0.76±0.04 | 0.38±0.04 | | |

| Residue | Fully Open, SPARTA+ | Partially Open, SPARTA+ | Fully Open, SHIFTX2 | Partially Open, SHIFTX2 | Fully Open, SPARTA+ | Partially Open, SPARTA+ | Fully Open, SHIFTX2 | Partially Open, SHIFTX2 |
|---|---|---|---|---|---|---|---|---|
| L59 | 2.7±0.0 | 2.1±0.0 | | | 0.88±0.04 | 0.27±0.05 | | |
| M96 | 2.1±0.0 | 1.6±0.0 | 1.5±0.0 | 1.0±0.0 | 0.28±0.01 | -0.18±0.02 | 0.35±0.02 | -0.17±0.02 |
| V97 | -0.7±0.0 | -0.4±0.0 | | | -0.18±0.02 | 0.05±0.02 | | |
| G99 | 1.8±0.0 | 1.4±0.0 | 1.3±0.0 | 0.7±0.0 | 0.02±0.02 | -0.29±0.02 | 0.27±0.04 | -0.35±0.03 |
| A108 | | | 0.1±0.0 | -0.3±0.0 | | | -1.70±0.03 | -2.08±0.02 |
| A109 | | | 1.9±0.0 | 1.6±0.0 | | | 0.54±0.02 | 0.32±0.02 |

TABLE S14: Difference between simulated and experimental chemical shifts for N for the two chemical shift prediction methods and the Partially Open and Fully Open states ($CS^{FO}_{sim}$,$CS^{PO}_{sim}$). The last 4 columns show the relative chemical shifts using the closed state as a reference.

| | $CS^{X}_{sim}-CS^{pH = 4, act}_{exp}$ | | | | $\Delta\Delta CS^{X}_{sim}$ | | | |
|---|---|---|---|---|---|---|---|---|
| Residue | Fully Open, SPARTA+ | Partially Open, SPARTA+ | Fully Open, SHIFTX2 | Partially Open, SHIFTX2 | Fully Open, SPARTA+ | Partially Open, SPARTA+ | Fully Open, SHIFTX2 | Partially Open, SHIFTX2 |
| L59 | -7.0±0.1 | -5.4±0.1 | -8.2±0.1 | -6.4±0.1 | -2.07±0.11 | -0.54±0.11 | -2.53±0.14 | -0.65±0.15 |
| I60 | | | 10.1±0.1 | 12.8±0.3 | | | -3.81±0.30 | -1.17±0.37 |
| T61 | | | 2.4±0.1 | 4.0±0.2 | | | 0.09±0.15 | 1.68±0.20 |
| Y62 | | | 0.2±0.1 | 1.2±0.1 | | | -0.29±0.12 | 0.74±0.12 |
| V93 | 1.2±0.0 | 1.8±0.0 | | | 0.51±0.08 | 1.12±0.06 | | |
| M96 | 2.9±0.0 | 1.7±0.0 | 1.7±0.0 | 1.0±0.1 | 0.73±0.04 | -0.42±0.05 | 0.26±0.06 | -0.44±0.08 |
| V97 | 3.4±0.0 | 4.4±0.0 | 2.1±0.1 | 3.2±0.1 | -0.76±0.05 | 0.23±0.06 | -1.03±0.09 | 0.04±0.10 |
| A98 | -3.6±0.0 | -2.8±0.0 | | | -1.18±0.04 | -0.34±0.04 | | |
| G99 | 0.5±0.0 | 1.1±0.0 | -2.1±0.1 | -1.3±0.1 | 0.06±0.04 | 0.59±0.03 | -0.38±0.09 | 0.44±0.07 |
| S102 | | | -0.5±0.1 | 0.7±0.0 | | | -0.40±0.08 | 0.79±0.06 |
| F103 | 3.3±0.0 | 1.1±0.0 | 1.7±0.1 | -0.1±0.1 | 1.50±0.08 | -0.76±0.08 | 1.27±0.09 | -0.57±0.10 |
| T107 | -2.5±0.0 | -0.3±0.1 | -3.3±0.1 | -1.1±0.1 | -2.01±0.05 | 0.17±0.07 | -2.63±0.12 | -0.38±0.13 |
| A108 | | | 1.9±0.1 | 1.1±0.1 | | | 0.72±0.08 | -0.08±0.09 |
